# Estimating the genome-wide contribution of selection to temporal allele frequency change

**DOI:** 10.1101/798595

**Authors:** Vince Buffalo, Graham Coop

**Affiliations:** Population Biology Graduate Group; Center for Population Biology, Department of Evolution and Ecology, University of California, Davis, CA 95616

## Abstract

Rapid phenotypic adaptation is often observed in natural populations and selection experiments. However, detecting the genome-wide impact of this selection is difficult, since adaptation often proceeds from standing variation and selection on polygenic traits, both of which may leave faint genomic signals indistinguishable from a noisy background of genetic drift. One promising signal comes from the genome-wide covariance between allele frequency changes observable from temporal genomic data, e.g. evolve-and-resequence studies. These temporal covariances reflect how heritable fitness variation in the population leads changes in allele frequencies at one timepoint to be predictive of the changes at later timepoints, as alleles are indirectly selected due to remaining associations with selected alleles. Since genetic drift does not lead to temporal covariance, we can use these covariances to estimate what fraction of the variation in allele frequency change through time is driven by linked selection. Here, we reanalyze three selection experiments to quantify the effects of linked selection over short timescales using covariance among time-points and across replicates. We estimate that at least 17% to 37% of allele frequency change is driven by selection in these experiments. Against this background of positive genome-wide temporal covariances we also identify signals of negative temporal covariance corresponding to reversals in the direction of selection for a reasonable proportion of loci over the time course of a selection experiment. Overall, we find that in the three studies we analyzed, linked selection has a large impact on short-term allele frequency dynamics that is readily distinguishable from genetic drift.

**Significance Statement:** A long-standing problem in evolutionary biology is to understand the processes that shape the genetic composition of populations. In a population without migration, the two processes that change allele frequencies are selection, which increases beneficial alleles and removes deleterious ones, and genetic drift which randomly changes frequencies as some parents contribute more or less alleles to the next generation. Previous efforts to disentangle these processes have used genomic samples from a single timepoint and models of how selection affects neighboring sites (linked selection). Here, we use genomic data taken through time to quantify the contributions of selection and drift to genome-wide frequency changes. We show selection acts over short timescales in three evolve-and-resequence studies and has a sizable genome-wide impact.

## 1 Introduction

A long-standing problem in evolutionary genetics is quantifying the roles of genetic drift and selection in shaping genome-wide allele frequency changes. Selection can affect allele frequencies, both directly and indirectly, with the indirect effect coming from the action of selection on correlated loci elsewhere in genome, e.g. linked selection (Charlesworth et al. 1993; Maynard Smith and Haigh 1974; Nordborg et al. 1996; see Barton 2000 for a review). Previous work has mostly focused on teasing apart the impacts of drift and selection on genome-wide diversity using population samples from a single contemporary timepoint, often by modeling the correlation between regional recombination rate, gene density, and diversity created in the presence of linked selection (Cutter and Payseur 2013; Sella et al. 2009). This approach has shown linked selection has a major role in shaping patterns of genome-wide diversity across the genomes of a range of sexual species (Andersen et al. 2012; Andolfatto 2007; Begun et al. 2007; Beissinger et al. 2016; Cutter and Choi 2010; Elyashiv et al. 2016; Macpherson et al. 2007; Sattath et al. 2011; Williamson et al. 2014), and has allowed us to quantify the relative influence of positive selection (hitchhiking) and negative selection (background selection; Andolfatto 2007; Elyashiv et al. 2016; Hernandez et al. 2011; Macpherson et al. 2007; McVicker et al. 2009; Nordborg et al. 2005). However, we lack an understanding of both how linked selection acts over short time intervals and of its full impact on genome-wide allele frequency changes.

There are numerous examples of rapid phenotypic adaptation (Franks et al. 2007; Grant and Grant 2006, 2011; Reznick et al. 1997) and rapid, selection-driven genomic evolution in asexual populations (Baym et al. 2016; Bennett et al. 1990; Good et al. 2017). Yet the polygenic nature of fitness makes detecting the impact of selection on genome-wide variation over short timescales in sexual populations remarkably difficult (Kemper et al. 2014; Latta 1998; Pritchard et al. 2010). This is because the effect of selection on a polygenic trait (such as fitness) is distributed across numerous loci. This can lead to subtle allele frequency shifts on standing variation that are difficult to distinguish from background levels of genetic drift and sampling variance. Increasingly, genomic experimental evolution studies with multiple timepoints, and in some cases multiple replicate populations, are being used to detect large-effect selected loci (Turner and Miller 2012; Turner et al. 2011) and differentiate modes of selection (Barghi et al. 2019; Burke et al. 2010; Therkildsen et al. 2019). In addition these temporal-genomic studies have begun in wild populations, some with the goal of finding variants that exhibit frequency changes consistent with fluctuating selection (Bergland et al. 2014; Machado et al. 2018). In a previous paper, we proposed that one useful signal for understanding the genome-wide impact of polygenic linked selection detectable from temporal genomic data is the temporal autocovariance (i.e. covariance between two timepoints) of allele frequency changes (Buffalo and Coop 2019). These covariances are created when the loci that underly heritable fitness variation perturb the frequencies of linked alleles; in contrast, when genetic drift acts alone in a closed population, these covariances are expected to be zero for neutral alleles. Mathematically, temporal covariances are useful because it is natural to decompose the total variance in allele frequency change across a time interval into the variances and covariances in allele frequency change between generations. Furthermore, biologically, these covariances reflect the extent to which allele frequency changes in one generation predict changes in another due to a shared selection pressures and associations to selected loci.

Here, we provide the first empirical analyses to quantify the impact of linked selection acting over short timescales (tens of generations) across two evolve and re-sequence studies (Barghi et al. 2019; Kelly and Hughes 2019), and an artificial selection experiment (Castro et al. 2019). These sequencing selection experiments have started to uncover selected loci contributing to the adaptive response; however it is as yet far from clear how much of genome-wide allele frequency changes are driven by selection or genetic drift. We repeatedly find a signal of temporal covariance, consistent with linked selection acting to significantly perturb genome-wide allele frequency changes across the genome in a manner that other approaches would not be able differentiate from genetic drift. We estimate a lower bound of the fraction of variance in allele frequency change caused by selection, as well as the correlation between allele frequency changes between replicate populations caused by convergent selection pressures. Overall, we demonstrate that linked selection has a powerful role in shaping genome-wide allele frequency changes over very short timescales in experimental evolution.

## Results

We first analyzed the dataset of Barghi et al. (2019), an evolve-and-resequence study with ten replicate populations exposed to a high temperature lab environment and evolved for 60 generations, and sequenced every ten generations. Using the seven timepoints and ten replicate populations, we estimated the genome-wide 6 × 6 temporal covariance matrix **Q** for each of the ten replicates. Each row of these matrices represent the temporal covariance Cov(Δ_10_ *p*_*s*_, Δ_10_ *p*_*t*_), between the allele frequency change (in ten-generation intervals, denoted Δ_10_ *p*_*t*_) of some initial reference generation *s* (the row of the matrix), and some later timepoint *t* (the column of the matrix). We corrected these matrices for biases created due to sampling noise, and normalized the entries for heterozygosity (see SI Appendix, sections S1.2 and S1.4). These covariances are expected to be zero when only drift is acting, as only heritable variation for fitness can create covariance between allele frequency changes in a closed population (Buffalo and Coop 2019). Averaging across the ten replicate temporal covariances matrices, we find temporal covariances that are statistically significant (95% block bootstraps CIs do not contain zero), consistent with linked selection perturbing genome-wide allele frequency changes over very short time periods. The covariances between all adjacent time intervals are positive and then decay towards zero as we look at more distant time intervals (Figure 1 A), as expected when directional selection affects linked variants’ frequency trajectories until ultimately linkage disequilibrium and the associated additive genetic variance for fitness decays (which could occur as a population reaches a new optimum, and directional selection weakens) (Buffalo and Coop 2019). The temporal covariances per replicate are noisier but this general pattern holds; see SI Appendix, Fig. S23.

**Figure 1:**
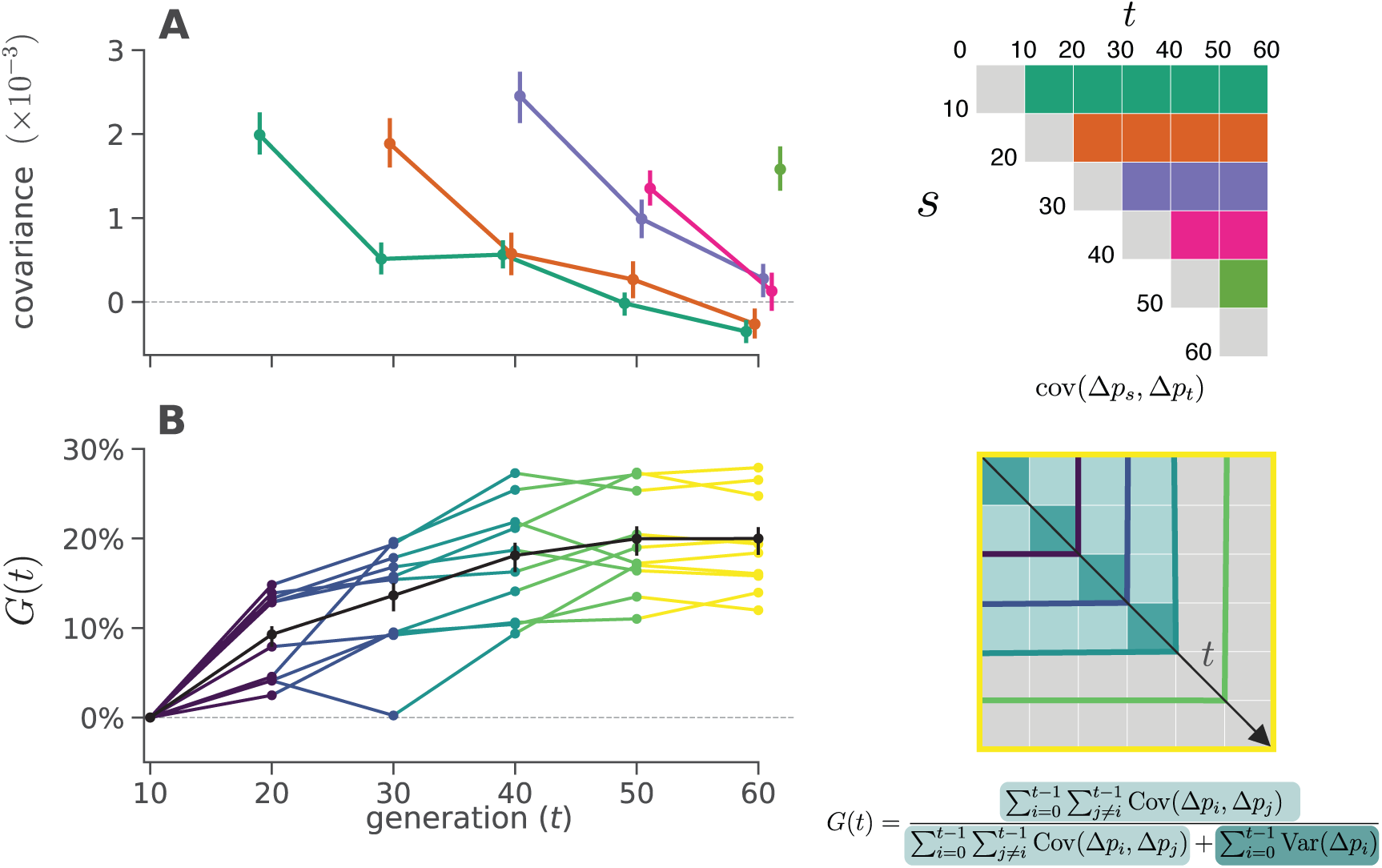
**A**: Temporal covariance, averaged across all ten replicate populations, through time from the Barghi et al. (2019) study. Each line depicts the temporal covariance Cov(Δ*p*_*s*_, Δ*p*_*t*_) from some reference generation *s* to a later time *t* which varies along the x-axis; each line corresponds to a row of the upper-triangle of the temporal covariance matrix with the same color (upper right). The ranges around each point are 95% block-bootstrap confidence intervals. **B**: A lower bound on the proportion of the total variance in allele frequency change explained by linked selection, *G*(*t*), as it varies through time *t* along the x-axis. The black line is the *G*(*t*) averaged across replicates, with the 95% block-bootstrap confidence interval. The other lines are the *G*(*t*) for each individual replicate, with colors indicating what subset of the temporal-covariance matrix to the right is being included in the calculation of *G*(*t*).

Since our covariances are averages over loci, the covariance estimate could be strongly affected by a few outlier regions. To test whether large outlier regions drive the genome-wide signal we see in the Barghi et al. (2019) data, we calculate the covariances in 100kb windows along the genome (we refer to these as windowed covariances throughout) and take the median windowed covariance, and trimmed-mean windowed covariance, as a measure of the genome-wide covariance robust to large-effect loci. These robust estimates (SI Appendix, Table S1 and SI Appendix, Fig. S24) confirm the patterns we see using the mean covariance, establishing that genomic temporal covariances are non-zero due to the impact of selection acting across many genomic regions.

While the presence of positive temporal covariances is consistent with selection affecting allele frequencies over time, this measure is not easily interpretable. We can calculate a more intuitive measure from the temporal covariances to quantify the impact of selection on allele frequency change: the ratio of total covariance in allele frequency change to the total variance in allele frequency change. We denote the change in allele frequency as Δ*p*_*t*_ = *p*_*t*+1_ − *p*_*t*_, where *p*_*t*_ is the allele frequency in generation *t*. Since the total variation in allele frequency change can be partitioned into variance and covariance components, 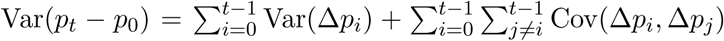 (we correct for biases due to sequencing depth), and the covariances are zero when drift acts alone, this is a lower bound on how much of the variance in allele frequency change is caused by linked selection (Buffalo and Coop 2019). We call this measure *G*(*t*), defined as

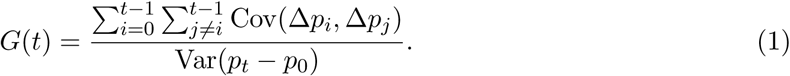

This estimates the impact of selection on allele frequency change between the initial generation 0 and some later generation *t*, which can be varied to see how this quantity grows through time. When the sum of the covariances is positive, this measure can intuitively be understood as a lower bound on relative fraction of allele frequency change normally thought of as “drift” that is actually due to selection. Additionally, *G*(*t*) can be understood as a short-timescale estimate of the reduction in neutral diversity due to linked selection (or equivalently the reduction in neutral effective population size needed to account linked selection, see SI Appendix, section S7). Since the Barghi et al. (2019) experiment is sequenced every ten generations, the numerator uses the covariances estimated between ten-generation blocks of allele frequency change; thus the strong, unobservable, covariances between adjacent generations do not contribute to the numerator of *G*(*t*). Had these covariances been measurable on shorter timescales, their cumulative effect would likely have been higher yet (see SI Appendix, sections S2 and S8.4 for more details). Additionally, selection inflates the variance in allele frequency change per generation; however, this effect cannot be easily distinguished from drift. For both these reasons, our measure *G*(*t*) is quite conservative (we demonstrate this through simulations in SI Appendix, section S8.4). Still, we find a remarkably strong signal. Greater than 20% of total, genome-wide allele frequency change over 60 generations is the result of selection (Figure 1 B). This proportion of variance attributable to selection builds over time in Figure 1B as the effects of linked selection are compounded over the generations unlike genetic drift. Our G(t) starts to plateau to a constant level as the covariances from earlier generations have decayed and so no longer contribute as strongly (Figure 1).

Additionally, we looked for a signal of temporal autocovariance in Bergland et al. (2014), a study that collected *Drosophila melanogaster* through Spring-Fall season pairs across three years. If there was a strong pattern of genome-wide fluctuating selection, we might expect a pattern of positive covariances between similar seasonal changes, e.g. Spring-Fall in two adjacent years, and negative covariances between dissimilar seasonal changes, e.g. Spring-Fall and Fall-Spring in two adjacent years. However, we find no such signal over years, and in reproducing their original analysis, we find that their number of statistically significant seasonal SNPs is not enriched compared to an empirical null distribution created by permuting seasonal labels; we discuss this in more depth in SI Appendix, section S6.

The replicate design of Barghi et al. (2019) allows us to quantify another covariance: the covariance in allele frequency change between replicate populations experiencing convergent selection pressures. These between-replicate covariances are created in the same way as temporal covariances: alleles linked to a particular fitness background are expected to have allele frequency changes in the same direction if the selection pressures are similar. Intuitively, where temporal covariances reflect that alleles associated with heritable fitness backgrounds are predictive of frequency changes between generations, replicate covariances reflect that heritable fitness backgrounds common to each replicate predict (under the same selection pressures) frequency changes between replicates; we note that there is not a direct one-to-one correspondence between temporal and replicate covariances, since the latter are driven by a shared selection pressure and the stochastic genetic backgrounds across replicate populations. We measure this through a statistic similar to a correlation, which we call the convergent correlation: the ratio of average between-replicate covariance across all pairs to the average standard deviation across all pairs of replicates,

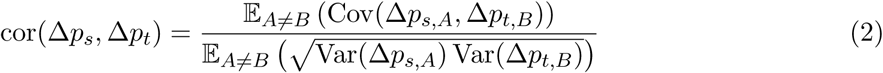

where *A* and *B* here are two replicate labels, and for the Barghi et al. (2019) data, we use Δ_10_ *p*_*t*_.

We have calculated the convergent correlation for all rows of the replicate covariance matrices. Like temporal covariances, we visualize these through time (Figure 2A), with each line representing the convergent correlation from a particular reference generation *s* as it varies with *t* (shown on the x-axis). In other words, each of the colored lines corresponds to the like-colored row of the convergence correlation matrix (upper left in Figure 2A). We find these convergent correlation coefficients are relatively weak, and decay very quickly from an initial value of about 0.1 (95% block bootstrap confidence intervals [0.094, 0.11]) to around 0.01 (95% CIs [0.0087, 0.015]) within 20 generations. This suggests that while a substantial fraction of the initial response is shared over the replicates, this is followed by a rapid decay, a result consistent with the primary finding of the original Barghi et al. (2019) study: that alternative loci contribute to longer term adaptation across the different replicates.

**Figure 2:**
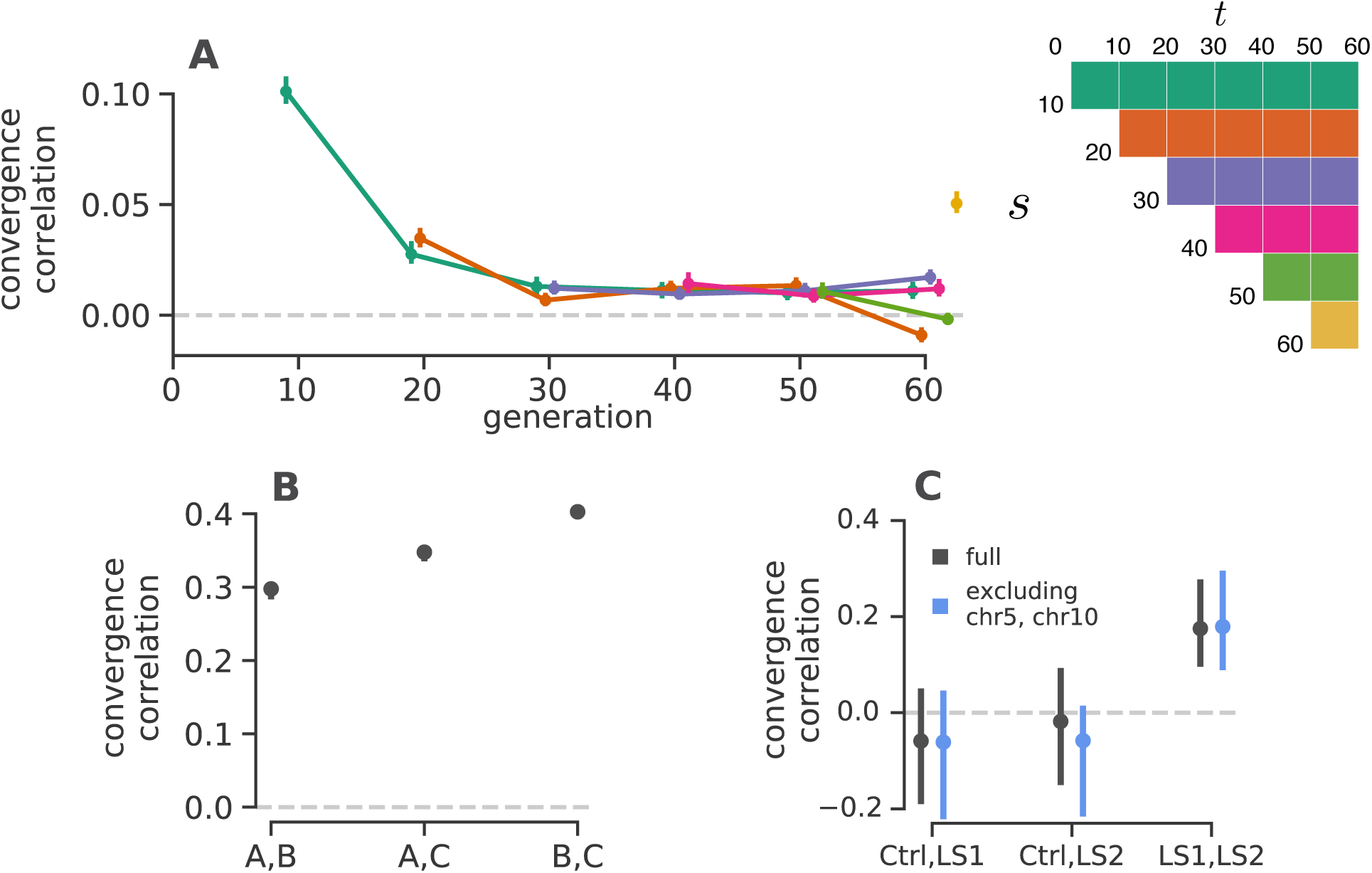
**A**: The convergence correlations, averaged across Barghi et al. (2019) replicate pairs, through time. Each line represents the convergence correlation cor(Δ*p*_*s*_, Δ*p*_*t*_) from a starting reference generation *s* to a later time *t*, which varies along the x-axis; each line corresponds to a row of the temporal convergence correlation matrix depicted to the right (where the diagonal elements represent the convergence correlations between the same timepoints across replicate populations). We note that convergent correlation for the last timepoint is an outlier; we are unsure as to the cause of this, e.g. it does not appear to be driven by a single pair of replicates. **B**: The convergence correlations between individual pairs of replicates in the Kelly and Hughes (2019) data (note the confidence intervals are plotted, but are small on this y-axis scale). **C**: The convergence correlations between individual pairs of replicates in Castro et al. (2019) data, for the two selection lines (LS1 and LS2) and the control (Ctrl); gray CIs are those using the complete dataset, blue CIs exclude chromosomes 5 and 10 which harbor the two regions Castro et al. (2019) found to have signals of parallel selection between LS1 and LS2. Through simulations, we have found that the differences in convergence correlation confidence interval widths between these *Drosophila* studies and the Longshanks study are due to the differing population sizes.

A benefit of between-replicate covariances is that unlike temporal covariances, these can be calculated with only two sequenced timepoints and a replicated study design. This allowed us to assess the impact of linked selection in driving convergent patterns of allele frequency change across replicate populations in two other studies. First, we reanalyzed the selection experiment of Kelly and Hughes (2019), which evolved three replicate wild populations of *Drosophila simulans* for 14 generations adapting to a novel laboratory environment. Since each replicate was exposed to the same selection pressure and share linkage disequilibria common to the original natural founding population, we expected each of the three replicate populations to have positive convergence correlations. We find all three convergent correlation coefficients between replicate pairs are significant (Figure 2B), and average to 0.36 (95% CI [0.31, 0.40]). Additionally, we can calculate the proportion of the total variance in allele frequency change from convergent selection pressure, analogous to our *G*(*t*), where the numerator is the convergent covariance and the denominator is the total variance (see SI appendix, section S4). We find that 37% of the total variance is due to shared allele frequency changes caused by selection (95% CI [29%, 41%]; these are similar to the convergence correlation, since the variance is relatively constant across the replicates.

Next, we reanalyzed the Longshanks selection experiment, which selected for longer tibiae length relative to body size in mice, leading to a response to selection of about 5 standard deviations over the course of twenty generations (Castro et al. 2019; Marchini et al. 2014). This study includes two independent selection lines, Longshanks 1 and 2 (LS1 and LS2), and an unselected control line (Ctrl) where parents were randomly selected. Consequently, this selection experiment offers a useful control to test our convergence correlations: we expect to see significant positive convergence correlations in the comparison between the two Longshanks selection lines, but not between each of the control line and Longshanks line pairs. We find that this is the case (gray confidence intervals in Figure 2C), with convergence correlations between each of the Longshanks lines to the control not being statistically different from zero, while the convergence correlation between the two Longshanks lines is strong (0.18) and statistically significant (CIs [0.07, 0.25]).

One finding in the Longshanks study was that two major-effect loci showed parallel frequency shifts between the two selection lines. We were curious to what extent our genome-wide covariances were being driven by these two outlier large-effect loci, so we excluded them from the analysis.

Since we do not know the extent to which linkage disequilibrium around these large-effect loci affects neighboring loci, we took the conservative precaution of excluding the entire chromosomes these loci reside on (chromosomes 5 and 10), and re-calculating the temporal covariances. We find excluding these large effect loci has little impact on the confidence intervals (blue confidence intervals in Figure 2C), indicating that these across-replicate covariances are indeed driven by a large number of loci. This is consistent with a signal of selection on a polygenic trait driving genome-wide change, although we note that large-effect loci can contribute to the indirect change at unlinked loci (Robertson 1961; Santiago and Caballero 1995).

The presence of an unselected control line provides an alternative way to partition the effects of linked selection and genetic drift: we can compare the total variance in allele frequency change of the control line (which excludes the effect of artificial selection on allele frequencies) to the total variance in frequency change of the Longshanks selection lines. This allows us to estimate the increase in variance in allele frequency change due to selection, which we can further partition into the effects of selection shared between selection lines and those unique to a selection line by estimating the shared effect through the observed covariance between replicates (see Materials and Methods 1 and SI Appendix, section S4 for more details). We estimate at least 32% (95% CI [21%, 48%]) of the variance in allele frequency change is driven by the effects of selection, of which 14% (95% CI [3%, 33%]) is estimated to be unique to a selection line, and 17% (95% CI [9%, 23%]) is the effect of shared selection between the two Longshanks selection lines.

We observed that in the longest study we analyzed Barghi et al. (2019), some genome-wide temporal covariances become negative at future timepoints (see the first two rows in Figure 1A). This shows that alleles that were on average going up initially are later going down in frequency, i.e. that the average direction of selection experienced by alleles has flipped. This might reflect either a change in the environment or the genetic background, due to epistatic relationships among alleles altered by frequency changes (which can occur during an optima shift; Hayward and Sella 2019) or recombination breaking up selective alleles. Such reversals in selection dynamics could be occurring at other timepoints but the signal of a change in the direction of selection at particular loci may be washed out when we calculate our genome-wide average temporal covariances. To address this limitation, we calculated the distribution of the temporal covariances over 100kb windowed covariances (Figure 3 shows these distributions pooling across all replicates; see SI Appendix, Fig. S26 for individuals replicates). The covariance estimate of each genomic window will be noisy, due to sampling and genetic drift, and the neutral distribution of the covariance is complicated due to linkage disequilibria, which can occur over long physical distances in E&R and selection studies (Baldwin-Brown et al. 2014; Nuzhdin and Turner 2013). To address this, we have developed a permutation-based procedure that constructs an empirical neutral null distribution by randomly flipping the sign of the allele frequency changes in each genomic window (i.e. a single random sign flip is applied to all loci in a window). This destroys the systematic covariances created by linked selection and creates a sampling distribution of the covariances spuriously created by neutral genetic drift while preserving the complex dependencies between adjacent loci created by linkage disequilibrium. This empirical neutral null distribution is conservative in the sense that the variances of the covariances are wider than expected under drift alone, as selection not only creates covariance between time intervals, but also inflates the magnitude of allele frequency change within a time-interval. We see (Figure 3 A and B) that there are an empirical excess of windows with positive covariances between close timepoints compared to the null distribution (a heavier right tail), and that this then shifts to an excess of windows with negative covariances between more distant timepoints (a heavier left tail).

**Figure 3:**
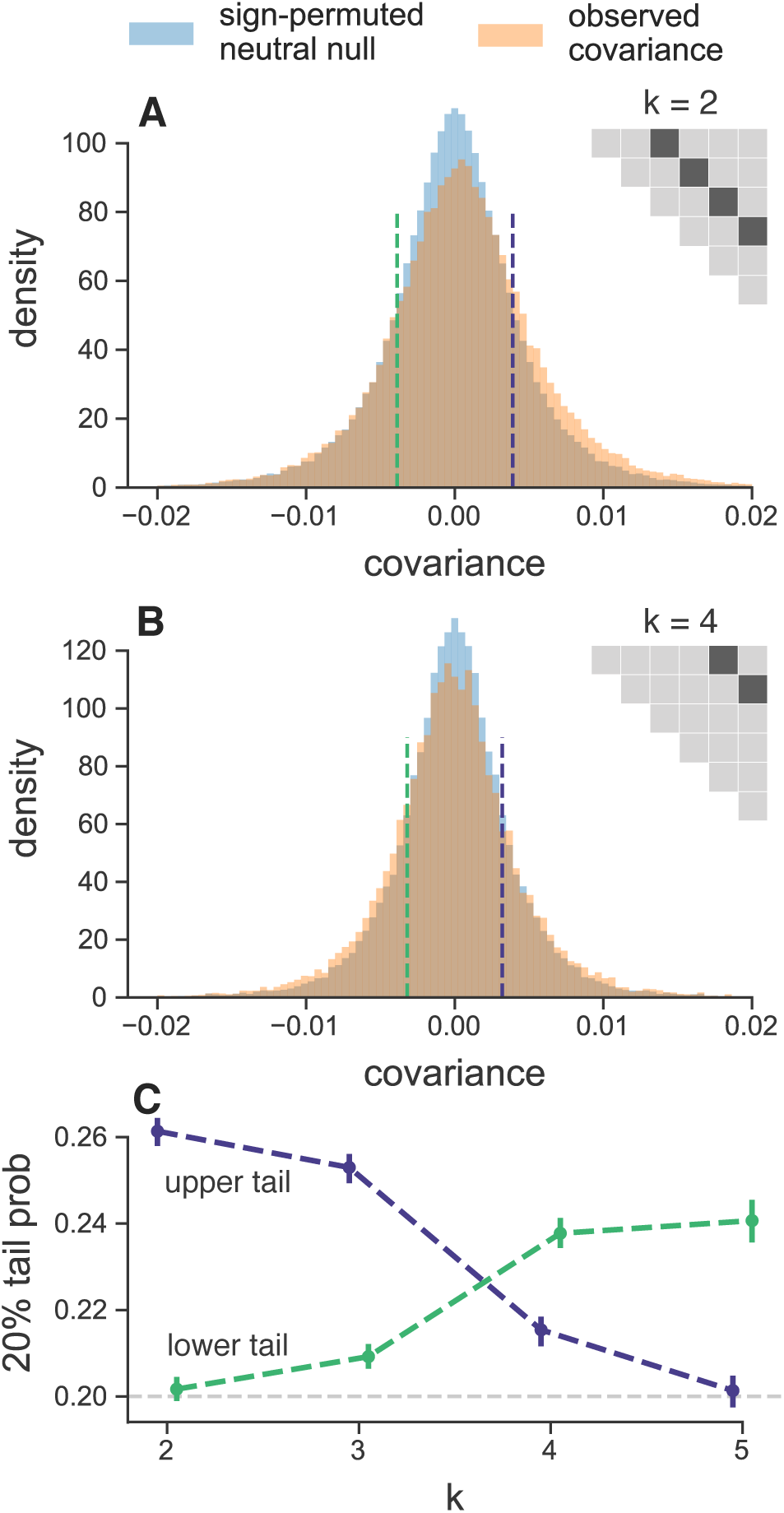
**A, B**: The distribution of temporal covariances calculated in 100kb genomic windows from the Barghi et al. (2019) study, plotted alongside an empirical neutral null distribution created by recalculating the windowed covariances on 1,000 sign permutations of allele frequency changes within tiles. The number of histogram bins is 88, chosen by cross validation (SI Appendix, section S25). In subfigure **A**, windowed covariances Cov(Δ*p*_*t*_, Δ*p*_*t*+*k*_) are separated by *k* = 2 × 10 generations and in subfigure **A** the covariances are separated by *k* = 4 × 10 generations; each *k* is an off-diagonal from the variance diagonal of the temporal covariance matrix (see cartoon of upper-triangle of covariance matrix in subfigures **A** and **B**, where the first diagonal is the variance, and the dark gray indicates which off-diagonal of the covariance matrix is plotted in the histograms). **C**: The lower and upper tail probabilities of the observed windowed covariances, at 20% and 80% quintiles of the empirical neutral null distribution, for varying time between allele frequency changes (i.e. which off-diagonal *k*). The confidence intervals are 95% block-bootstrap confidence intervals, and the light gray dashed line indicates the 20% tail probability expected under the neutral null. Similar figures for different values of *k* are in SI Appendix, section S27.

We quantified the degree to which the left and right tails are inflated compared to the null distribution as a function of time, and see excesses in both tails in Figure 3C. This finding is also robust to sign-permuting allele frequency changes on a chromosome-level, the longest extent that gametic linkage disequilibria can extend (SI Appendix, Fig. S29). We see a striking pattern that the windowed covariances not only decay towards zero, but in fact become negative through time, consistent with many regions in the genome having had a reversed fitness effect at later timepoints.

Finally we used forward-in-time simulations to explore the conditions under which temporal and convergent correlations arise. We show a subset of our results for a model of stabilizing selection on a phenotype where directional selection is induced by a sudden shift in the optimum phenotype of varying magnitudes (Figure 4A). We find that positive temporal covariances are produced by such selection (Figure 4B), and that these positive temporal covariances can compound together to generate a large proportion of allele frequency change being due to selection (i.e. large *G*(*t*)) over the relatively short time periods similar to our analyzed selection datasets span (Figure 4C). The magnitude of *G*(*t*) increases with the strength of selection, i.e. the variance in fitness, such that stronger selection generates larger proportions of allele frequency change. We find a similar picture of stronger convergent selection pressures generating larger convergence correlations (Figure 4D; see also SI Appendix, Fig. S12 for how other factors impact convergence correlations).

**Figure 4:**
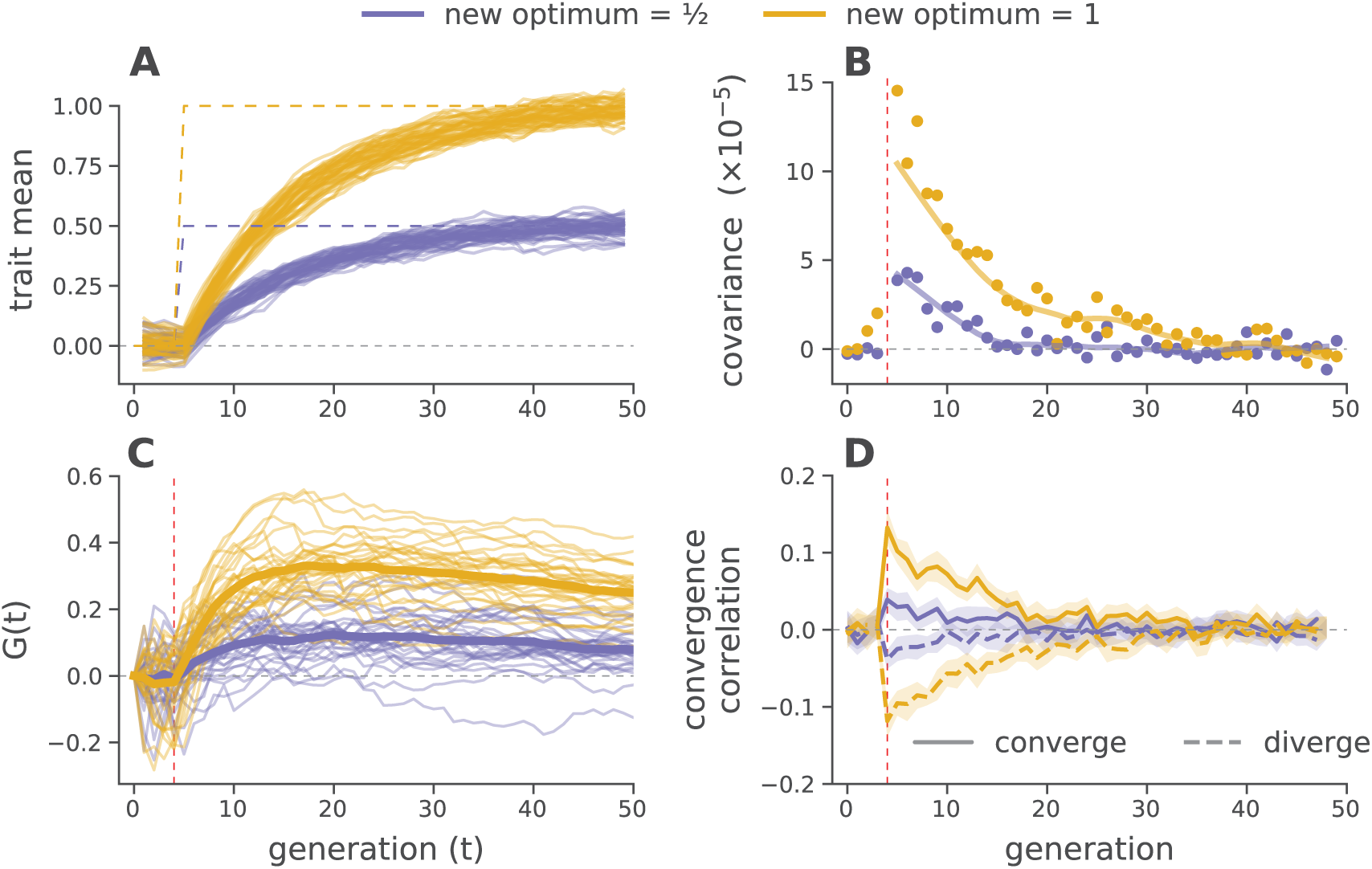
Forward-in-time simulations demonstrate how temporal covariance, *G*(*t*) trajectories, and convergence correlations arise during optima shifts of two different magnitudes, under Gaussian stabilizing selection. (**A**) Trait means across 30 replicate before and after optima shifts (solid lines), for two different magnitudes (indicated by color). The optimal trait values are indicated by the purple and yellow dashed lines. (**B**) Mean temporal covariance Cov(Δ*p*_5_, Δ*p*_*t*_) across 30 simulation replicates, where *t* varies along the x-axis (points), with a loess-smoothed average (solid line). (**C**) *G*(*t*) trajectories through time, for 30 replicate simulations across two optima shifts. The solid line is a loess-smoothed average. (**D**) The convergence correlations between two populations (each 1000 diploids) split from a common population, that underwent either an optima shift in the same direction (converge) and opposite directions (diverge) at generation five. In subfigures (**B**), (**C**), and (**D**), directional selection begins at generation five, when the optima shifts; this is indicated by the vertical dashed red line (see SI Appendix Section S8.2 for details on these simulations).

Averaging across replicates, these simulation results show *G*(*t*) is relatively insensitive to the number of loci underlying the trait. However, if only a small number of loci influence the trait, the *G*(*t*) trajectories are typically much more stochastic *across* replicates. This reaffirms that the genome-wide linked selection response we see in the Barghi et al. (2019) data is highly polygenic (compare Figure 1B to SI Appendix, Fig. S6). Furthermore, using our simulations we find that sampling only every 10 generations does indeed mean that our estimates of *G*(*t*) are an underestimate of the proportional effect of linked selection as they cannot include the covariance between closely spaced generations (see SI Appendix, Fig. S14).

Additionally, we explored other modes of selection with simulations. We find that the long term dynamics of the covariances under directional truncation selection, which generates substantial epistasis, are richer than we see under Gaussian Stabilizing Selection (GSS) and multiplicative selection (SI Appendix, Fig. S18). We also conducted simulations of purifying selection alone (i.e. background selection) and find that this can also generate positive temporal covariances (SI Appendix, Fig. S16) and under some circumstances, can even generate convergence correlations (SI Appendix, Fig. S17). Thus it is unlikely that the signatures of linked selection we see are entirely the result of the novel selection pressure the populations are exposed to, and some of this signature may be ongoing purifying selection. Only in the case of the Longshanks experiment, does the control line allow us to conclude that selection that is almost entirely due to the novel selection pressure.

While none of our experiments have selected the populations in divergent directions, in our simulations we find that such selection can generate negative convergent correlations (Figure 4D). This suggests that selection experiments combining multiple replicates, control lines, as well as divergent selection pressures might be quite informative in disentangling the contribution of particular selection pressures from genome-wide allele frequency changes.

## Discussion

Since the seminal analysis of Maynard Smith and Haig (1974) demonstrating that linked neutral diversity is reduced as an advantageous polymorphism sweeps to fixation, over four decades of theoretical and empirical research has bettered our understanding of linked selection. One underused approach to understand the genome-wide effects of selection on polygenic trait (e.g. on standing variation), stems from an early quantitative genetic model of linked selection Robertson (1961) and its later developments (Santiago and Caballero 1995, 1998; Woolliams et al. 1993; Wray and Thompson 1990; see also Barton 2000 for a comparison of these models with classic hitchhiking models). Implicit in these models is that autocovariance between allele frequency change is created when there is heritable fitness variation in the population, a signal that may be readily detected from temporal genomic data (Buffalo and Coop 2019). Depending on how many loci affect fitness, even a strong effect of linked selection may not be differentiable from genetic drift using only single contemporary population samples or looking at temporal allele frequency change at each locus in isolation. In this way, averaging summaries of temporal data allows us to sidestep the key problem of detecting selection from standing variation: that the genomic footprint leaves too soft of a signature to differentiate from a background of genetic drift. In fact we find that the temporal covariance signal is detectable even in the extremely difficult to detect case of selection on highly polygenic traits (Buffalo and Coop 2019).

It is worth building some intuition why temporal covariance allows us to detect such faint signals of polygenic linked selection from temporal genomic data. Variance in allele frequency change is subject to both drift and sampling noise, which at any single locus may swamp the temporal covariance signal due to selection, or create spurious covariances when selection is not acting. However, these spurious covariances do not share a directional signal whereas the covariances created by linked selection do; consequently, averaging across the entire genome, the temporal signal exceeds sampling noise.

Our analyses reveal that a sizable proportion of allele frequency change in these experimental evolution populations is due to the (likely indirect) action of selection. Capitalizing on replicated designs, we characterized the extent to which convergent selection pressures lead to parallel changes in allele frequencies across replicate populations, and found that a substantial proportion of the response is shared across short timescales. These likely represent substantial under-estimates of the contribution of linked selection because the studies we have reanalyzed do not sequence the population each generation, preventing us from including the effects of stronger correlations between adjacent generations. Furthermore, our estimation methods are intentionally conservative, for example they exclude the contribution of selection that does not persist across generations and selection that reverses sign; thus they can be seen as a lower bound of the effects of selection, which we have confirmed through forward-in-time simulations. Finally, through simulation results, we show that for a given level of additive genetic variance, the strengths of temporal and replicate covariances depend on the mode of selection, the details of the populations or selection experiment, and the level of linkage disequilibria, yet the level of temporal covariance is relatively invariant to the number of loci underlying fitness, as long as fitness is sufficiently polygenic.

These estimates of the contribution of selection could be refined by using patterns of linkage disequilibria (LD) and recombination which would allow us to more fully parameterize a linked-selection model of temporal allele frequency change (Buffalo and Coop 2019). The basic prediction is that regions of higher linkage disequilibrium and lower recombination should have greater temporal autocovariance than regions with lower LD and higher recombination. However, one limitation of these pooled sequence datasets is that none of the studies we reanalyzed estimated linkage disequilibria data for the evolved populations. While there are LD data for a natural population of *D. simulans* (Howie et al. 2018; Signor et al. 2018), we did not find a relationship between temporal covariance and LD. We believe this is driven by the idiosyncratic nature of LD in evolve-and-resequence populations, which often extends over large genomic distances (Kelly and Hughes 2019; Nuzhdin and Turner 2013). Future studies complete with LD data and recombination maps would allow one to disentangle the influence of closely linked sites from more distant sites in causing temporal autocovariance, and allow the fitting of more parametric models to estimate population parameters such as the additive genetic variance for fitness directly from temporal genomic data alone (Buffalo and Coop 2019). Future work could refine our *G*(*t*) estimates by including selection’s impact on the variance in allele frequency terms (e.g. see equation 26 of (Buffalo and Coop 2019)), and possibly quantifying the covariances missed when sequencing is not done each generation; both would lead to less conservative estimates that could show a large impact of selection.

Our primary focus here has been on evolution in laboratory populations. It is unclear whether we should expect a similar impact of selection in natural populations. In some of these experiments, selection pressures may have been stronger or more sustained than in natural populations (Hairston et al. 2005; Hendry and Kinnison 1999). Conversely, these lab populations were maintained at relatively small census sizes (Table 1), which will amplify the role of genetic drift, and increase the frequency of rare deleterious alleles in selection lines due to founder effects. The advantage of lab experiments is that they are closed populations; in natural populations temporal covariance could also arise from the systematic migration of alleles from differentiated populations. Adapting these methods to natural populations will require either populations that are reasonably closed to migration, or for the effect of migration to be accounted for possibly either by knowledge of allele frequencies in source populations or the identification of migrant individuals.

**Table 1:**
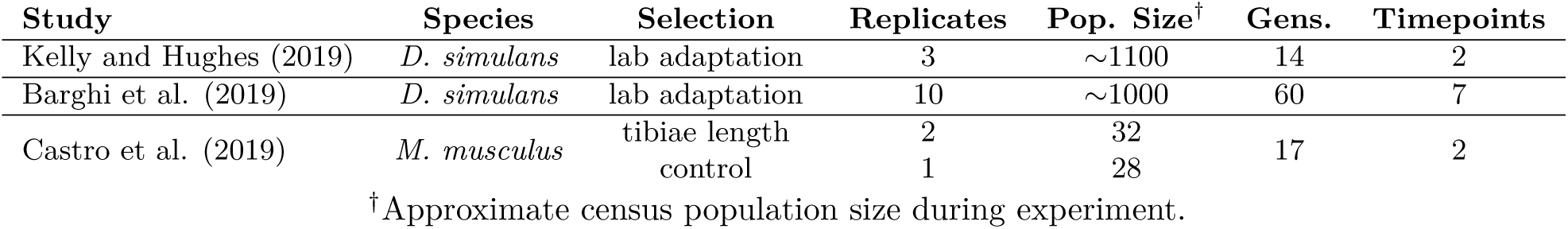
Summary of the main selection studies we analyzed

| Study | Species | Selection | Replicates | Pop. Size<sup>†</sup> | Gens. | Timepoints |
| --- | --- | --- | --- | --- | --- | --- |
| Kelly and Hughes (2019) | *D. simulans* | lab adaptation | 3 | ~1100 | 14 | 2 |
| Barghi et al. (2019) | *D. simulans* | lab adaptation | 10 | ~1000 | 60 | 7 |
| Castro et al. (2019) | *M. musculus* | tibiae length | 2 | 32 | 17 | 2 |
|  |  | control | 1 | 28 |  |  |

<sup>†</sup>Approximate census population size during experiment.

While it challenging to apply temporal methods to natural populations there is a lot of promise for these approaches (Bergland et al. 2014; Machado et al. 2018). Efforts to quantify the impact of linked selection have found obligately sexual organisms have up to an 89% reduction in genome-wide diversity over long time periods (Comeron 2014; Coop 2016; Corbett-Detig et al. 2015; Elyashiv et al. 2016; McVicker et al. 2009) Thus linked selection makes a sizeable contribution to long-term allele frequency change in some species, and there is reason to be hopeful that we could detect this from temporal data, which would help to resolve the timescales that linked selection acts over in the wild. In our reanalysis of the Barghi et al. (2019) study, we find evidence of complex linked selection dynamics, with selection pressures flipping over time due to either environmental change, the breakup of epistatic combinations or advantageous haplotypes. Such patterns would be completely obscured in samples from only contemporary populations. Thus, we can hope to have a much richer picture of the impact of selection as temporal sequencing becomes more common, allowing us to observe the effects of ecological dynamics in genomic data (Hairston et al. 2005).

Furthermore, understanding the dynamics of linked selection over short timescales will help to unite phenotypic studies of rapid adaptation with a detectable genomic signature, to address long-standing questions concerning linked selection, evolutionary quantitative genetics, and the overall impact selection has on genetic variation.

## Materials and Methods

### Datasets Analyzed

We used available genomic data data from four studies: pooled population resequencing (pool-Seq) data from Barghi et al. (2019), Kelly and Hughes (2019), and Bergland et al. (2014), and individual-level sequencing data from Castro et al. (2019). In all cases, we used the variants kept after the filtering criteria of the original studies.

### Variance and Covariance Estimates

To remove systematic covariances in allele frequency change caused by tracking the reference or minor allele, we randomly choose an allele to track frequency for each locus. Then, we calculate the variance-covariance matrix of allele frequency changes using a Python software package we have written, available at http://github.com/vsbuffalo/cvtk. This simultaneously calculates temporal variances and covariances, and replicate covariances and uses the sampling depth and number of diploid individuals to correct for bias in the variance estimates and a bias that occurs in covariance estimates between adjacent timepoints due to shared sampling noise (see SI Appendix, sections S1.2, S1.3, and S1.4 for mathematical details of these estimators). We assess that our bias correction procedure is working adequately through a series of diagnostic plots that ensure that the procedure removes the relationship between sampling depth and uncorrected variance and covariances (SI Appendix, Fig. S4). Through our simulations we find that our estimates can differ based on how fixations and losses are handled in long time-series (SI Appendix, section S8.7) but none of our findings in the main text are qualitatively altered by this decision (SI Appendix, Figs. S19 and S20).

### Estimating Uncertainty with a Block Bootstrap

To infer the uncertainty of covariance, convergence correlation, and *G*(*t*) estimates, we used a block bootstrap procedure. This bootstrap procedure resamples blocks of loci, rather than individual loci, to infer the uncertainty of a statistic in the presence of unknown correlation between loci. As most estimators in this paper are ratios (e.g. covariance standardize by sample heterozygosity, *G*(*t*), and the convergence correlation), which we estimate with a ratio of averages, we exploit the linearity of expectation for efficient computation of bootstrap samples (see SI Appendix, Fig. S3 for details).

### Partitioning Unique and Shared Selection Effects in the Longshanks Study

The unselected control line in the Longshanks experiment allows us to additionally partition the total variance in allele frequency change into drift, shared effects of selection, and unshared effects of selection between selected replicates. We begin by decomposing the allele frequency change in Longshanks line 1 (LS1) as Δ*p*_*t*,LS1_ = Δ_*D*_ *p*_*t*,LS1_ + Δ_*U*_ *p*_*t*,LS1_ + Δ_*S*_ *p*_*t*,LS_ where these terms are the drift in Longshanks replicate 1 (Δ_*D*_ *p*_*t*,LS1_), selection unique to the LS1 replicate (Δ_*U*_ *p*_*t*,LS1_), and selection response shared between the two Longshanks replicates (Δ_*S*_ *p*_*t*,LS_) respectively (and similarly for the Longshanks line 2, LS2). By construction, this decomposition assumes that each of these terms are uncorrelated within replicates, so the contribution of each term to the total variance in allele frequency change, Var(Δ*p*_*t*,LS1_), is the variance of that term’s allele frequency change.

We estimate the effects of selection by first calculating the fraction of the total variance explained by drift. We assume the variance in allele frequency change observed in the unselected control line (Var(Δ*p*_*t*,Ctrl_)) is driven entirely by neutral genetic drift, and since an identical breeding scheme was used across all three replicates (except breeders for the control line were chosen at random), we can use this as an estimate of the contribution of neutral genetic drift in the selected lines, Var(Δ*p*_*t*,Ctrl_) = Var(Δ_*D*_ *p*_*t*,LS1_) = Var(Δ_*D*_ *p*_*t*,LS2_). Then, we can estimate the increase in variance in allele frequency change due to selection as (Var(Δ*p*_*t*,LS1_) + Var(Δ*p*_*t*,LS2_))*/*2 − Var(Δ*p*_*t*,Ctrl_) and the shared effect of selection across selected lines as Cov(Δ*p*_*t*,LS1_, Δ*p*_*t*,LS2_). Finally, the covariance in allele change between replicates is used to estimate the shared effects of selection between lines, Cov(Δ*p*_*t*,LS1_, Δ*p*_*t*,LS2_) = Var(Δ_*S*_ *p*_*t*,LS_).

### Windowed Covariance and the Empirical Neutral Null

Throughout the paper, we use genomic windows for the block-bootstrap procedure. For the *D. simulans* and *D. melanogaster* data from the Barghi et al. (2019), Kelly and Hughes (2019), and Bergland et al. (2014) studies, we used large megabase windows for the block bootstrap procedure, while we used a ten megabase window for the large mouse genome data from the Castro et al. (2019) study.

Given evidence of a reversal in the direction of selection at later timepoints in the Barghi et al. (2019) study, we calculated windowed temporal covariances on 10 kilobase windows and looked at the distribution of these covariances through time. We compare these distributions of windowed covariances to an empirical neutral null created by randomly permuting the sign of allele frequency change at the block level (to preserve the correlation structure between loci due to LD). This destroys the systematic covariances in allele frequency change created by linked selection, which emulates a frequency trajectory under drift. This approach is conservative, since heritable fitness variation also inflates the magnitude of allele frequency change more than expected under drift, but we do not change these magnitudes. Using this empirical neutral null distribution of windowed covariances, we calculate how much of the observed windowed covariance distribution falls outside of empirical null distribution for different tail probabilities. While the comparison between the distribution of 10 kilobase windowed covariances to the empirical neutral null created from sign-permuting 10 kilobase windows is most natural, we wanted to ensure that our finding that the shift from mostly positive to mostly negative windowed covariances through time (Figure 3) was robust to LD extending beyond the range of these 10 kilobase windows. We took the conservative approach of also sign-permuting at the chromosome-level, and found the same qualitative shift (SI Appendix, Fig. S29).

### Forward-in-time Simulations

To explore how aspects of genetic architecture, models of selection, and experimental design impact temporal covariance, the *G*(*t*) trajectories, and convergence correlations, we ran extensive forward-in-time simulations using SLiM (Haller and Messer 2019); here we discuss the Gaussian Stabilizing Selection simulations in Figured 4, but SI Appendix, section S8 describes these simulation routines and others in detail.

We simulated directional selection on a trait by first evolving each population of *N* = 1000 diploids to equilibrium (we will refer to this as the burnin hereafter) under GSS for 10*N* generations with the stabilizing selection variance *V*_*s*_ = 1 and an optima set at zero. We note that the small burnin population size means that these simulations should not be taken as reflecting any specific natural population and they are for illustrative purposes only. We simulated a polygenic architecture by setting the trait mutation rate to 10^−8^ per basepair, per generation, in addition to having a separate neutral mutation of 10^−8^ which created neutral mutations which we used to calculate the temporal covariances. Our simulated region was 50 megabases in length (about one quarter of a *Drosophila* chromosome), and trait alleles were randomly selected to have a *±* 0.01 effect size. By tracking the trait mean through the burnin, we found it converged to the optimum as expected. After the burnin, the population was split into two different replicate populations, to capture the effect of bottlenecks in selection experiments (these population sizes varied between 50, 500, and 1000 diploids; the later representing no bottleneck). Each population then underwent an optima shift of either 0.1, 0.5, or 1 on generation five, with the first four generations serving as a control. These optima shifts were either in the same direction (converging), different directions (diverging), or only one optima shifted (as a control). By tracking the trait mean, we saw that it converged as expected during burnin, and the trait showed the expected directional response to selection (SI Appendix, Fig. S7). Using the neutral population frequency data from these simulations, we calculated the temporal covariances, *G*(*t*) trajectories, and convergence correlations.

## Supporting information

revision difference

## Data Availability

All analysis was done in Python, using numpy, matplotlib, and Jupyter notebooks (Hunter 2007; Kluyver et al. 2016; Oliphant 2006; Rossum 1995); code to reproduce these analyses is available on Github, https://github.com/vsbuffalo/cvtk/. All data is from previous studies and available; Barghi et al. (2019) data was downloaded from https://datadryad.org/resource/doi:10.5061/dryad.rr137kn, Kelly and Hughes (2019) data was downloaded from https://gsajournals.figshare.com/articles/Supplemental_Material_for_Kelly_and_Hughes_2018/7124963, Bergland et al. (2014) data was downloaded from https://datadryad.org/stash/dataset/doi:10.5061/dryad.v883p, and Castro et al. (2019) data was downloaded from http://ftp.tuebingen.mpg.de/fml/ag-chan/Longshanks/.

## Acknowledgments

We would like to thank the authors of the original studies we have analyzed, including Neda Barghi, Nick Barton, Alan Bergland, Frank Chan, Kimberly Hughes, John Kelly, Dmitri Petrov, Campbell Rolian, Christian Schlötterer. We would also like to thank Doc Edge for helpful statistical advice, and Dave Begun, Erin Calfee, Sarah Friedman, Andy Kern, Chuck Langley, Michael Turelli, Matt Osmond, Peter Ralph, and Sivan Yair for helpful discussions. Additionally, we thank Guy Sella and an anonymous reviewer whose comments greatly improved the manuscript. This research was supported by an NSF Graduate Research Fellowship grant awarded to VB (1650042), and NIH (R01-GM108779) and NSF (1353380) awarded to GC.

## Supplementary Information

### S1 Estimator Bias Correction

#### S1.1 Correcting variance bias with a single depth sampling process

Following Waples (1989), we have that the variance in allele frequency change at a locus in the initial generation, which is entirely due to a binomial sampling process, is Var(*p*_0_) = *p*_0_(1−*p*_0_)*/d*_0_ where *d*_0_ is the number of binomial draws (e.g. read depth). At a later timepoint, the variance in allele frequency is a result of both the binomial sampling process at time *t* and the evolutionary process. Using the law of total variation we can partition the variation from each process,

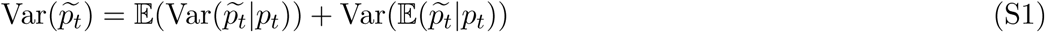

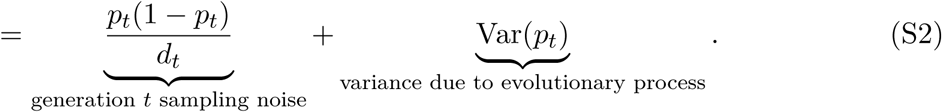

Under a drift-only process, 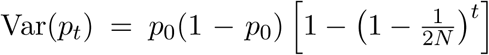. However, with heritable variation in fitness, we need to consider the covariance in allele frequency changes across generations (Buffalo and Coop 2019). We can write

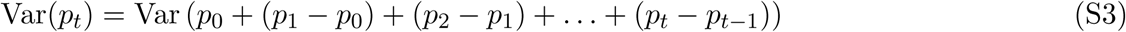

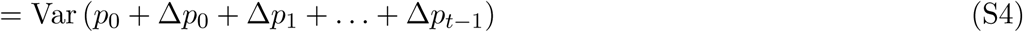

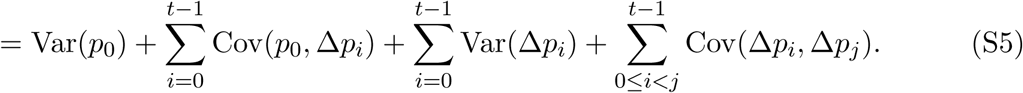

Each allele frequency change is equally like to be positive as it is to be negative; thus by symmetry this second term is zero. Additionally Var(*p*_0_) = 0, as we treat *p*_0_ as a fixed initial frequency. We can write,

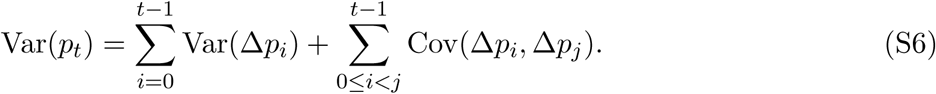

The second term, the cumulative impact of variance in allele frequency change can be partitioned into heritable fitness and drift components (Buffalo and Coop 2019; Santiago and Caballero 1995)

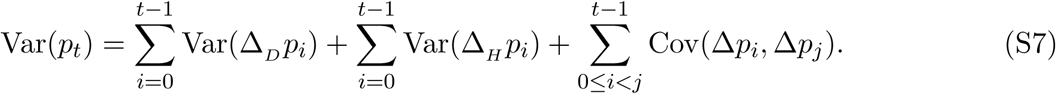

where Δ_H_ *p*_*t*_ and Δ_D_ *p*_*t*_ indicate the allele frequency change due to heritable fitness variation and drift respectively. Then, sum of drift variances in allele frequency change is

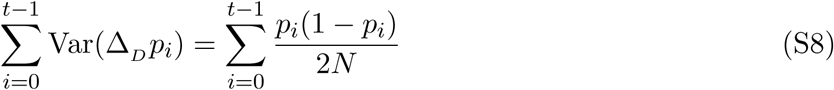

replacing the heterozygosity in generation *i* with its expectation, we have

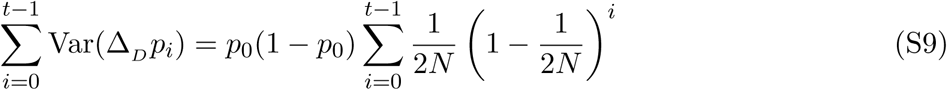

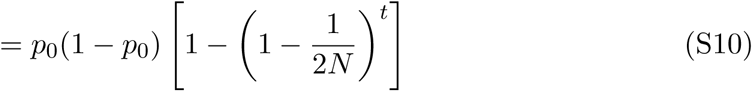

which is the usual variance in allele frequency change due to drift. Then, the total allele frequency change from generations 0 to *t* is 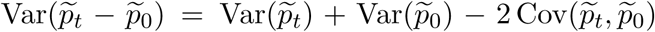, where the covariance depends on the nature of the sampling plan (see Nei and Tajima 1981; Waples 1989). In the case where there is heritable variation for fitness, and using the fact that 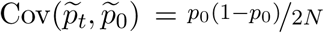 for Plan I sampling procedures (Waples 1989), we write,

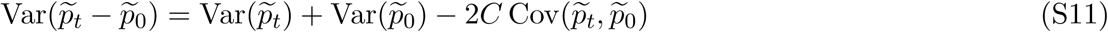

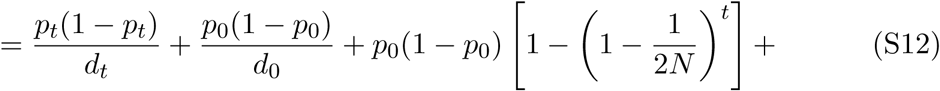

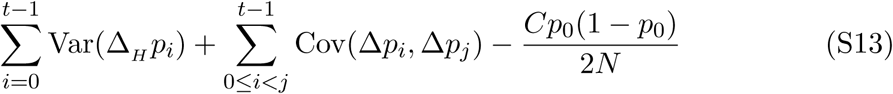

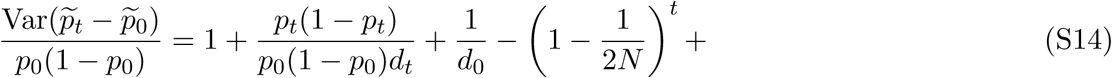

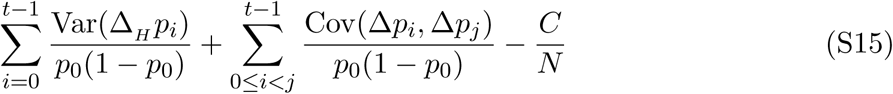

where *C* = 1 if Plan I is used, and *C* = 0 if Plan II is used (see Waples 1989, p. 380 and Figure 1 for a description of these sampling procedures; throughout the paper we use sampling Plan II). Rearranging, we can create a bias-corrected estimator for the population variance in allele frequency change, and replace all population heterozygosity terms with the unbiased sample estimators, e.g. 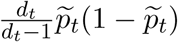,

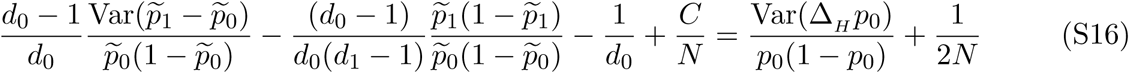

#### S1.2 Correcting variance bias with individual and depth sampling processes

Here, we extend the sampling bias correction described above to handle two binomial sampling processes: one as individuals are binomially sampled from the population, and another as reads are binomially sampled during sequencing. (see also Jónás et al. 2016). Let *X*_*t*_ ∼ Binom(*n*_*t*_, *p*_*t*_) where *X*_*t*_ is the count of alleles and *n*_*t*_ is the number of diploids sampled at time *t*. Then, these individuals are sequenced at a depth of *d*_*t*_, and *Y*_*t*_ ∼ Binom(*d*_*t*_, *X*_*t*_*/n*_*t*_) reads have the tracked allele. We let 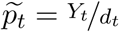 be the observed sample allele frequency. Then, the sampling noise is

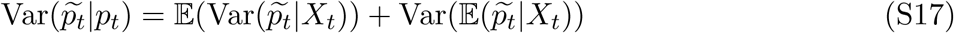

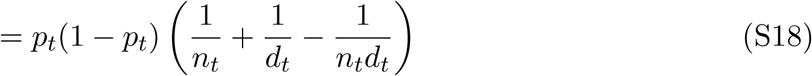

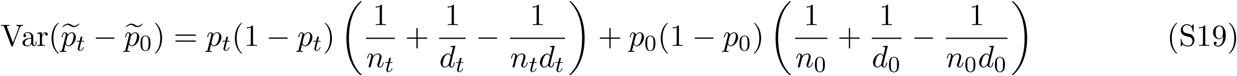

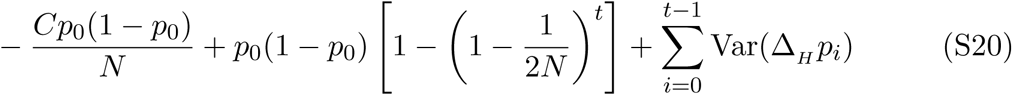

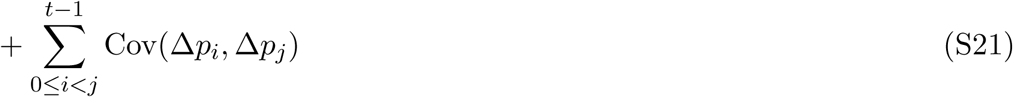

Through the law of total expectation (see Kolaczkowski et al. 2011 Supplementary File 1 for a sample proof), one can find that an unbiased estimator of the half the heterozygosity is

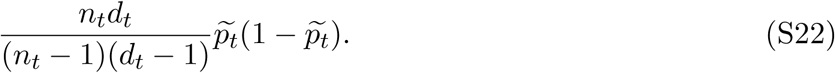

Replacing this unbiased estimator for half of the heterozygosity into our expression above, the total sample variance is

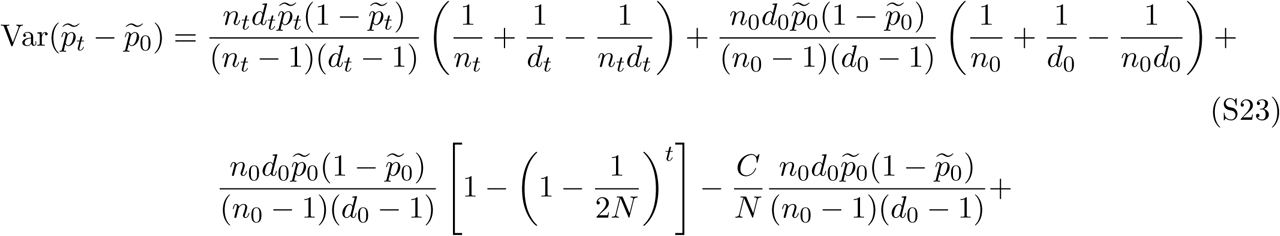

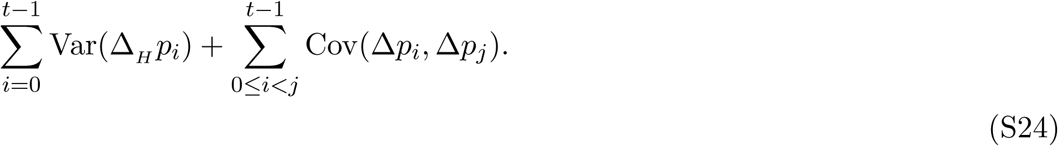

As with equation (S16), we can rearrange this to get a biased-corrected estimate of the variance in allele frequency change between adjacent generations, Var(Δ*p*_*t*_).

#### S1.3 Covariance Correction

We also need to apply a bias correction to the temporal covariances (and possibly the replicate covariances if the initial sample frequencies are all shared). The basic issue is that 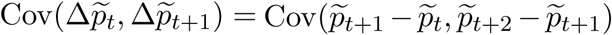, and thus shares the sampling noise of timepoint *t* + 1. Thus acts to bias the covariance by subtracting off the noise variance term of 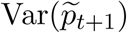, so we add the expectation of this bias, derived above, back in. We discuss this in more detail below in deriving the bias correction for the temporal-replicate variance covariance matrix.

#### S1.4 Temporal-Replicate Covariance Matrix Correction

In practice, we simultaneously estimate the temporal and replicate covariance matrices for each replicate, which we call the temporal-replicate covariance matrix. This needs a bias correction; we extend the bias corrections for single locus variance and covariance described in Supplementary Material Sections S1.1, S1.2, and S1.3 to multiple sampled loci and the temporal-replicate covariance matrix here. With frequency data collected at *T* + 1 timepoints across *R* replicate populations at *L* loci, we have multidimensional arrays **F** of allele frequencies, **D** of sequencing depths, and **N** of the number of individuals sequenced, each of dimension *R* × (*T* + 1) × *L*. We calculate the array **ΔF** which contains the allele frequency changes between adjacent generations, and has dimension *R* × *T* × *L*. The operation flat(**ΔF**) flattens this array to a (*R* · *T*) × *L* matrix, such that rows are grouped by replicate, e.g. for timepoint *t*, replicate *r*, and locus *l* such that for allele frequencies *p*_*t,r,l*_, the frequency change entries are

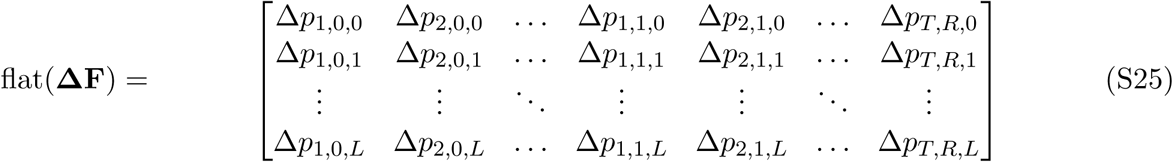

where each Δ*p*_*t,r,l*_ = *p*_*t*+1,*r,l*_ − *p*_*t,r,l*_. Then, the sample temporal-replicate covariance matrix **Q**′ calculated on flat(**ΔF**) is a (*R* ·*T*) × (*R* ·*T*) matrix, with the *R* temporal-covariance block submatrices along the diagonal, and the *R*(*R* − 1) replicate-covariance submatrices matrices in the upper and lower triangles of the matrix,

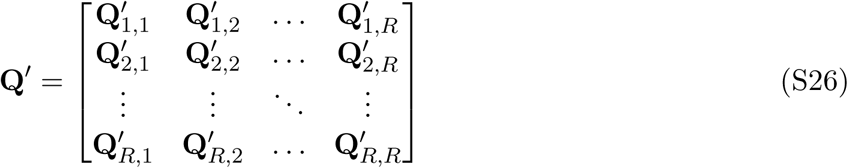

where each submatrix 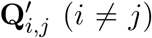 is the *T* × *T* sample replicate covariance matrix for replicates *i* and *j*, and the submatrices along the diagonal 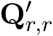 are the temporal covariance matrices for replicate *r*.

Given the bias of the sample covariance of allele frequency changes, we calculated an expected bias matrix **B**, averaging over loci,

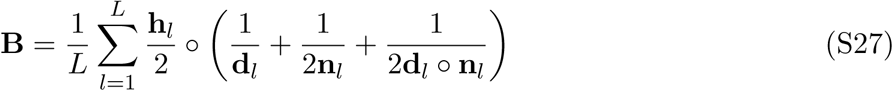

where ∘ denotes elementwise product, and **h**_*l*_, **d**_*l*_, and **n**_*l*_, are rows corresponding to locus *l* of the unbiased heterozygosity arrays **H**, depth matrix **D**, and number of diploids matrix **N**. The unbiased *R* × (*T* + 1) × *L* heterozygosity array can be calculated as

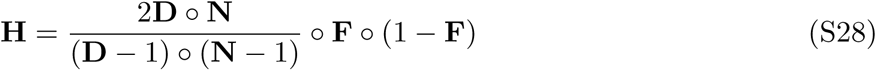

where division here is elementwise. Thus, **B** is a *R* × (*T* +1) matrix. As explained in Supplementary Material Section S1.2 and S1.3, the temporal variances and covariances require bias corrections, meaning each temporal covariance submatrix **Q**_*r,r*_ requires two corrections. For an element *Q*_*r,t,s*_ = Cov(Δ*p*_*t*_, Δ*p*_*s*_) of the temporal covariance submatrix for replicate *r*, **Q**_*r,r*_, we apply the following correction

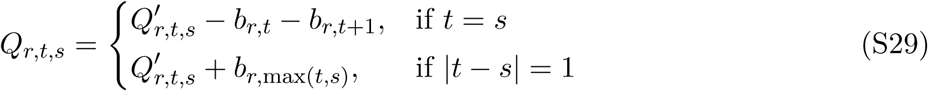

where *b*_*r,t*_ is element in row *r* and column *t* of **B**.

### S2 Barghi et al. (2019) Temporal Covariances

Since each replicate population was sequenced every ten generations, the timepoints *t*_0_ = 0 generations, *t*_1_ = 10 generations, *t*_2_ = 20 generations, etc., lead to observed allele frequency changes across ten generation blocks,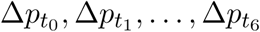. Consequently, the ten temporal covariance matrices for each of the ten replicate populations have off-diagonal elements of the form 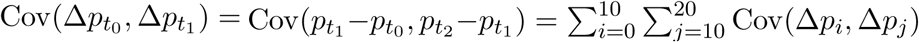. Each diagonal element has the form 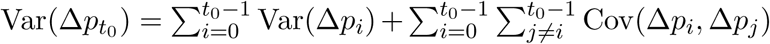, and is thus a combination of the effects of drift and selection, as both the variance in allele frequency changes and cumulative temporal autocovariances terms increase the variance in allele frequency. With sampling each generation, one could more accurately partition the total variance in allele frequency change (Buffalo and Coop 2019); while we cannot directly estimate the contribution of linked selection to the variance in allele frequency change here, the presence of a positive observed covariance between allele frequency change can only be caused linked selection.

### S3 Block Bootstrap Procedure

The estimators used in this paper are predominantly ratios, e.g. temporal-replicate covariance standardized by half the heterozygosity, *G*(*t*) which is the ratio of cumulative covariance to total variance, and the convergence correlation (equation (2)). In these cases, we can exploit the linearity of the expectation to make the bootstrap procedure more computationally efficient, by pre-calculating the statistics of the ratio’s numerator and denominator, *N* (**x**_*i*_) and *D*(**x**_*i*_), on the data **x**_*i*_ for all blocks *i* ∈ {1, 2, …, *W*} in the genome. Then we draw *W* bootstrap samples with replacement, and compute the estimate for bootstrap sample *b* with an average weighted by the fraction *w*_*i*_ of total loci contained in each block,

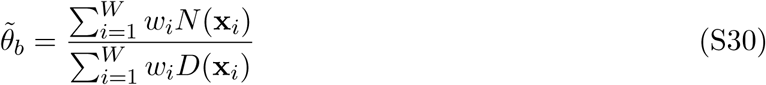

Note that computing the ratio of averages rather than the average of a ratio is a practice common for population genetic statistics like *F*_*ST*_ (Bhatia et al. 2013). With these *B* bootstrap estimates, we calculate the *α/*2 and 1 − *α/*2 quantiles, which we use to estimate the 1 − *α* = 95% pivot confidence intervals (p. 33 Wasserman 2006, p. 194 Davison and Hinkley 2013) throughout the paper,

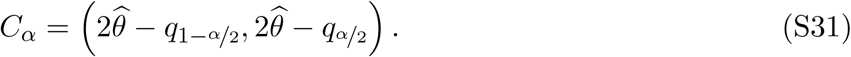

where 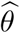 is the estimate, and *q*_*x*_ is bootstrap quantile for probability *x*.

### S4 Replicate *G*(*t*) and Partitioning the Variance in Allele Frequency

We define a statistic similar to *G*(*t*) for estimating the proportion of allele frequency change common between two replicate populations due to linked selection. Covariance in allele frequency change between two replicate populations is due to convergent selection pressure selecting haplotypes shared between the two replicate populations, which acts to perturb linked neutral variation in parallel way.

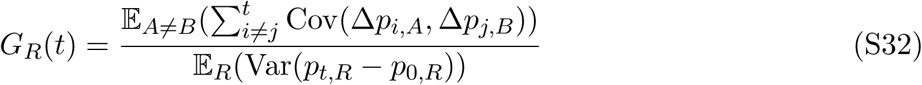

where 𝔼_*A*≠*B*_ indicates that the expectation is taken over all ordered pairs of replicates (e.g. summing all off-diagonal elements replicate covariances), and 𝔼_*R*_ indicates taking expectation over all replicates. This measures the fraction of variance in allele frequency change (averaged across replicates) due to shared selection pressure.

Extending our theoretic work in Buffalo and Coop (2019), we can partition the allele frequency change in two replicates into drift, and shared selection and replicate-specific selection components of allele frequency change. For two replicates, *A* and *B*,

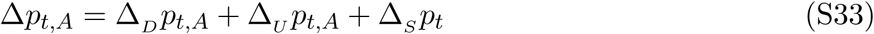

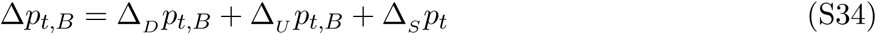

where Δ_*D*_ *p*_*t,A*_ is allele frequency change due to drift (this is specific to a replicate, and equal to Δ_*N*_ *p*_*t,A*_ + Δ_*M*_ *p*_*t,A*_ in the notation of Buffalo and Coop 2019), Δ_*U*_ *p*_*t,A*_ is the allele frequency change from indirect selection specific to replicate *A* (and likewise with Δ_*U*_ *p*_*t,A*_ for replicate *B*), and Δ_*S*_ *p*_*t*_ is the allele frequency change from indirect selection shared across the replicates *A* and *B* (this term lacks a replicate subscript since by construction it is identical between replicates). By construction, each of these terms is uncorrelated, so the variance and be written as:

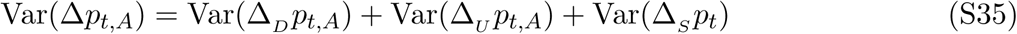

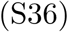

The shared effects of indirect selection can be quantified from the observed allele frequency changes, since the covariance in allele frequency change across replicates is the covariance of the shared term by construction,

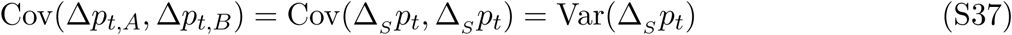

In artificial selection studies with a control (non-selected) line, such as the Castro et al. (2019) study, this allows us to estimate the contribution of the effects of shared and unique indirect selection. In the case of this study, we can estimate the drift, unique selection effect, and shared selection effect terms using the fact that,

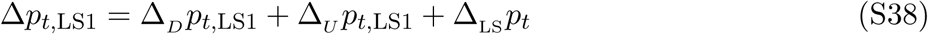

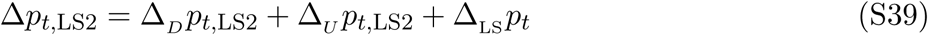

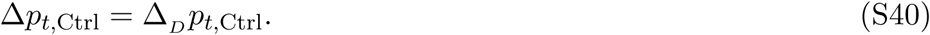

Note that since the control replicate does not undergo artificial selection, we assume that its allele frequency changes are determined entirely by genetic drift. With free mating individuals (such as in a cage population), this may not be the case, and sequencing adjacent generations would allow one to differentiate the effects of selection and drift.

We assume that we can approximate the contribution of genetic drift in the Longshanks selection lines with the observed variance in the control line, or Var(Δ*p*_*t*,Ctrl_) = Var(Δ_*D*_*p*_*t*,LS1_) = Var(Δ _*D*_*p*_*t*,LS2_). Then, the combined effects of selection can be estimated by averaging the variances of the two Longshanks selection lines, and subtracting the variance in allele frequency change in the control line, which we treat as driven by drift alone (since matings are random). Note that each variance is bias-corrected according to the methods described in Supplementary Materials S1.4, and the average sequencing depths between lines are nearly identical. Thus, we have

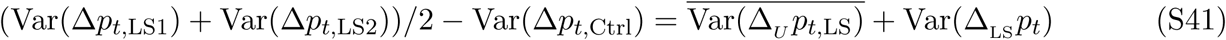

where the bar indicates values averaged both Longshanks selection lines. Additionally, use the fact that

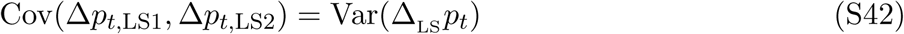

we can also separate out the unique and shared components by subtracting off this covariance,

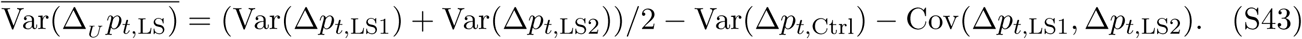

Finally, we can divide each of these values by the total variance to get the proportion of total variance drift, and unique and shared effects of selection contribute towards the total. To derive confidence intervals for the estimates of unique and shared effects of selection, we use a block bootstrap procedure as described in Supplementary Materials Section S3.

### S5 The Empirical Neutral Null Windowed Covariance Distribution

To detect an excess of genomic regions with unusually high or low covariances, we need to compare the distribution of observed windowed covariances to a null distribution of windowed covariances that we would expect under no selection. While we could construct a theoretic sampling distribution of the spurious covariances created by neutral genetic drift at particular site, the unknown linkage disequilibrium between sites would mean that this is not an adequate null model for the distribution of windowed covariances in our data.

To address this limitation, we construct a neutral null model by sign-permuting the observed allele frequency changes. This destroys the covariances built up by selection, mimicking a neutral allele’s frequency trajectory. This approach is conservative, since selection also acts to increase the magnitude of allele frequency changes (see equation 1 of Buffalo and Coop 2019), but this magnitude is not affected by the sign-permutation procedure. Consequently, the resulting empirical null distribution has higher variance than would be expected under neutrality alone.

Still, we wanted to ensure that LD between sign-permuted blocks, which will affect the variance of the empirical null distribution, does not impact our primary finding that the distribution of temporal covariances becomes increasingly negative in the Barghi et al. (2019) dataset through time. To address this, we also sign-permuted at the whole chromosome level finding we recapitulate the same pattern (Supplementary Figure S29).

### S6 Bergland et al. (2014) Re-Analysis

We also applied our temporal covariance approach to Bergland et al. (2014), which found evidence of genome-wide fluctuating selection between Spring and Fall seasons across three years *Drosophila melanogaster*. As described in Buffalo and Coop (2019), if fluctuating selection pressure among time-periods are the dominant genome-wide pattern, we might expect positive covariances between like seasons changes (e.g. Spring 2010 to Fall 2010 and Spring 2011 to Fall 2011), and negative covariances between dislike seasonal changes (e.g. Fall 2009 to Spring 2010 and Fall 2010 to Spring 2011). However, while we find temporal covariances that are non-zero, we find only weak support for a seasonal fluctuating model driving these covariances. In Supplementary Figure S1, we show the temporal covariances from varying reference generations, across seasonal transitions that are alike (e.g. the covariance between the allele frequency changes between Fall 2009 and Spring 2009, and frequency changes between Fall 2010 and Spring 2010), and dislike (e.g. the covariance between the allele frequency change between Fall 2009 and Spring 2009, and the frequency changes between Spring 2010 and Fall 2009). The first row of temporal covariance matrix is consistent with fluctuating selection operating for two timepoints, as the first covariance is negative, and the second is positive, and later covariances are not statistically differentiable from zero (which could occur if LD and additive genetic variance decay). However, all other temporal covariances do not fit the pattern we would expect under genome-wide fluctuating selection.

We wanted to establish that our temporal-covariance matrix bias correction was working correctly. We find that it corrects the relationship between depth and both variance and covariance (Supplementary Figure S4) as expected.

It is unclear how strong the fluctuations would have to be to generate a genome-wide average signal of fluctuating selection from temporal covariances. For example, many loci could still show a signal of fluctuating selection, but the average signal could be overwhelmed by other signals of other selection. To investigate whether there was a genome-wide excess of loci showing evidence of fluctuating selection we reanalyzed the data of (Bergland et al. 2014) using the same seasonal fluctuating model as the original paper. This model is a Binomial logit-linked GLM fit per-locus, where the frequencies are regressed on the Spring/Fall seasons are encoded as a dummy variable. We use the same binomial weighting procedure as Bergland et al. (2014), where the weights are determined by the effective number of chromosomes, *N*_*eff*_ = (2*n*_*t*_*d*_*t*_ − 1)*/*(2*n*_*t*_ + *d*_*t*_) (*n*_*t*_ and *d*_*t*_ are the number of diploid individuals and the read depth at timepoint *t*, respectively). We fit this model on all loci marked as used in the VCF provided with the Bergland et al. (2014) study (doi:10.5061/dryad.v883p). Overall, our p-values for the Wald test for each locus closely match those of the original paper (Pearson correlation coefficient 0.98, p-value < 2.2 × 10^−16^; see Supplementary Figure S2 A), and the histograms of the p-values are nearly identical (Supplementary Figure S2 B). Bergland et al. (2014) find loci with a significant association with season after a Benjamini and Hochberg FDR p-value adjustment (Benjamini and Hochberg 1995), however, the null hypothesis of the Wald test does not give us an idea of the expected number of variants that may spuriously fit the pattern of seasonal fluctuating selection as it does not account for genetic drift or other forms of hitchhiking.

**Figure S1:**
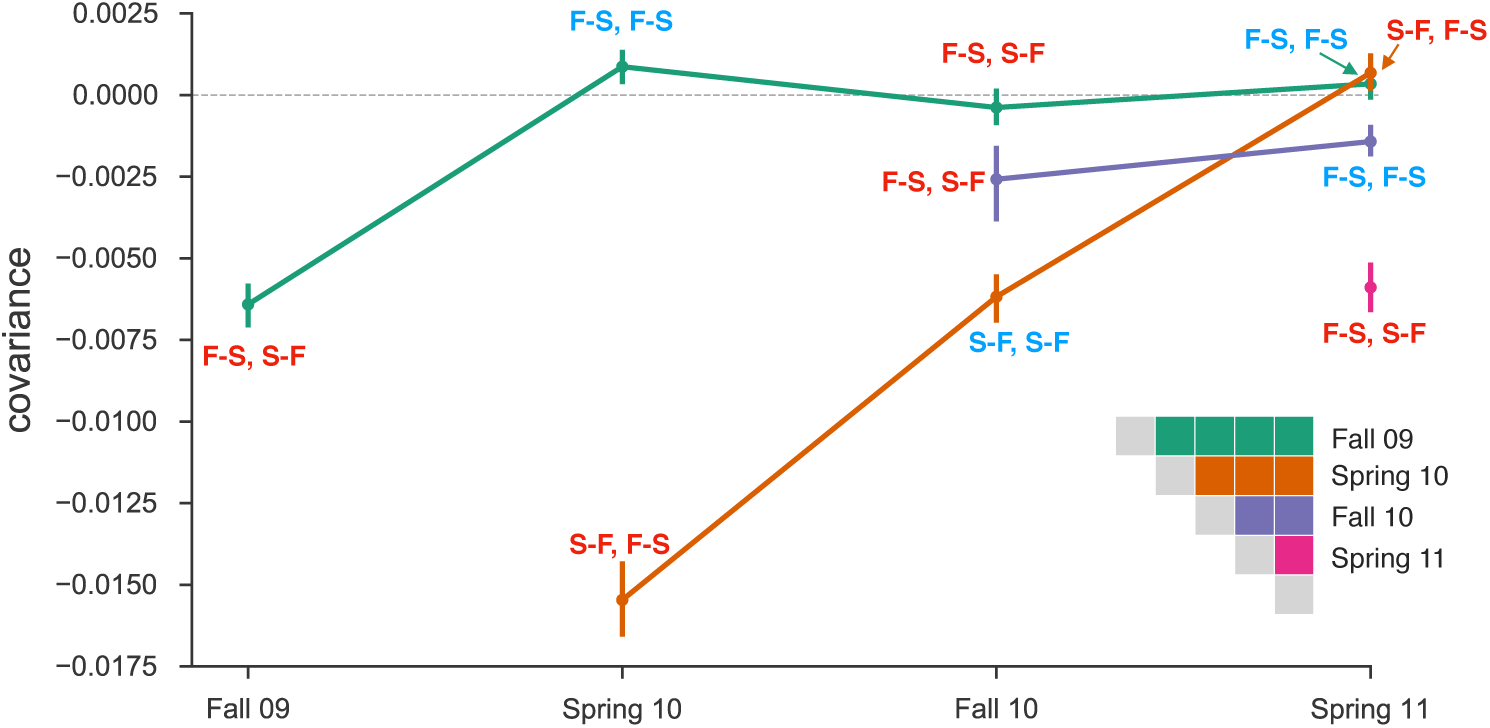
Temporal covariances from the Bergland et al. (2014) study, from varying reference generations (e.g. rows along the temporal covariance matrix). Each covariance is labeled indicating whether the covariance is between two like seasonal transitions (e.g. the covariance between allele frequency changes from fall to spring in one year, and fall to spring in another) or two dislike seasons (e.g. the covariance between fall to spring in one year, and spring to fall in another year). Covariances between like transitions are expected to be positive when there is a genome-wide effect of fluctuating selection (and these labels are colored blue), while covariances between dislike transitions are expected to be negative (and these labels are colored red). 95% confidence intervals were constructed by a block-bootstrapping procedure where the blocks are megabase tiles.

**Figure S2:**
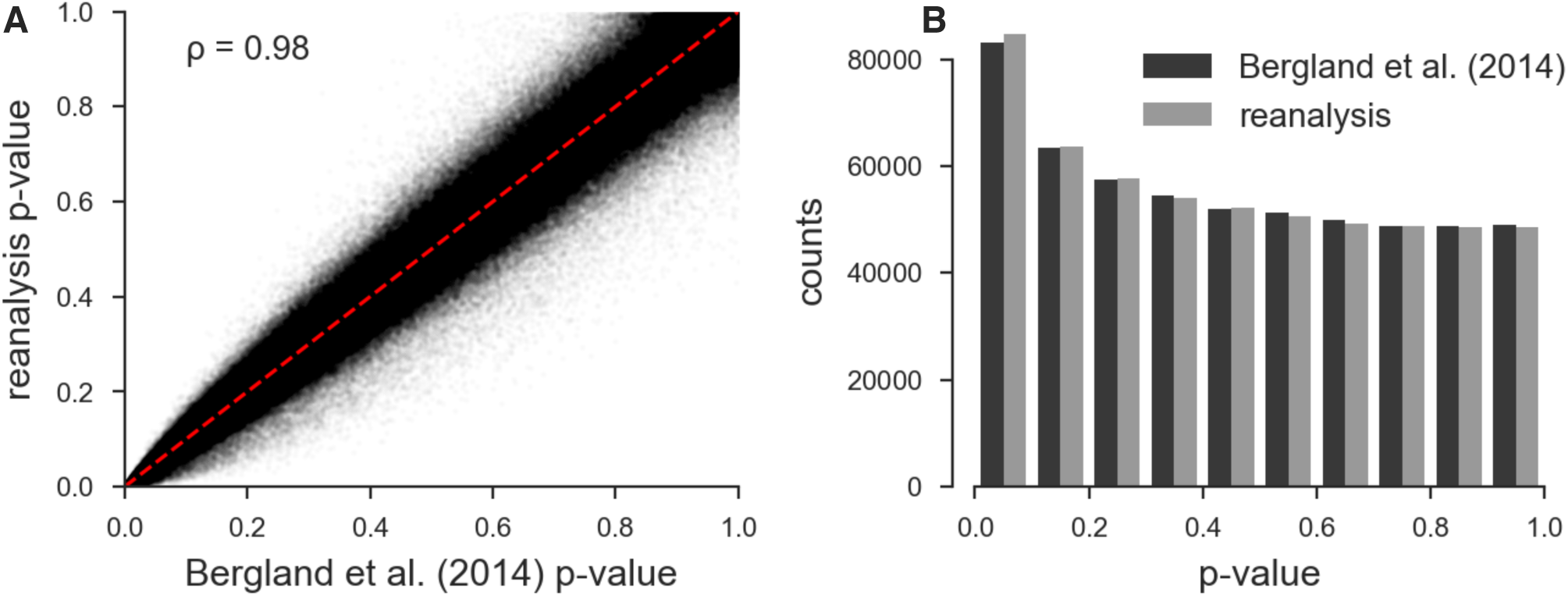
**A**: Scatterplot of the original unadjusted p-values from Bergland et al. (2014) and the p-values from our reanalysis of the same data using the same statistical methods; the minor discrepancy is likely due to software version differences. **B**: The histograms of the p-values of our reanalysis and the original Bergland et al. (2014) data; again the minor discrepancy is likely due to software differences. Overall, our implementation of Bergland et al.’s statistical methods produces results very close to the original analysis.

**Figure S3:**
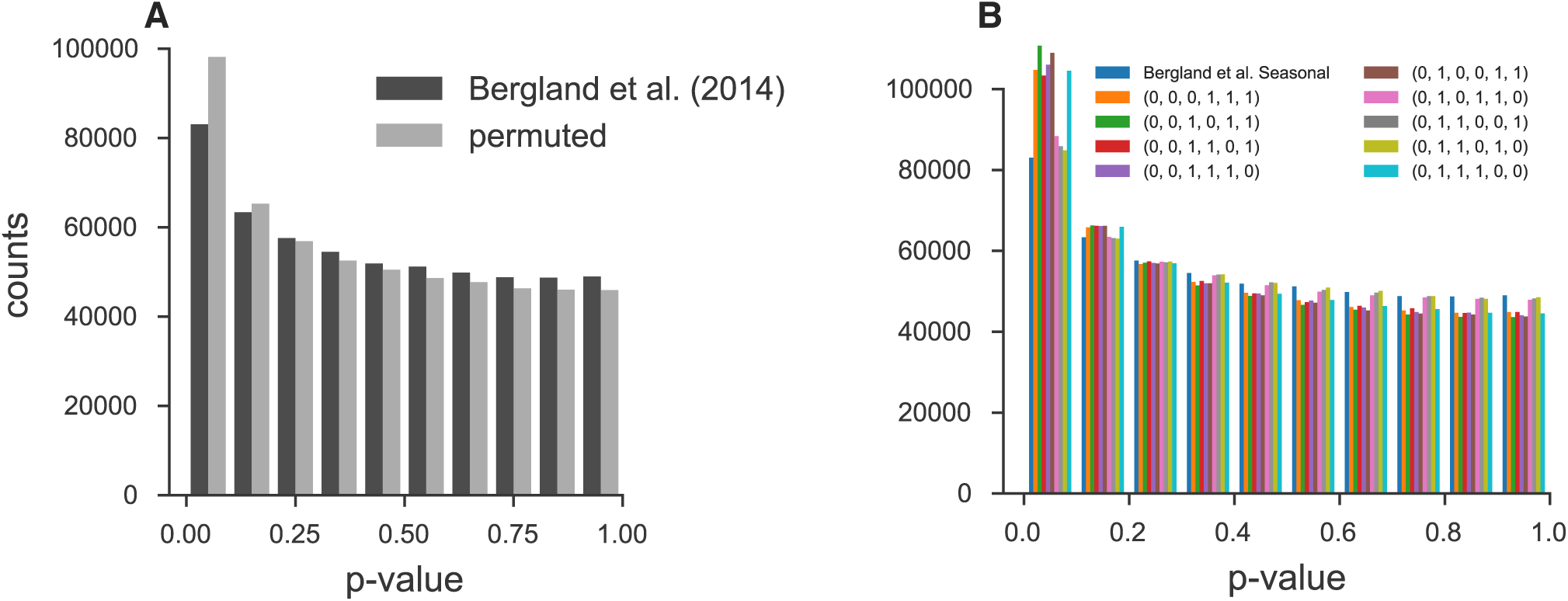
**A**: Histogram of original Bergland et al. (2014) seasonal p-values and p-values creating by randomly permuting the seasons at each locus. **B**: Histogram of original Bergland et al. (2014) p-values alongside all unique permutations (ignoring symmetries that lead to identical p-values).

To investigate whether there is a genome-wide evidence of an enrichment of fluctuating selection we created an empirical null distribution by randomly permuting the season labels and re-running the per-locus seasonal GLM model, as proposed by Machado et al. (2018). We find, regardless of whether we permute at the locus-level or the permutation replicate-level, that the observed seasonal p-value distribution Bergland et al. (2014) is not enriched for significant p-values beyond what we would expect from the permutation null. In fact, there appears there is more enrichment for low p-values when seasonal labels are randomly permuted (Supplementary Figure S3, suggesting by random chance we might expect more variants with a seasonal fluctuating pattern than found in the original Bergland et al. (2014) study. While surprising, this could be explained by the presence of temporal structure across the samples not consistent with seasonal fluctuating selection. Some fraction of the permutations happen to fit non-seasonal structure well, leading to an enrichment of small p-values. We note that genetic drift is not accounted for in the model used to estimate the significance of seasonally fluctuations and so some of these issues from non-seasonal structure may be due to a poorly calibrated null model. Furthermore, non-seasonal temporal structure is also evident in our temporal covariances (Supplementary Figure S1), where we see strong evidence of selection (non-zero temporal covariances), yet the pattern does not follow that of seasonal fluctuating selection.

**Figure S4:**
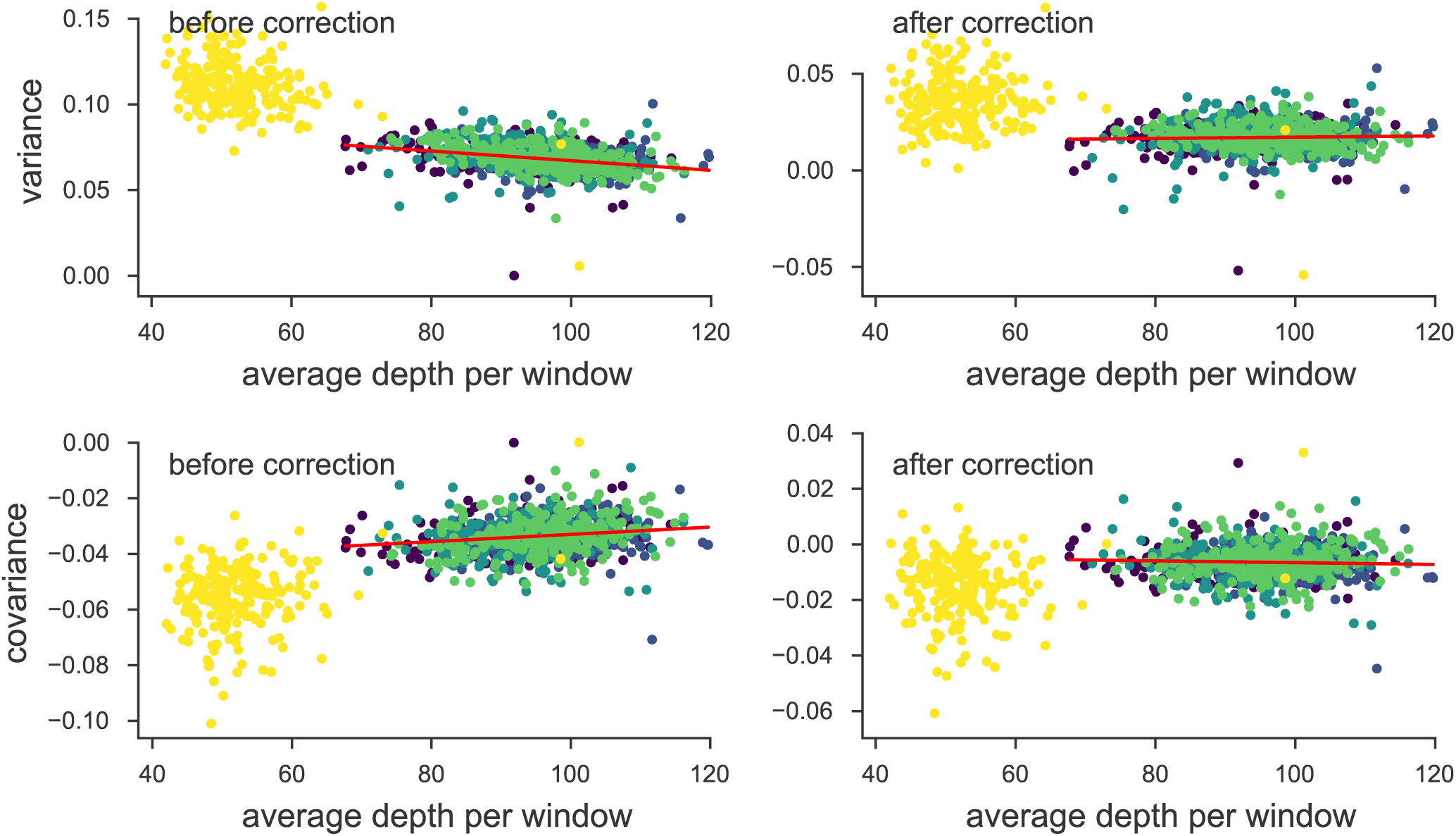
The variance and covariances from the Bergland et al. (2014) study, calculated in 100kb genomic windows plotted against average depth in a window before and after bias correction. Each panel has a least-squares estimate between the variance and covariance, and the average depth. The bias correction procedure is correcting sampling bias in both the variance and covariance such that the relationship with depth is constant. Colors indicate the different chromosomes of *D. melanogaster*; we have excluded the X chromosome (yellow points; chromosome 4 was not in the original study) from the regression due to large differences in average coverage.

### S7 Approximating the Reduction in Diversity from *G*(*t*)

To help reconcile our measure of linked selection, *G*(*t*), with familiar expressions as a reduction in neutral diversity, as parameterized by *N*_*e*_, we develop some rough approximations here. Note, however, that since linked selection generates temporal covariance, the overall effect is qualitatively different than just a simple reduction in *N*_*e*_, as drift alone cannot generate temporal covariances. First, we introduce some convenient notation. Let *V*_*T*_ = Var(*p*_*t*_ *–p*_*0*_) be the total observed variance in allele frequency, 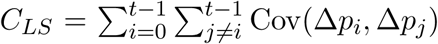 be the contribution of all pairwise temporal covariances (observable), *V*_*D*_ be the unobservable drift-only variance in allele frequency, and 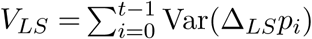 which is the (unobservable) effect linked selection has on the variances in allele frequency change. Then,

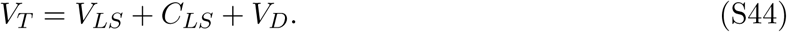

Our measure *G*(*t*) is then,

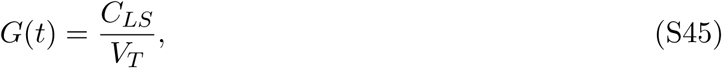

meaning we can express the fraction of total variance due to drift alone as

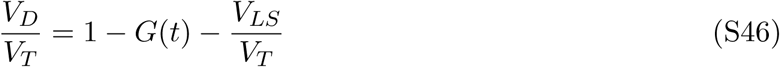

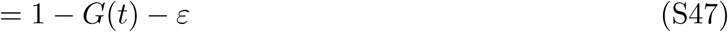

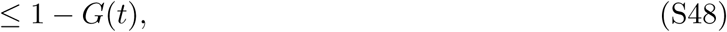

where *ε ≥* 0 since these variances are positive. Throughout this section, for convenience, we assume that the covariances contributing to *C*_*LS*_, and thus *G*(*t*), are all positive.

Rather than expressing this relationship in terms of variances in allele frequencies, we can express the same relationship in terms of *N*_*e*_. In a neutral Wright–Fisher population, the total variation in allele frequency change over *t* generations is

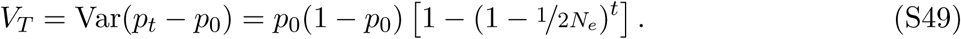

For small *t*, a common temporal estimator for the variance effective p opulation size *N*_*e*_ is,

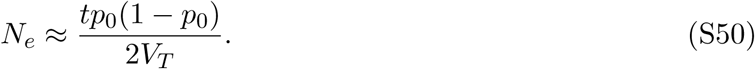

Then, the drift-only *N*_*e*_ estimate (that is, removing the effects of linked selection) replaces the observable *V*_*T*_ with unobservable *V*_*D*_, and uses the *G*(*t*) estimate to bound this:

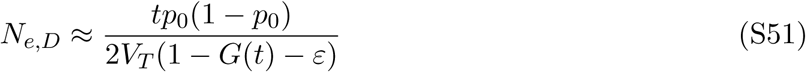

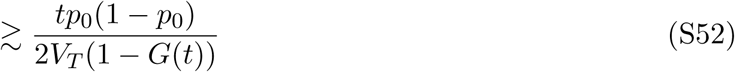

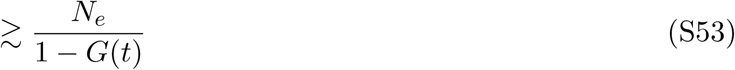

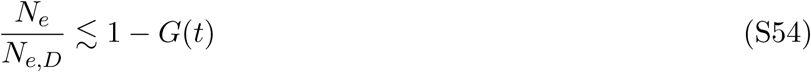

In the linked selection literature, it is common to translate the impact of linked selection as a reduction in the level of neutral pairwise diversity in the absence of linked selection, *π*_0_. Under the coalescent, *π*_0_ = 2*µ*𝔼(*T*_2_) where 𝔼 (*T*_2_) is the pairwise time to coalescence, which under drift alone in a constant population of size *N*_*e*_, is 𝔼 (*T*_2_) = 2*N*. The fraction of neutral diversity (in the absence of linked selection) reduced by selection is then, 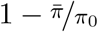, or equivalently, 1 − *N*_*e*_*/N*_*e,D*_. Then,

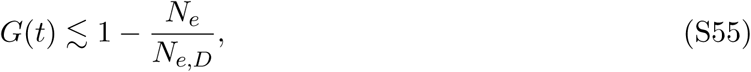

which shows that our measure *G*(*t*) is a lower bound, over much shorter timescales, to the familiar measure 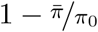.

Elyashiv et al. (2016) estimated that linked selection in *Drosophila melanogaster* had resulted in a 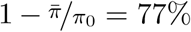 reduction in the level of neutral diversity. Thus our estimate of *G*(*t*) ≈ 20% in *Drosophila simulans* is smaller than that seen over long timespans in a closely related species.

### S8 Simulation Results

We conducted extensive simulations to understand how temporal covariance, *G*(*t*), and convergence correlations behave under (1) different quantitative genetic fitness models, (2) different trait architectures (e.g. varying levels of *V*_*A*_ for fitness and the number of sites affecting fitness), (3) background selection, and (4) different sampling periods. Furthermore, we use two replicate population simulations to investigate how convergence correlations depend on (1) the population sizes of each selection line sampled from the main population, and (2) the direction the trait is selected on in each line (i.e. in the same direction, differing directions, or only one lines elected).

Due to the high computational burden of forward simulations over this wide breadth of parameters, we modeled a single 50 megabase region in a population of *N* = 1000 diploid individuals with a neutral variant mutation rate of 10^−8^ and a recombination rate of 10^−8^ per basepair. This is roughly analogous to a quarter of an autosome of *Drosophila melanogaster*; however with this small population size and mutation rate, the population mutation rate *θ* for the entire region leads to far fewer neutral sites to calculate covariances and other statistics on than expected for a region this length in *D. melanogaster*. Since our main goal is to understand the dynamics of statistics used in the paper and how they are affected by different quantitative genetic fitness models, background selection, and trait architecture, we use population frequencies rather than sampling frequencies.

All forward simulations were conducted using SLiM (Haller and Messer 2019) and run and processed using Snakemake (Köster and Rahmann 2012); all simulation routines are available in the Github repository https://github.com/vsbuffalo/cvtk/.

#### S8.1 The Effects of the Genetic Architecture under Exponential Directional Selection

We first investigated the effects of the selected trait’s genetic architecture on temporal covariances and *G*(*t*) by neutrally burning in a population for 10*N* generations, and selecting on the trait with an exponential fitness function (where fitness of a trait value *z* is *w*(*z*) ∝ exp(*z*)). The exponential fitness function corresponds to multiplicative selection across sites and so serves as the simplest directional selection model of a trait to understand the effects of genetic architecture on the statistics we have used in the paper.

**Figure S5:**
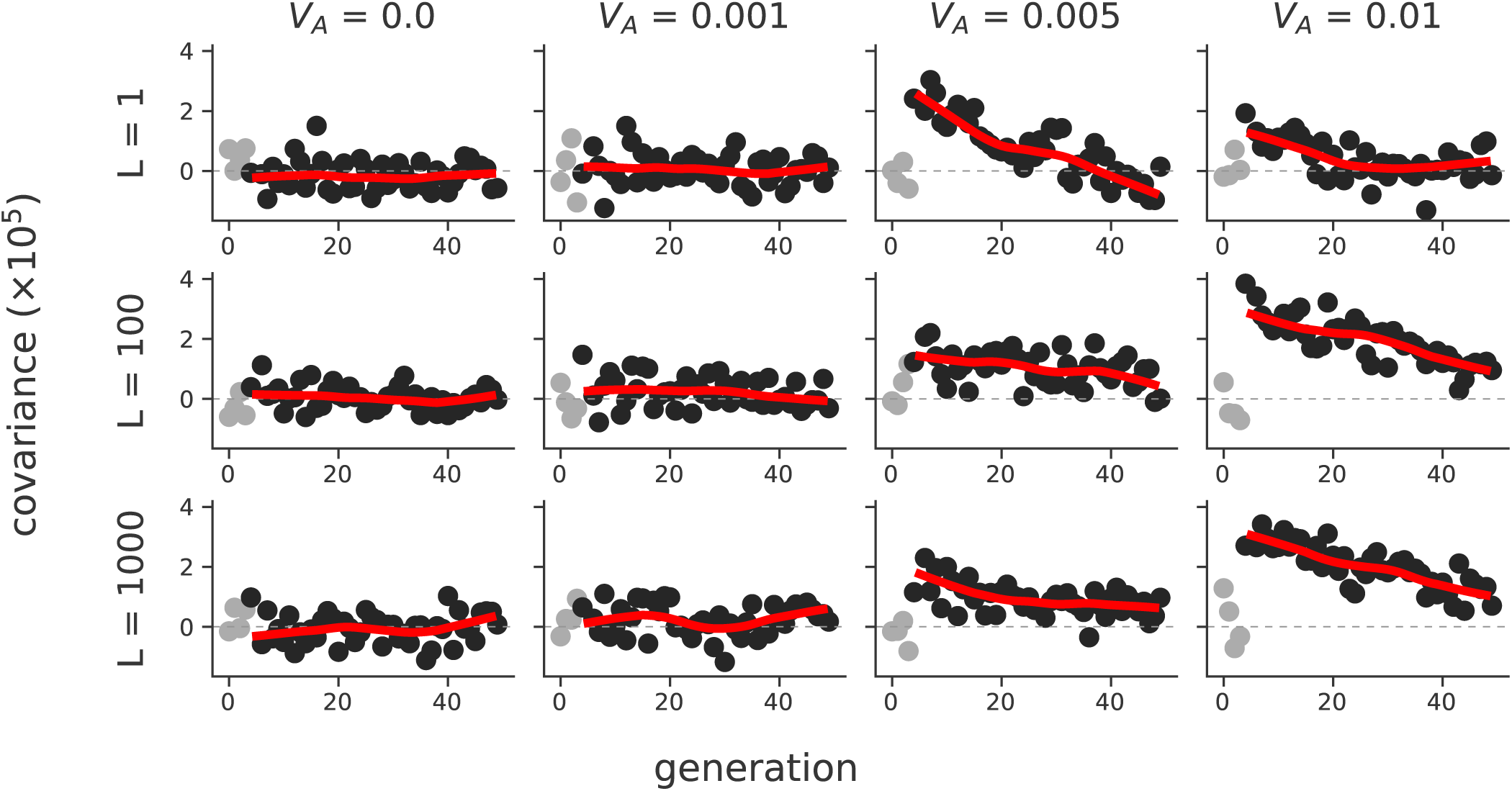
The temporal covariances Cov(Δ*p*_5_, Δ*p*_*t*_) from the onset of selection (generation 5) to a later time point *t*, which varies along the x-axis, across a variety of different initial trait additive genetic variances (*V*_*A*_, columns) and number of sites contributing to the trait (*L*, rows). Each point is the temporal covariance averaged over 50 replicate simulations; dark gray points are temporal covariances after the onset of selection, and light gray points are before. The red line is a loess-smoothed curve through the covariances after the onset of selection. Selection on the trait was imposed through an exponential fitness function.

During this burnin, sites were either marked as neutral (with mutation rate *µ*_neutral_ = 10^−8^ per gamete per generation) or contributed to the trait’s value (with mutation rate *µ*_trait_), but were not selected until generation 10*N* + 5 (the five generations after burnin serve as a neutral control). The trait mutation rate, *µ*_trait_ was set by targeting a particular architecture, the number of selected sites, *L*. Each site contributing to the trait’s value was randomly chosen to have effect size *± α* with equal probability, where *α* was set to target a particular additive genetic variance for the trait, *V*_*A*_, for the target number of selected sites *L*.

Overall, we again see a finding of Buffalo and Coop (2019): that the initial expected temporal covariance conditioned on *V*_*A*_, is invariant to the number of loci determining the trait’s value, *L* (Supplementary Figure S5). We do find some evidence that the decay in temporal covariance is faster when the trait has a monogenic basis (see the third column of Supplementary Figure S5); this is expected since the selection coefficients are larger for these monogenic simulations, leading to faster allele frequency changes and a rapid change in additive genetic variance.

In our previous work, we did not investigate the affect of trait architecture on our measure *G*(*t*). Using the exponential fitness function simulations, we also calculated *G*(*t*) for each of the replicate simulations. We find that the *G*(*t*) trajectories can vary considerably across replicates depending on the number of sites (*L*) determining the trait’s value (Supplementary Figure S6). When a trait is reasonably monogenic (*L* ≈ 1), *G*(*t*) trajectories vary considerably across replicate lines, as certain lines may stochastically lose the few copies of the selected alleles (top row of Supplementary Figure S6). However, with a polygenic trait, (*L ≥* 100), the *G*(*t*) trajectories across replicates are similar as each replicate contains an abundance of trait alleles (bottom rows of Supplementary Figure S6). Comparing the simulated *G*(*t*) replicate trajectories of Supplementary Figure S6 with the Barghi et al. (2019) *G*(*t*) trajectories in Figure 1B, we again confirm a finding of Barghi et al. (2019): that there is considerable genetic redundancy among beneficial alleles, meaning because of the polygenic architecture, there are multiple routes to adaptation. We should note that our simplified simulation routines are slightly different from the Barghi et al. (2019) study in that the burnin populations are all independent; however we expect the same qualitative result.

**Figure S6:**
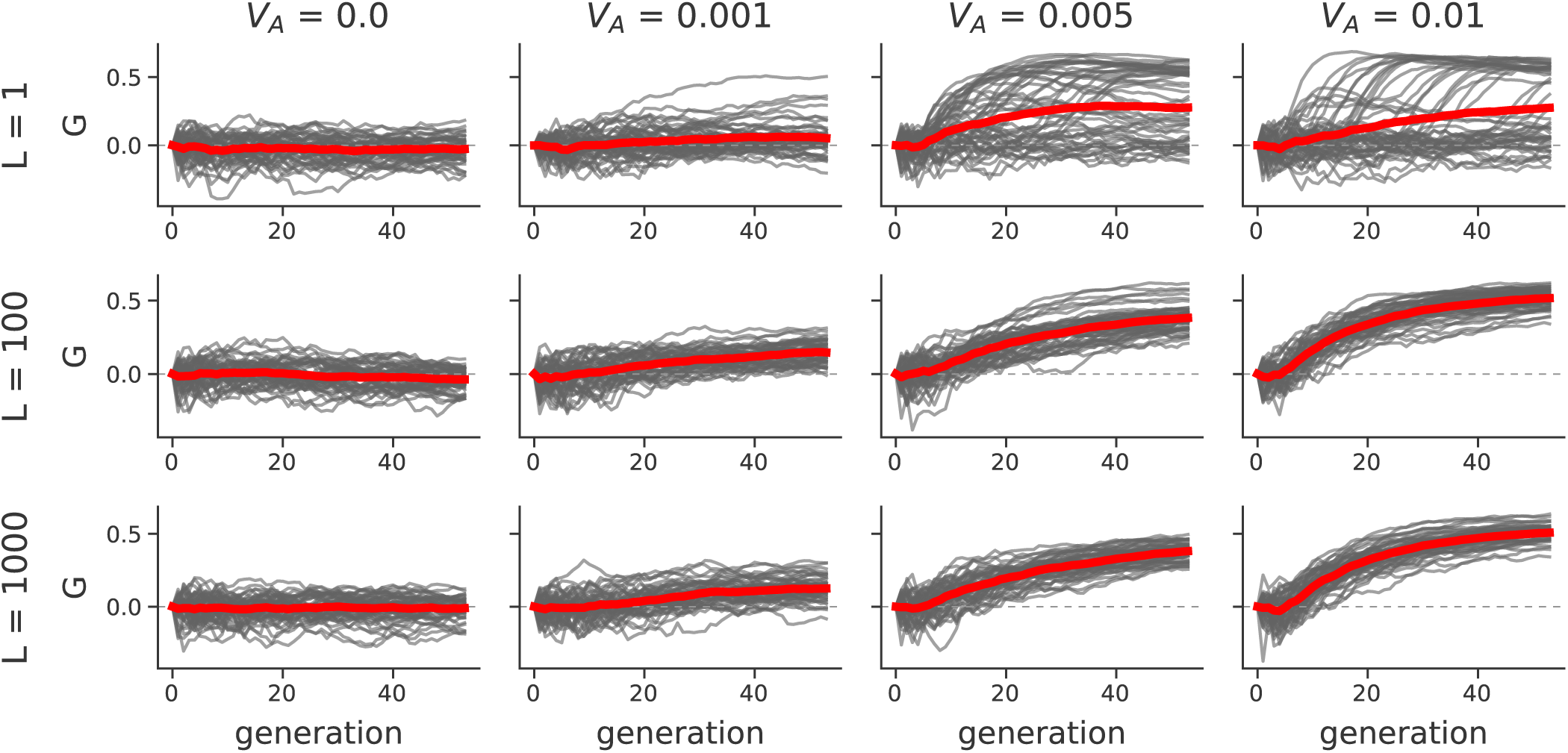
The *G*(*t*) trajectories of 50 replicate simulations, across different trait architectures (*L* is the target number of sites affecting the trait’s value, and *V*_*A*_ is the target trait additive genetic variance). The red line is the mean trajectory across all replicate simulations. Like Supplementary Figure S5, the onset of selection is five generations after the 10*N* generation burnin; this is evident by the initial flat period of the *G*(*t*) trajectory.

#### S8.2 Temporal Covariances and *G*(*t*) under Gaussian Stabilizing Selection

Additionally, we wanted to ensure that our temporal covariances and *G*(*t*) trajectories were robust to more complicated, but realistic fitness models. To this end, we also simulated Gaussian stabilizing selection (GSS) on a trait during burnin, followed by one of two optimum shift routines: (1) sudden optima shifts of *µ*_sudden_ = {0.1, 0.5, 1}, and (2) very graduate optima shifts of *µ*_gradual_ = {0.001, 0.01} per generation using the same two population simulation scheme described above (these shifts are in standard deviations, since *V*_*S*_ = 1). We used a polygenic architecture for these simulations, with trait alleles assigned a *±* 0.01 effect size with equal probability, trait mutation rate 10^−8^, and the optima shift began at five generations after a 10*N* generation burnin. Across our GSS simulations, we see the expected selection response (Supplementary Material Figure S7).

Overall, we see the same qualitative results under Gaussian stabilizing selection with optima shifts as under exponential directional selection. Stronger directional selection, here determined by larger sudden optima shifts or larger gradual shifts per generation, lead to stronger temporal covariances (Supplementary Materials Figure S8). Furthermore, we see a stronger effect of linked selection, as measured by *G*(*t*), under stronger directional selection (Supplementary Material Figure S9).

**Figure S7:**
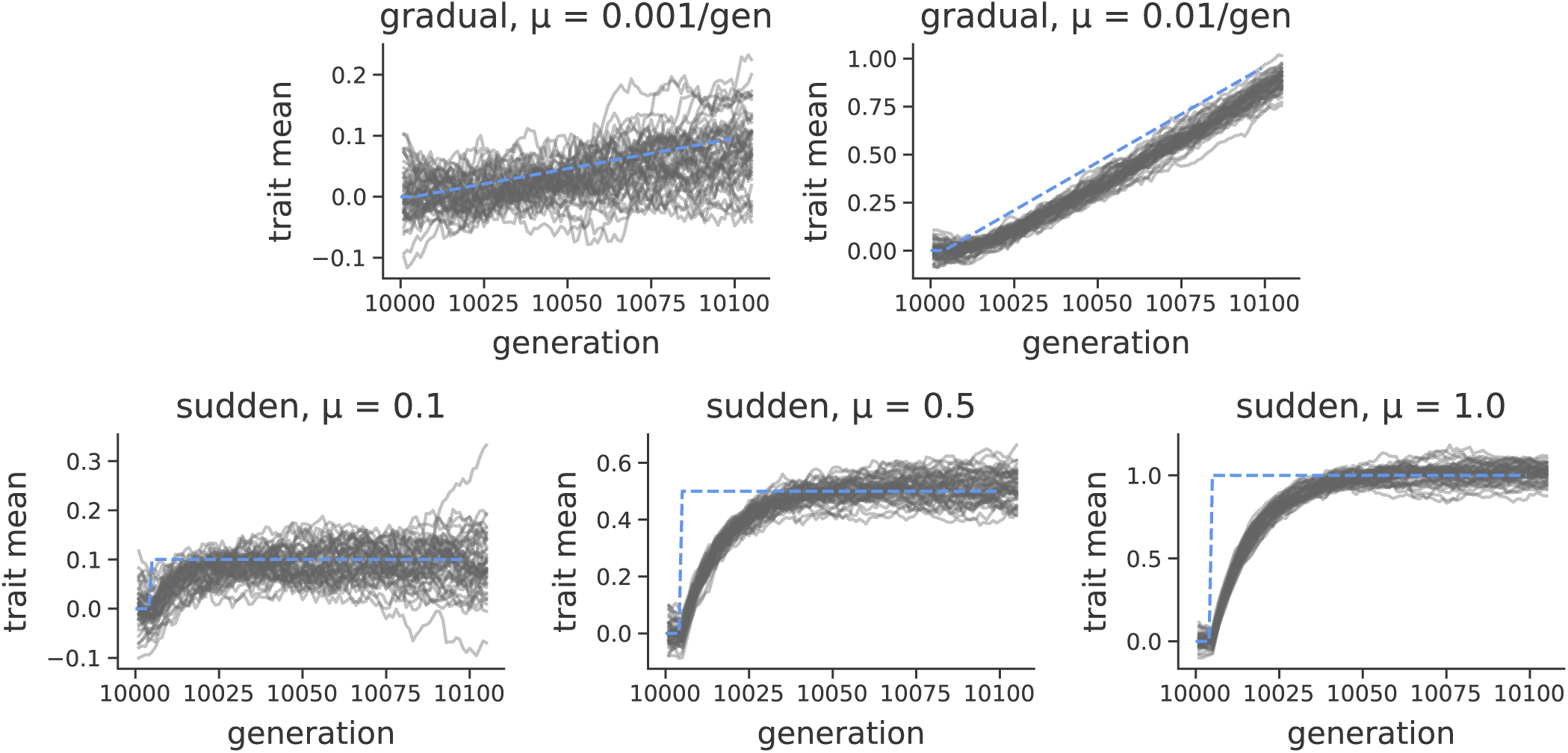
The population mean trait value under the Gaussian stabilizing selection simulations (gray lines) and the trait optima (dashed blue lines). The first row shows the selection response during a graduate shift in optima per generation, while the second row shows the selection response during a sudden optima shift.

**Figure S8:**
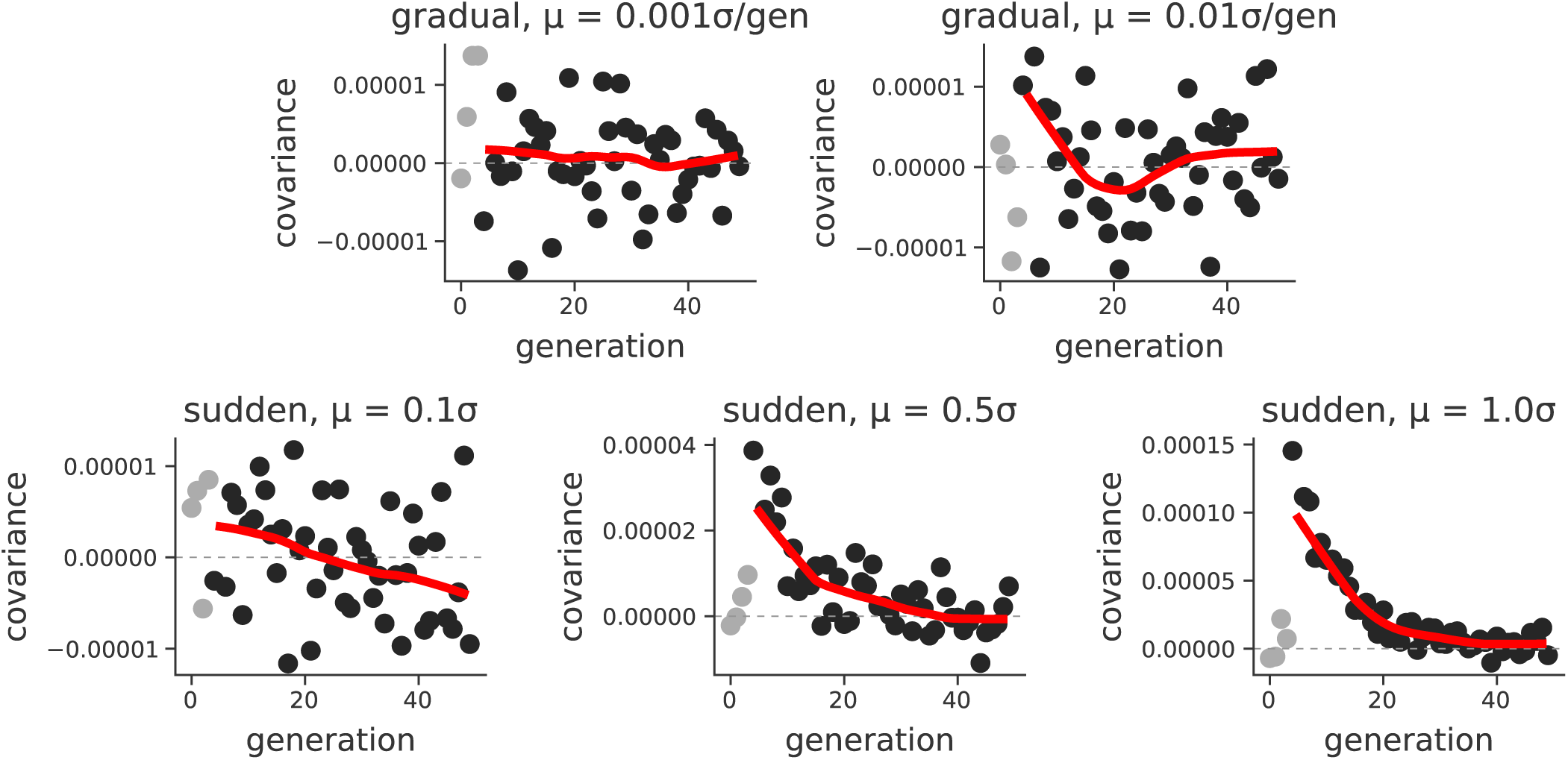
Mean temporal covariance (Cov(Δ*p*_5_, Δ*p*_*t*_), with *t* varying across the x-axis) across 30 replicate simulations (light gray points are before the onset of selection; dark gray points are after selection begins), under different Gaussian stabilizing selection with optima shift regimes. The solid red line is a loess-smoothed average of these points.

**Figure S9:**
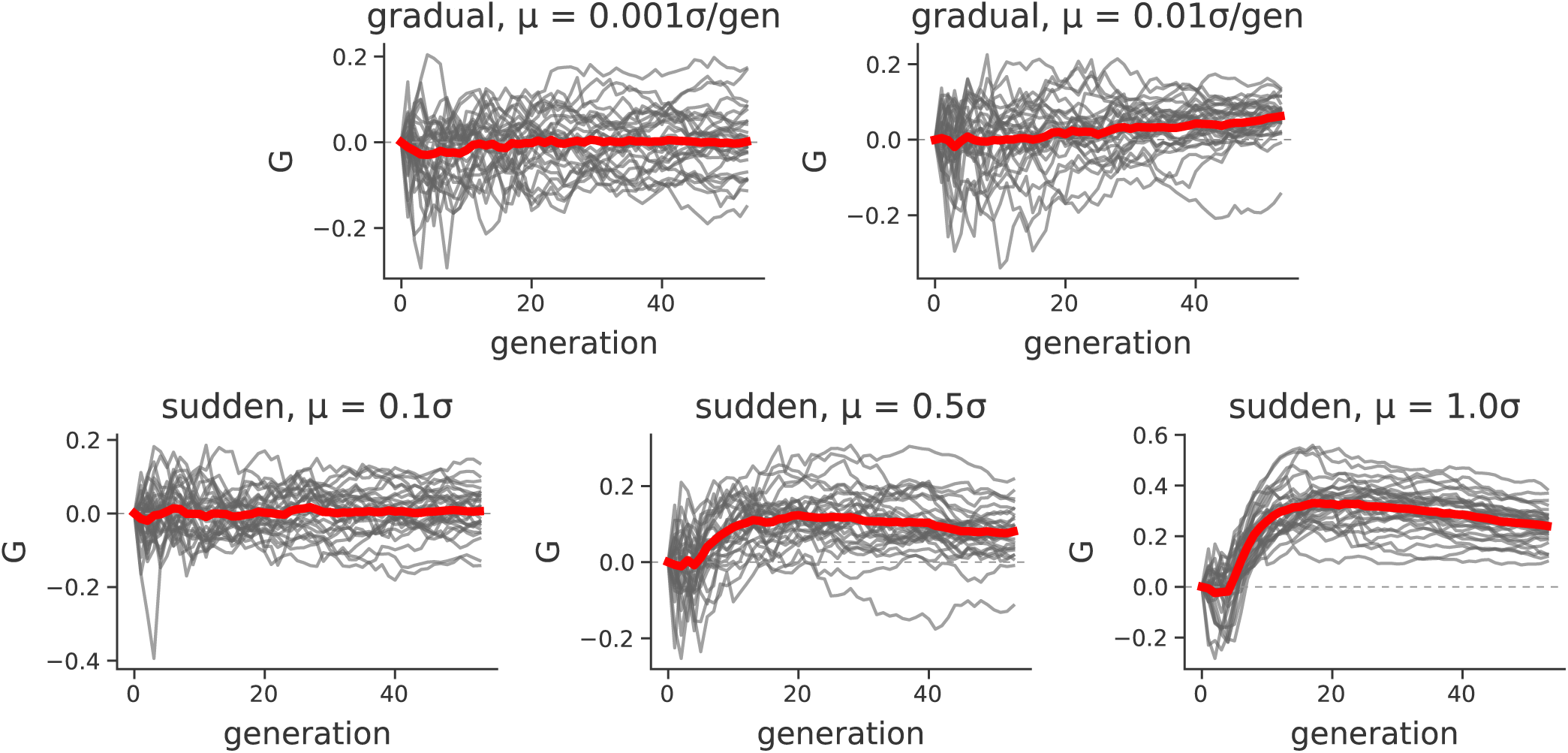
*G*(*t*) trajectories across 30 replicate Gaussian stabilizing selection with optima shift regimes. The solid red line is a loess-smoothed average across replicates.

Additionally, we looked at the effect of the replicate population size drawn from the same population has on a single population’s *G*(*t*) trajectories. These simulations had the same 10*N* generation burnin, followed by a change in population size emulating the bottlenecks associated with creating selection lines. Overall, we find that smaller population sizes lead to a reduced *G*(*t*) (Supplementary Material Figure S10). This is expected, as the denominator of *G*(*t*) is Var(*p*_*t*_ − *p*_0_), which has an inverse relationship with *N*_*e*_; as replicate population size is reduced, the proportion of allele frequency change driven by linked selection is lower, since the rate of drift is increased. To isolate the effects of varying replicate population size, we also looked at just the magnitude of temporal covariances (Supplementary Material Figure S11). We find that smaller replicate population sizes lead to *larger* temporal covariances. We then looked at the initial trait variance, which as expected, does not vary with replicate population size (since all burnin populations had the same size). This implies that the linkage disequilibria is higher in smaller populations, due to founder effects, which has the effect inflating the temporal covariance as predicted by our theory (Buffalo and Coop 2019).

#### S8.3 Convergence Correlations

Using the same exponential fitness function simulations described above, we also investigated how the convergence correlation is impacted by (1) genetic architecture, (2) the design of the selection experiment, e.g. how many individuals are selected for each line from the founding population, and (3) the direction of selection across the two populations “lines”. After burning in *N* = 1000 diploid populations for 10*N* generations, we simulated two equally-sized lines of sizes *n* = {50, 500, 1000} diploids, and imposed three selection schemes across different simulation runs. First, we imposed a convergent selection scheme, where the populations undergo exponential directional selection in the same direction. We expect that the convergent correlation under this convergent scheme should be positive, as the two lines should share some haplotypes carrying beneficial alleles, and these are selected in the same direction across the two lines. Second, we imposed divergent selection, where the two lines again undergo exponential directional selection, except in different directions. Here, we expect the convergence correlation to be negative, as haplotypes that increase the selected trait in one population are beneficial in the upward selected line, but deleterious in the downward selected line. Third, we have a control selection scheme, where one line is selected and the other is not; this is akin to the control line in the Castro et al. (2019) study (see Figure 2C). In this case, we expect to see no convergence correlation, as only one line is being selected. Finally, across these two-line simulation studies, we expect that smaller selection line sizes should show weaker convergent correlations, as the probability that the same haplotypes are selected between the two lines decreases with size.

**Figure S10:**
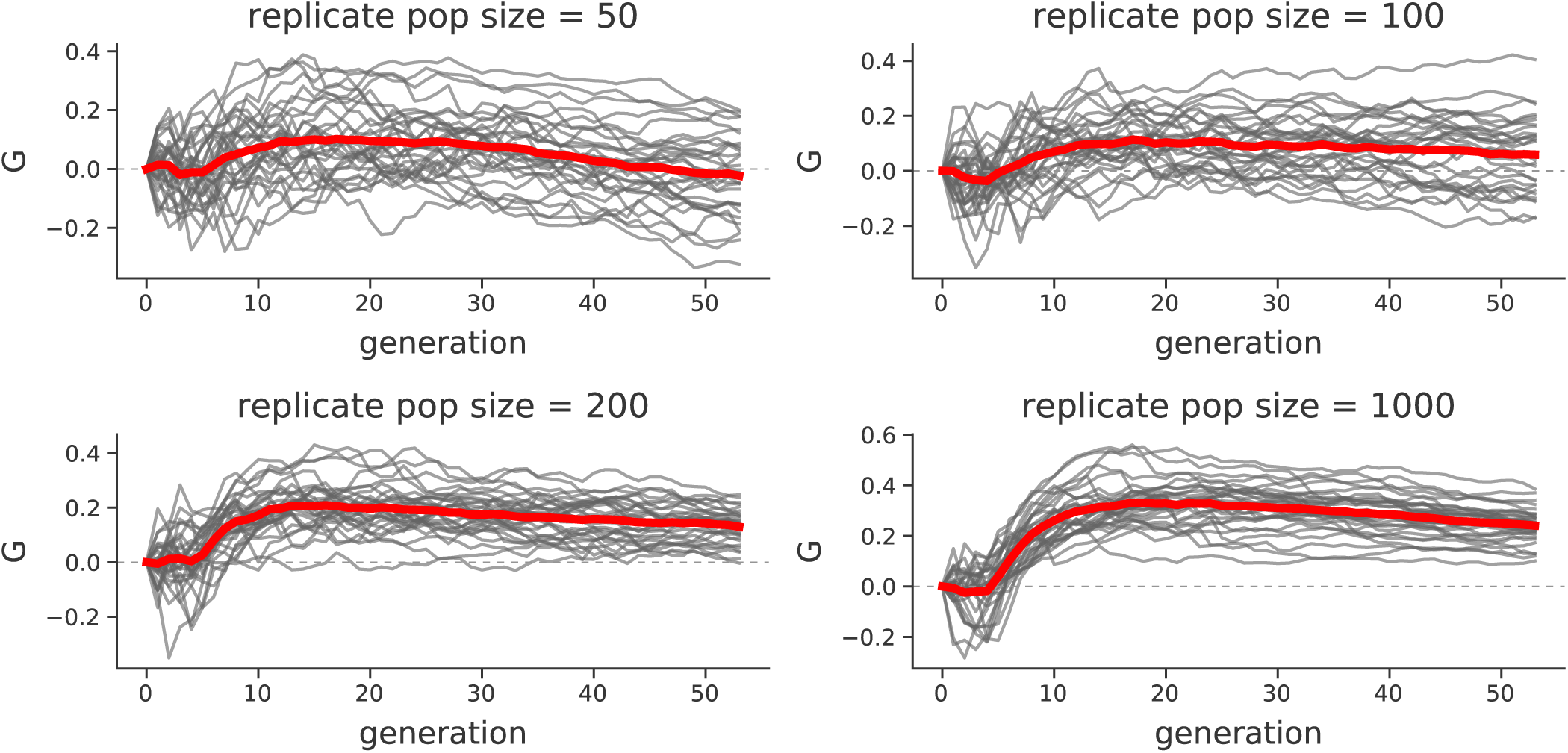
*G*(*t*) trajectories under GSS after sudden optima shift of 1 at generation five, for varying replicate population sizes.

**Figure S11:**
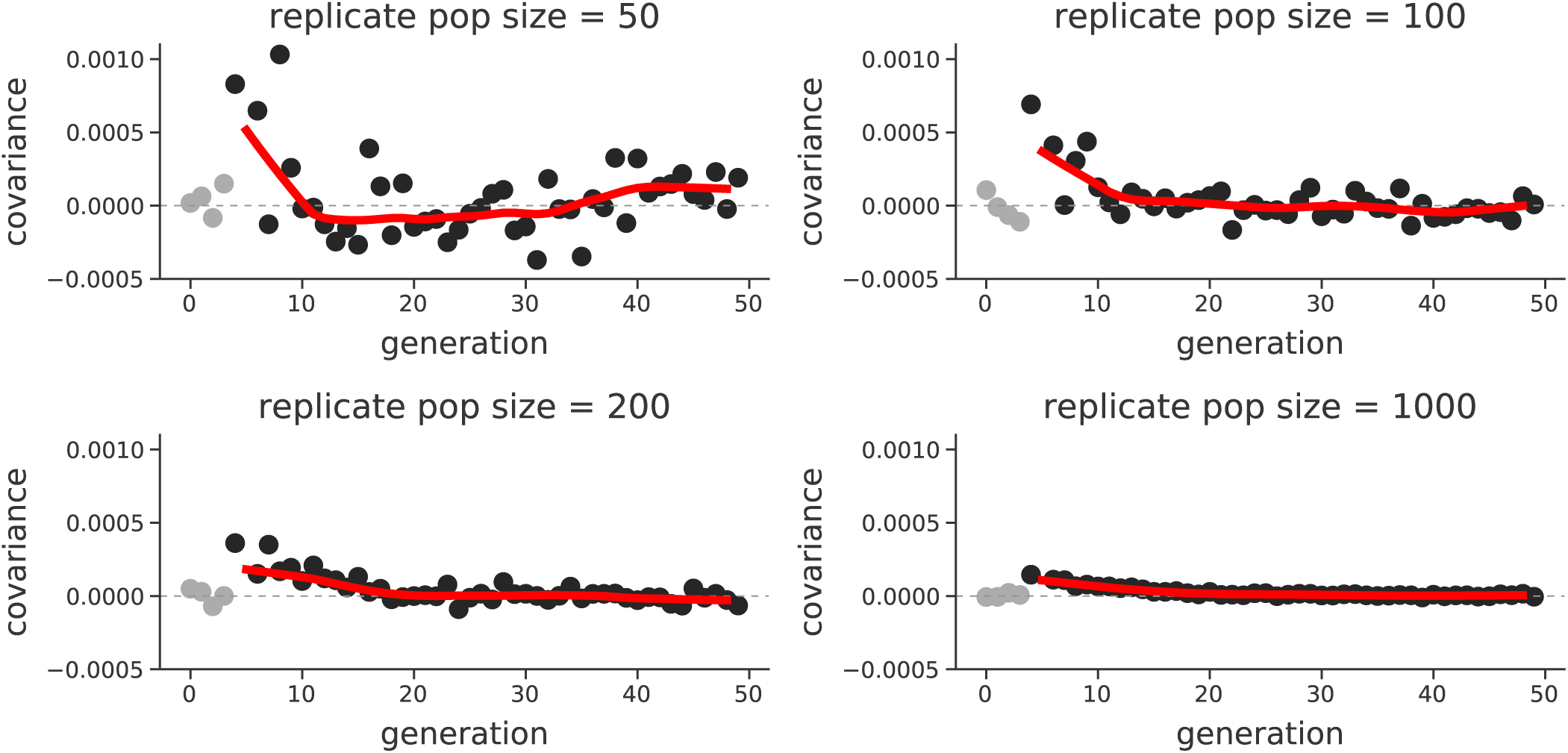
Temporal covariance Cov(Δ*p*_5_, Δ*p*_*t*_), where *t* varies along the x-axis, for a sudden optima shift of 1, for varying replicate population sizes. The reference time point is the first generation of selection; dark gray points are the temporal covariance after selection began, and the light gray points are before.

**Figure S12:**
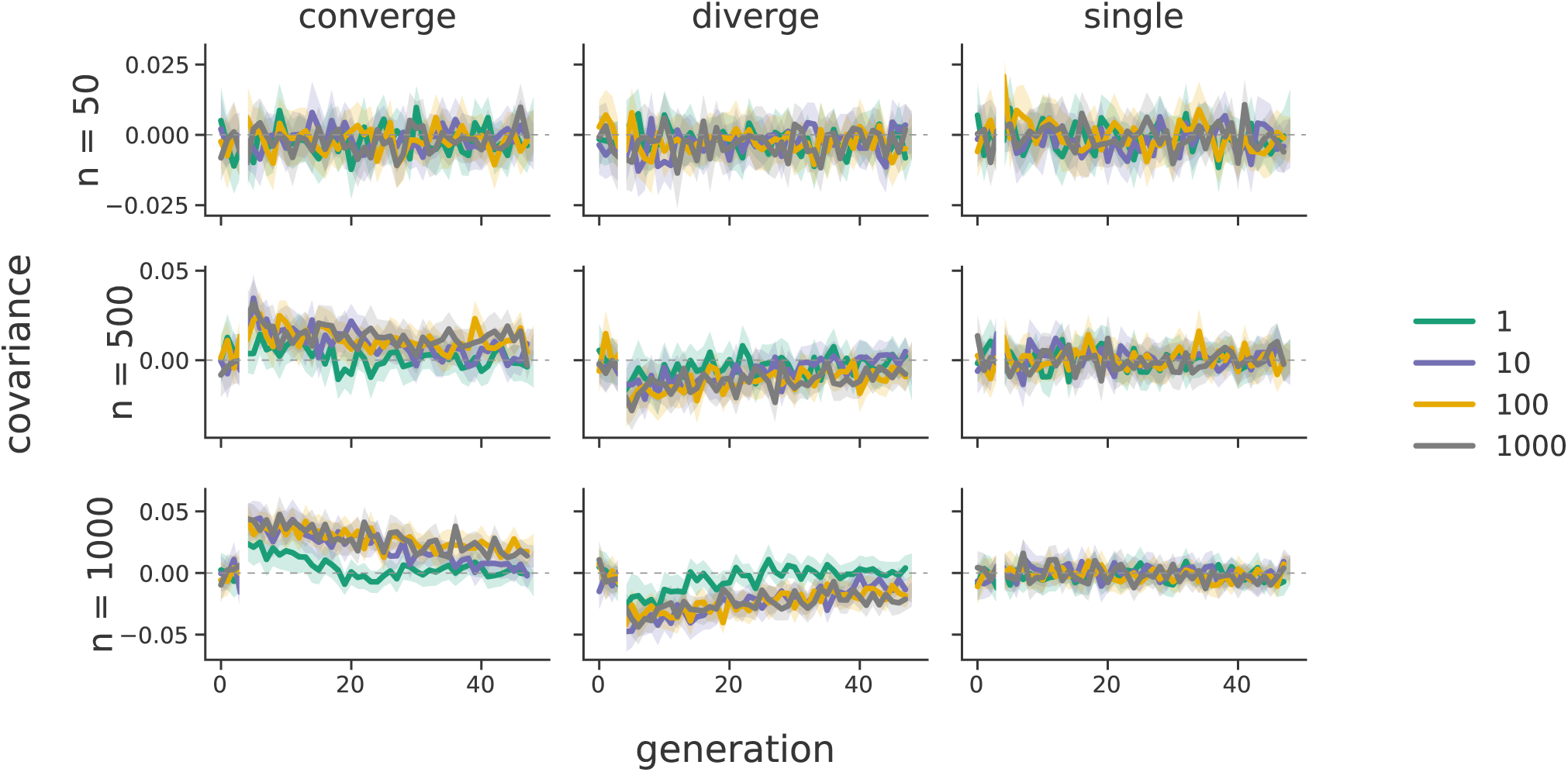
The convergence correlations across the two population line exponential directional selection simulations; panel rows are for differing line population sizes, and panel columns are the modes of selection across the lines (convergent, divergent, and only a single selected line control). Line color indicates the target genetic architecture, in number of loci affecting the trait’s value. 95% confidence intervals are also shown. Note that selection begins at generation five, which is the reference generation; this is indicated by the split in the lines.

Overall, our simulations confirm our hypotheses; see Supplementary Material Figure S12. We also find that in simulations with a monogenic genetic architecture (i.e. the target number of trait-affecting loci is *L* = 1), the convergence correlations are generally much weaker than those under a polygenic architecture. However, this effect is mediated by the line population size; the difference in convergence correlation between *L* = 1 and *L* = 1000 are more dissimilar when the line population sizes are larger (compare the first column, last two rows). Like the convergence correlations calculated on the Barghi et al. (2019) data, we find in simulations convergence correlations decay through time. Additionally, populations selected in opposite directions lead to negative convergence correlations, as expected. Overall, we find that the convergence correlation is affected by both genetic architecture and the size of the selected population lines.

We also wanted to test whether we see similar convergence correlations under Gaussian stabilizing selection. In these simulations, rather than targeting a particular *V*_*A*_, we fix the trait mutation rate at 10^−8^ (thus region-wide *θ* = 2000). Like the exponential directional selection simulations, we impose directional selection in the same direction across the two populations (converge), different directions (diverge), and only in one population (single). We also vary the type (gradual versus sudden) and magnitude of optima shifts in the two populations. Overall, simulations show convergence correlations for the sudden optima shifts in Supplementary Figure S13. Again, optima shifts in the opposite direction cause negative convergence correlations. We found that for slow moving optima shifts, the convergence correlations are generally too weak to be distinguished from zero reliably (not shown).

**Figure S13:**
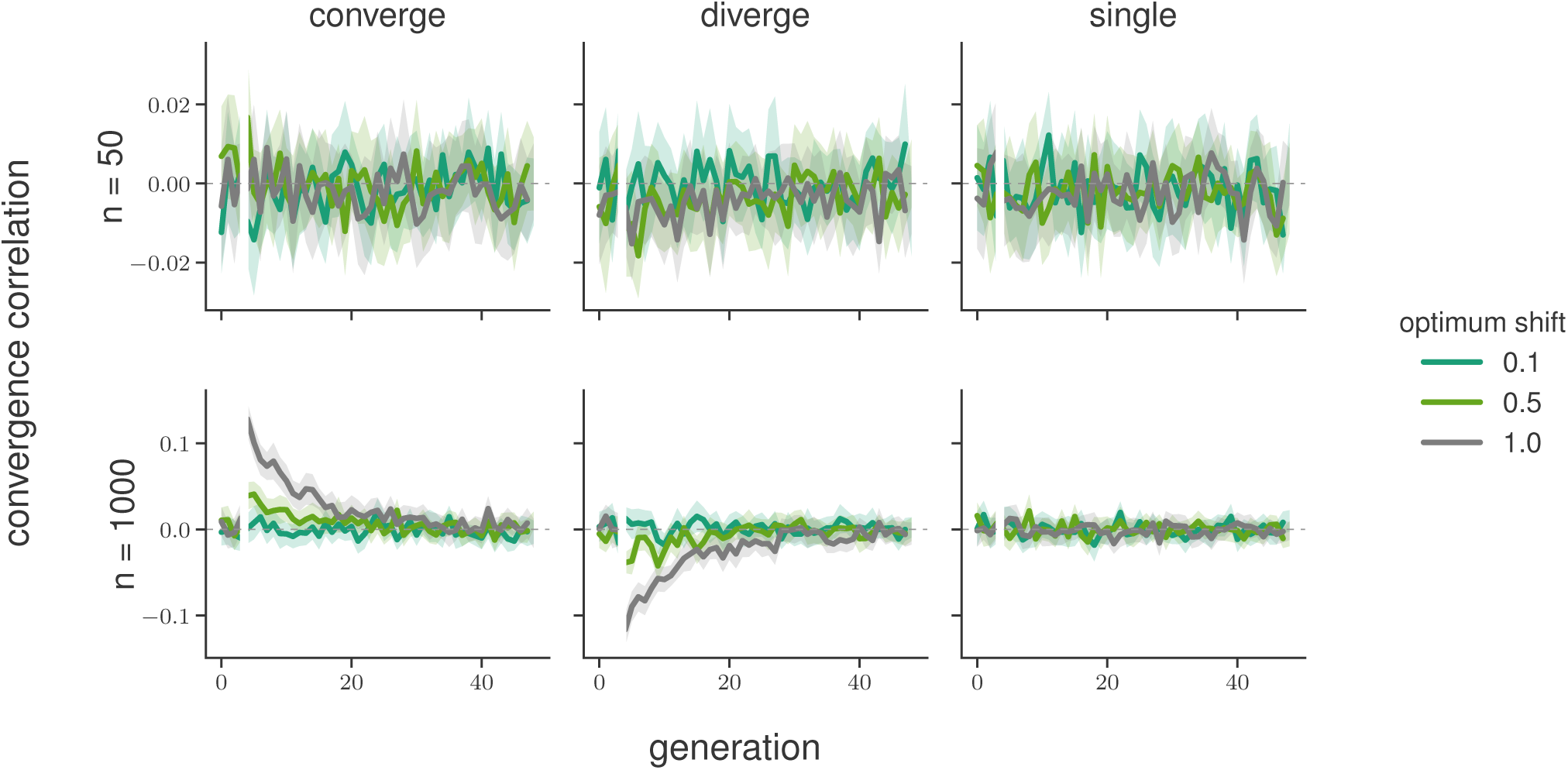
The convergence correlations across the two population line Gaussian stabilizing selection sudden optima shift simulations; selection line population sizes vary across rows, and panel columns are the modes of selection across the lines (convergent, divergent, and only a single selected line control). All simulations have a target number of loci affecting the trait of *L* = 1000; line color indicates the size of the sudden optima shift in standard deviations of *V*_*S*_ 95% confidence intervals are also shown. Note that selection begins at generation five, which is the reference generation; this is indicated by the split in the lines.

#### S8.4 Sampling in Temporal Blocks

In our analysis of the Barghi et al. (2019) data, we describe our statistic *G*(*t*) as a lower bound for two reasons: (1) the population is sequenced every ten generations, meaning the temporal covariances between adjacent generations cannot contribute to the numerator of *G*(*t*) but contributes to the denominator, and (2) the estimate of *G*(*t*) ignores linked selection’s contribution to the per-generation variance in allele frequency change. In Buffalo and Coop (2019), we define an alternative estimator that includes selection’s effects on these variance terms,

**Figure S14:**
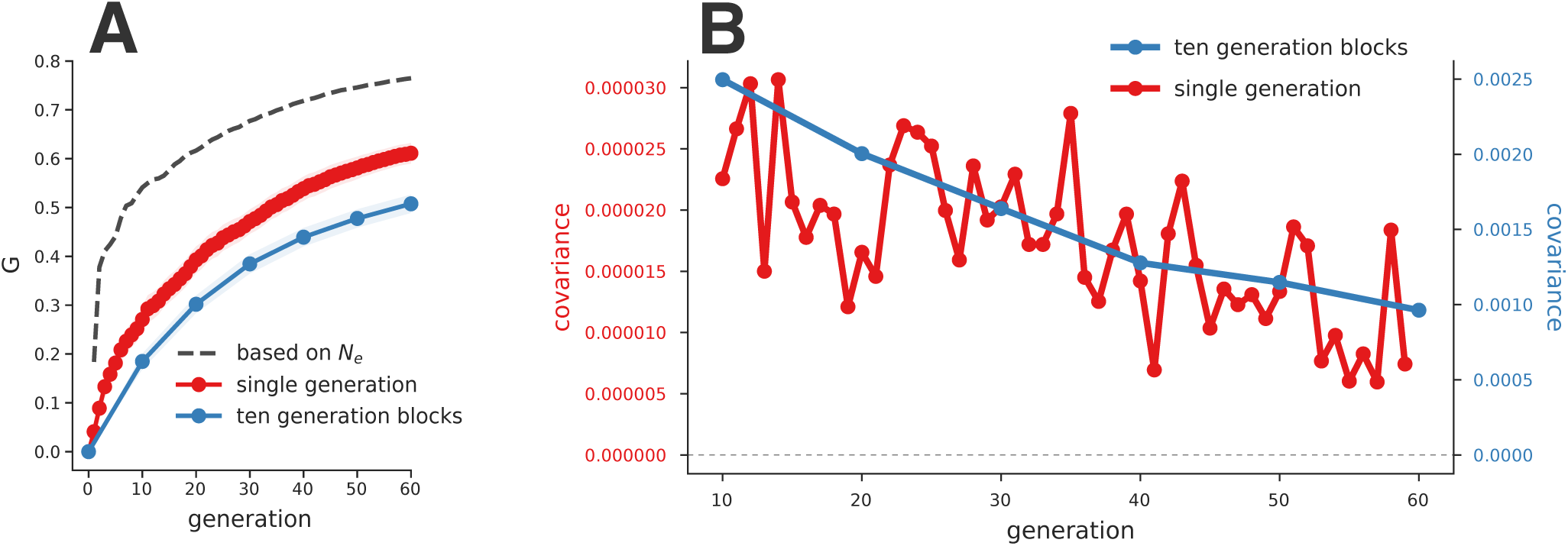
A: The *G*(*t*) averaged over 50 replicate simulations with *V*_*A*_ = 0.01 and *L* = 1000. The blue line shows *G*(*t*) calculated over ten generation blocks, similar to the calculation of temporal covariances of the Barghi et al. (2019) study. The red line shows the average *G*(*t*) estimates when the population is sampled every generation and all covariances can contribute to the numerator of *G*(*t*). The dashed dark gray line indicates the *G*^*′*^(*t*) estimate, which uses the known drift effective population size of the simulations. B: The temporal covariances calculated each generation (red line) and on ten generation blocks (blue line) using the same simulation data.

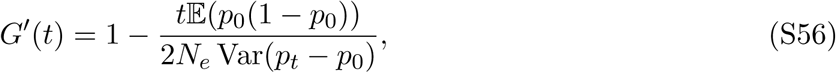

which can be calculated when the drift-effective population size *N*_*e*_ can be estimated (see equation 26 in Buffalo and Coop 2019 for details).

To verify that *G*(*t*) estimated every ten generations is indeed a lower bound, we used a simulation procedure similar to the exponential fitness function simulations (described in Supplementary Material Section S8.1), and calculated the temporal covariances and *G*(*t*) both each generation, and every ten generations. Unlike the simulations described in S8.1 where selection began at 10*N* + 5 generations, selection starts at generation 10*N* here, and used trait *V*_*A*_ = 0.01 and targeted *L* = 1000 sites affecting the trait.

First, comparing *G*(*t*) when sampling population frequencies every generation versus every ten generations, we confirm that the ten-generation block *G*(*t*) is a lower bound of the *G*(*t*) trajectory when sampling is every generation (red and blue lines in Supplementary Figure S14A). Furthermore, since we control the population size and reproductive sampling scheme in our simulations at *N* = 1000 diploids, we know the drift-effective population size in the absence of selection, which allows us to estimate *G*′(*t*). Plugging in the drift-effective population size *N*_*e*_ = 1000 into the expression for *G*′(*t*) and using the Var(*p*_*t*_ − *p*_0_) calculated for different *t*’s, we see that *G*(*t*) calculated every generation does not account for linked selection’s inflation of Var(Δ*p*_*t*_) and underestimates the true impact of linked selection as expected (dashed gray line in Supplementary Figure S14A).

To further understand the effects of calculating temporal covariances every ten generations rather than every generation, we also compared their magnitudes and decay rates using the simulations described above. We find that ten generation block temporal covariances are orders of magnitude larger but decay at similar rates (see Supplementary Figure S14B; note the two y-axis scales are different). The larger magnitude is expected, as each ten generation block temporal covariance is the sum of 45 temporal covariances between adjacent generations (e.g. 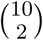).

#### S8.5 Background Selection

In our previous work, Buffalo and Coop (2019), we did not investigate whether background selection can lead to temporal autocovariance. Here, using forward-in-time simulations, we find that background selection can indeed generate temporal autocovariance and lead to convergence correlations when deleterious haplotypes are shared between populations and both are removed by selection.

We simulated background selection in a 50 megabase region, where deleterious alleles are randomly introduced by mutation. Following background selection literature (Charlesworth et al. 1993; Hudson and Kaplan 1995; Hudson and Kaplan 1994; Nordborg et al. 1996), we parameterize the mutation rate as the total number of deleterious mutations introduced per diploid genome, per generation, and simulate values *U* = {0.5, 1.0, 1.5}. Note that for our 50Mb region, our choice of BGS *U* parameters are on the higher end of the spectrum expected for *Drosophila*, but we wanted to ensure the strength was sufficient to see a signal. Note that if *U* ≈ 1.6 (Elyashiv et al. 2016), and the *Drosophila* genome is ≈ 140Mb, our region is ≈36% of the genome; this implies a reasonable *U* for our region is *U* ≈ 0.57, which is close to the bottom end of our parameter range. We also vary the strength of selection against the deleterious mutations, *s* = {0.01, 0.05, 0.1}, as well as different recombination rates (*r*_*bp*_ = {10^−7^, 10^−8^}). Like other simulations, we burnin the population for 10*N* generations under backgrounds selection. Overall, we find background selection does create temporal covariance (Supplementary Material Figure S16), which are stronger under (1) higher deleterious mutation rates and (2) larger selection coefficients. This latter point initially seems at odds with background selection theory, as the level of pairwise diversity in a region under strong background selection is invariant with respect to the selection coefficient. However, looking at the background selection *G*(*t*) trajectories, we find that over time, background selection appears to trend towards an asymptote in the *r*_*BP*_ = 10^−7^ subfigure, and reaches an equilibrium in the *r*_*BP*_ = 10^−8^ subfigure that seems reasonably invariant to the choice of *s* (Supplementary Material Figure S15). We believe that these observations can be reconciled by *Cov*(Δ*p*_*t*_, Δ*p*_*t*′_) being larger for larger *s* when |*t*′ − *t*| is small, but also decaying more rapidly with |*t*′ − *t*|, such that the overall contribution of selection to allele frequency variance is invariant to *s* (for strong background selection). However, further work is needed to fully explore and understand the temporal covariance dynamics of background selection.

Additionally, we investigated whether background selection can create convergence correlations between two replicate populations. Much like the exponential directional selection and Gaussian stabilizing selection simulations, we burned in a population for 10*N* generations with background selection, which continued after the population was split into two replicate populations. These simulations fixed *U* = 1.0, *r*_*BP*_ = 10^−8^, and varied the replicate population size *n* = {200, 1000}. We find that background selection can create convergence correlations (Supplementary Material Figure S17). We find the convergence correlation is weaker in smaller replicate population sizes, as there are fewer shared haplotypes carrying the same deleterious alleles between the two populations.

Finally, we found in processing these background selection simulation results that including or excluding sites fixed or lost through time can lead to differences in the estimated *G*(*t*) trajectories and temporal covariance. We discuss this extensively in a subsequent section, Supplementary Materials Section S8.7.

**Figure S15:**
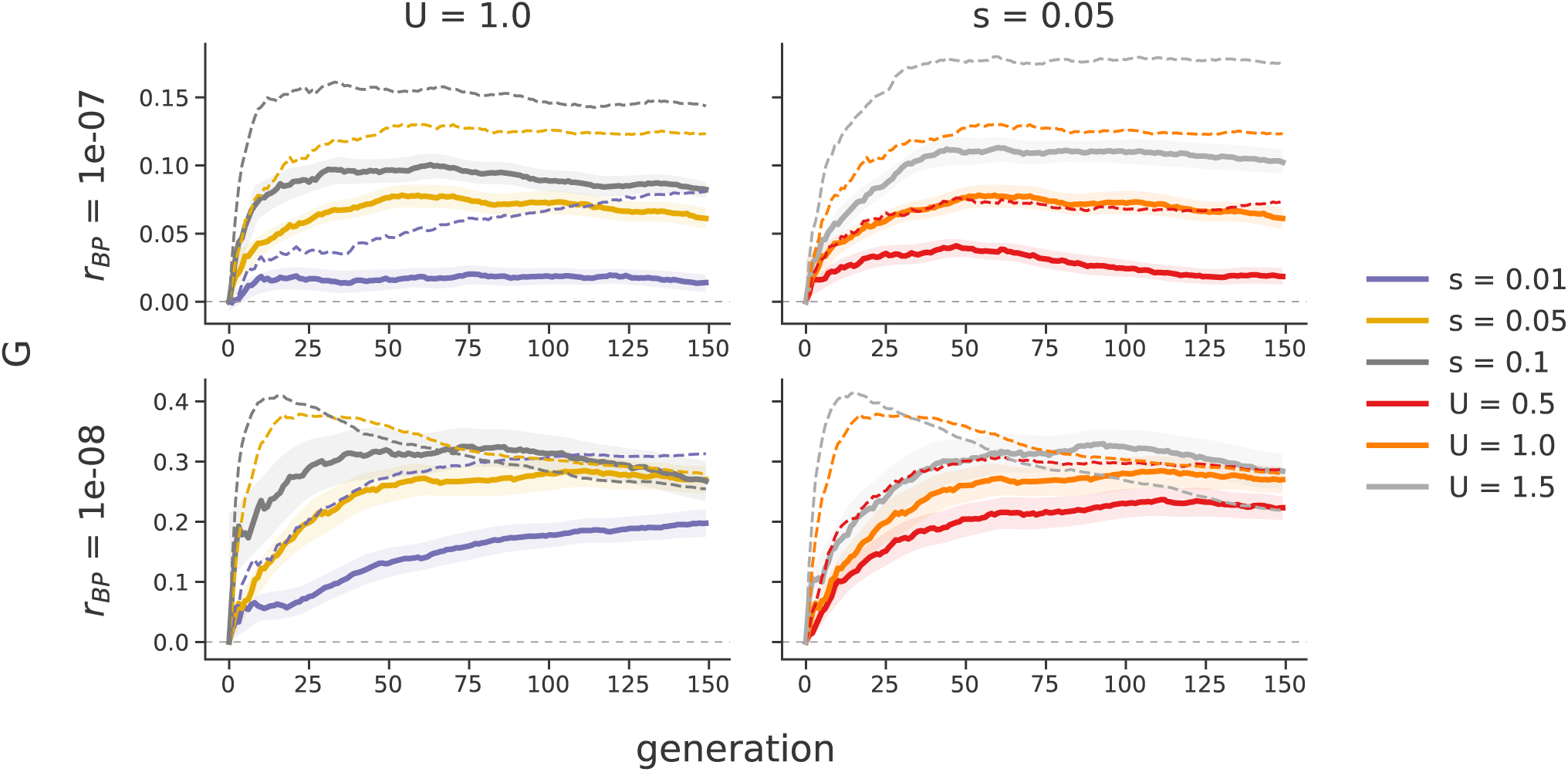
The trajectories of *G*(*t*) through time under background selection, under different recombination rates (*r*_*BP*_, rows), selection coefficients (*s*), and deleterious mutation rates (*U*). The first column fixes *U* = 1.0, and *s* varies, while in the second column *s* = 0.05 is held constant, and *U* varies. *G*(*t*) is calculated using both including fixed sites (solid lines) and not including fixed sites (dashed lines).

#### S8.6 Truncation Selection

We also explored how directional truncation selection generates temporal covariances. In these simulations we select the top 10%, 25%, or 50% of the phenotypic distribution of individuals to form the next generation. We burnin these simulations using a neutral burnin routine, where trait alleles are selectively neutral until directional selection is imposed. More extreme directional truncation selection generated larger initial covariances (Supplementary Material Figure S18B). However, weaker truncation selection generated more sustained positive covariances and so have a larger long-term impact on the variance in allele frequency change. We again noticed fixation or loss of sites has a strong effect on temporal covariance and *G*(*t*) under truncation selection. Here, since only a (potentially small) fraction of individuals contribute to the next generation, sites can fix over very short timescales. Furthermore, the small number of effective breeders contributing to the next generation shrinks *N*_*e*_ considerably, which increases Var(*p*_*t*_ − *p*_0_), the denominator of *G*(*t*). We see both the effect of handling fixed/lost sites differently and the faster rate of drift in Supplementary Materials Figure S18A, where weaker truncation selection actually has higher levels of *G*(*t*). Looking just at temporal covariances, we find that stronger truncation selection (e.g. a smaller tail of individuals selected) does lead to greater temporal covariances. Overall, these truncation selection simulations demonstrate the value of considering both absolute measures of selection’s effect on allele frequency changes, i.e. temporal covariance, as well as measures relative to drift, i.e. the *G*(*t*) trajectories. While selecting a smaller tail of individuals is associated with *stronger* selection, it also is leading to a faster rate of drift, which can distort conclusions inferred from considering the *G*(*t*) trajectories alone.

**Figure S16:**
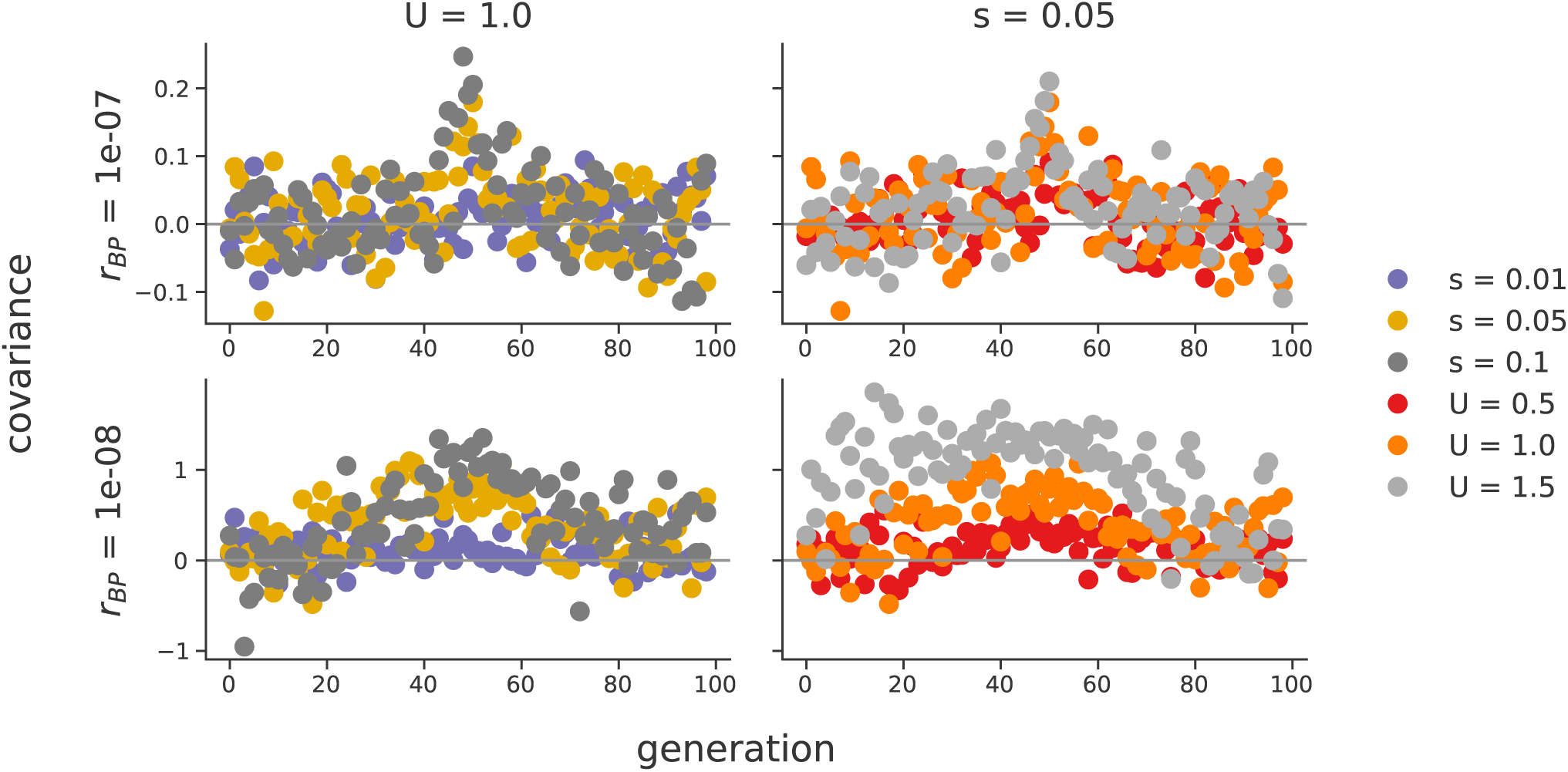
The temporal covariances, Cov(Δ*p*_50_, Δ*p*_*t*_) (where *t* varies along the x-axis) created by background selection, under different recombination rates (*r*_*BP*_, rows), selection coefficients (*s*), and deleterious mutation rates (*U*). Unlike directional selection figures, where we choose the reference generation to be the first generation after the onset of selection, here we choose an arbitrary reference generation (generation 50). The symmetry of temporal covariance around the reference generation, is expected, since unlike directional selection the level of additive genetic variance for fitness has hit mutation-selection-drift balance. Note that the first column sets constant *U* = 1.0, and *s* varies, while the second column sets *s* = 0.05 constant, and varies *U*.

#### S8.7 The Effect of Fixations in the Empirical Datasets

In our simulation results, we noticed the temporal covariances and *G*(*t*) statistics can differ depending on how allele frequencies of zero or one are handled. Generally, temporal covariances should be calculated on polymorphic sites; once a site has reached fixation or loss, its allele frequency change Δ*p*_*t*_ = 0 and including these sites in the temporal covariance calculation can lead to biases. However, with *sample* allele frequencies, rather than population frequencies, a site with observed frequency zero or one may still be segregating, but by chance not sampled at a timepoint. Here, we discuss the effect of including sites with frequency zero or one, and show our empirical results are not qualitatively different when analyzed excluding fixed sites.

With empirical data calculated on sample allele frequencies, low frequency minor alleles may not be sampled at some timepoints, and excluding these observations (instead of treating it as a trajectory that has a 0 frequency timepoint) biases estimates of quantities such as *N*_*e*_ towards intermediate frequency alleles. Additionally, we tried only dropping fixed or loss sites from the temporal covariance calculations that were at the end or the beginning of a trajectory (e.g., as if the site was created by a new mutation or fixed); while this ameliorated some of this bias it did not remove all of it. Overall, we found by trying all these approaches that not removing fixed or lost sites was the best way to deal with sample allele frequencies that could be missing from some timepoints.

To ensure that our findings were robust to handling sites with a frequency of zero or one differently, we regenerated Figures 1 and Figure 3 but excluded frequencies of zero or one. Specifically, we wanted to ensure that our finding that temporal covariances were negative at later timepoints was not spuriously caused by the way fixed or lost sites are handled. We see no qualitative difference (Supplementary Material Figures S19 and S20) that emerges when sites with frequency zero or one are excluded.

**Figure S17:**
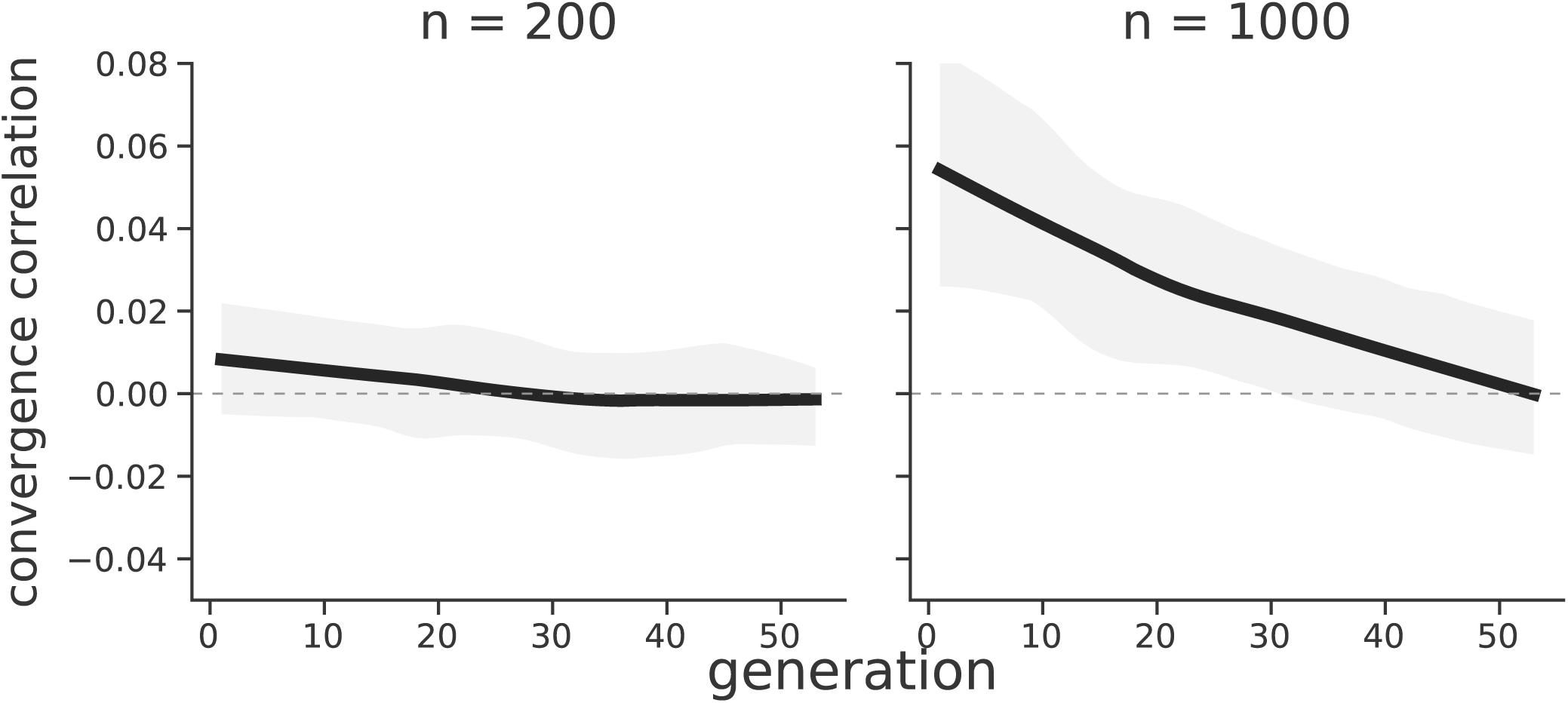
The convergence correlation created by background selection through time, since the population split. The replicate population size varies between the two panels. Values are averaged over 30 replicate simulations, while the interval is a 95% confidence interval.

**Figure S18:**
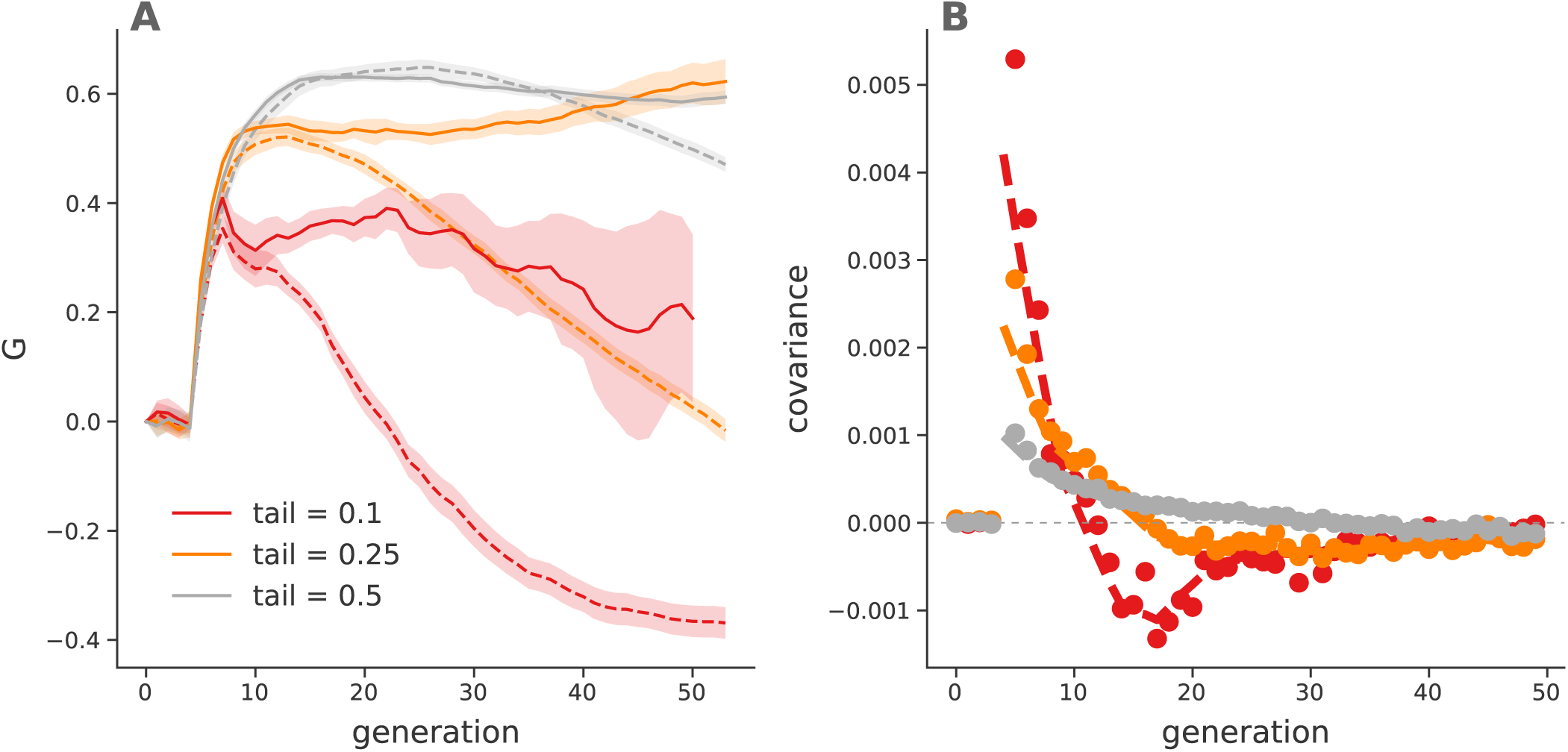
*G*(*t*) trajectories (A) and temporal covariances (B) from truncation selection simulations for different numbers of individuals selected (line color). Dashed lines indicate *G*(*t*) trajectories and temporal covariances calculated *including* fixed sites, while the solid lines exclude fixed sites. All values are averaged over 30 replicate simulations; the lines in the right figure are loess smoothed, while points are averages. The solid lines of the temporal covariances have been excluded in the left figure for clarity, but are similar except they do not become negative.

**Figure S19:**
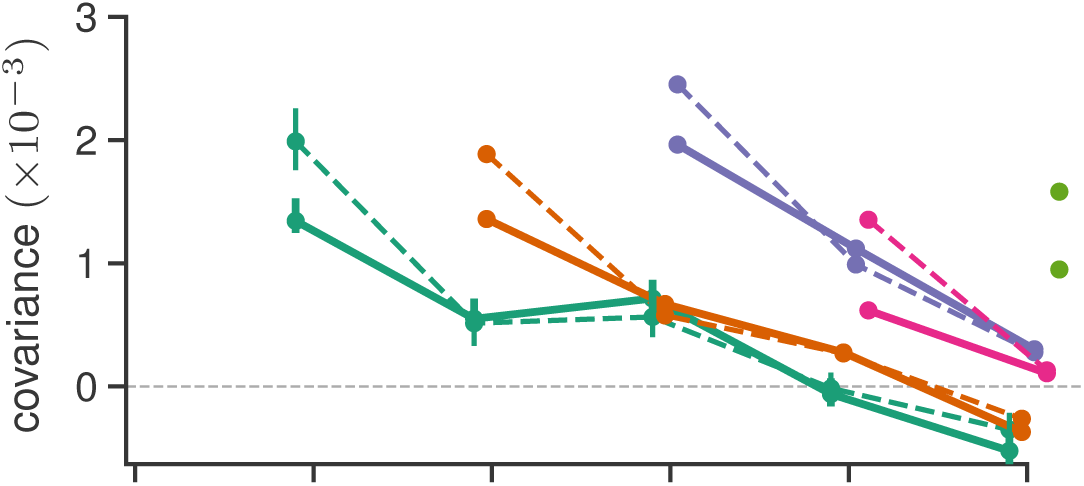
The effect of excluding fixed/lost sites in the calculation of the temporal covariances and *G*(*t*) trajectories of the Barghi et al. (2019) data. Dashed lines are those including fixed/lost sites (i.e. the original Figure 1), and solid lines are excluding fixed/lost sites.

**Figure S20:**
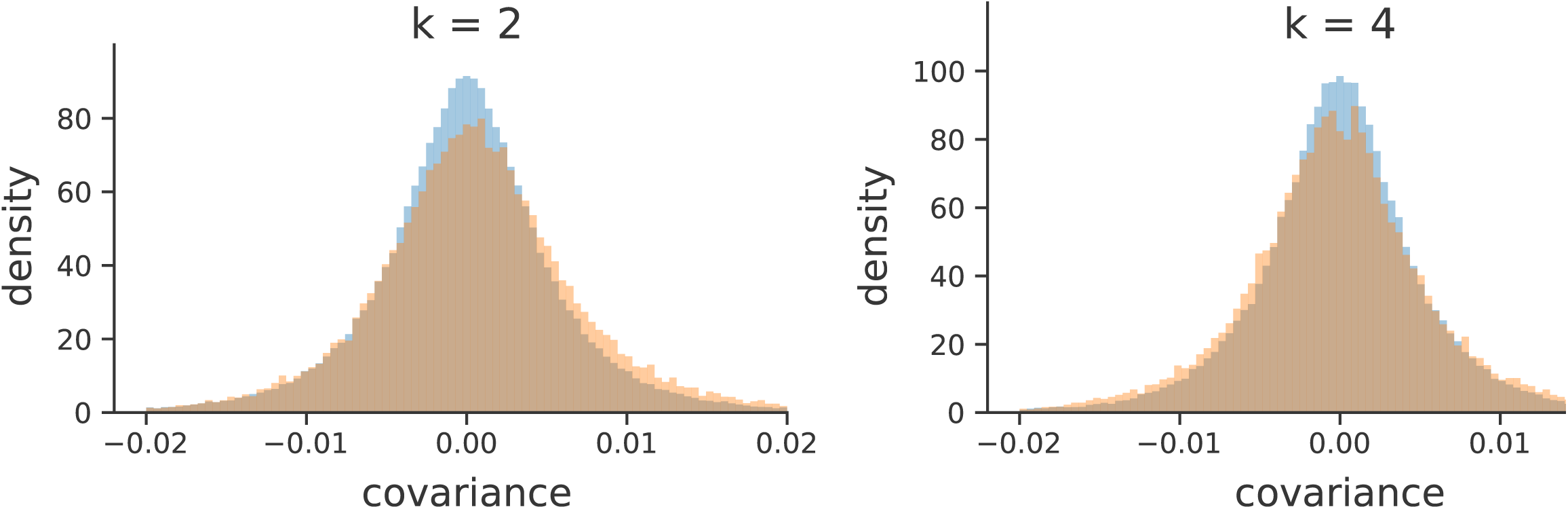
A version of Figure 3 (A) and (B) excluding fixed and lost sites. The same qualitative pattern holds as the original figure, which did not exclude fixed and lost sites: there is an enrichment of positive temporal covariances between near timepoints (*k* = 2) in the Barghi et al. (2019) study, and an excess of negative temporal covariances at more distant timepoints (*k* = 2).

### S9 Additional Figures

#### S9.1 PCA of Barghi et al. (2019) replicates

**Figure S21:**
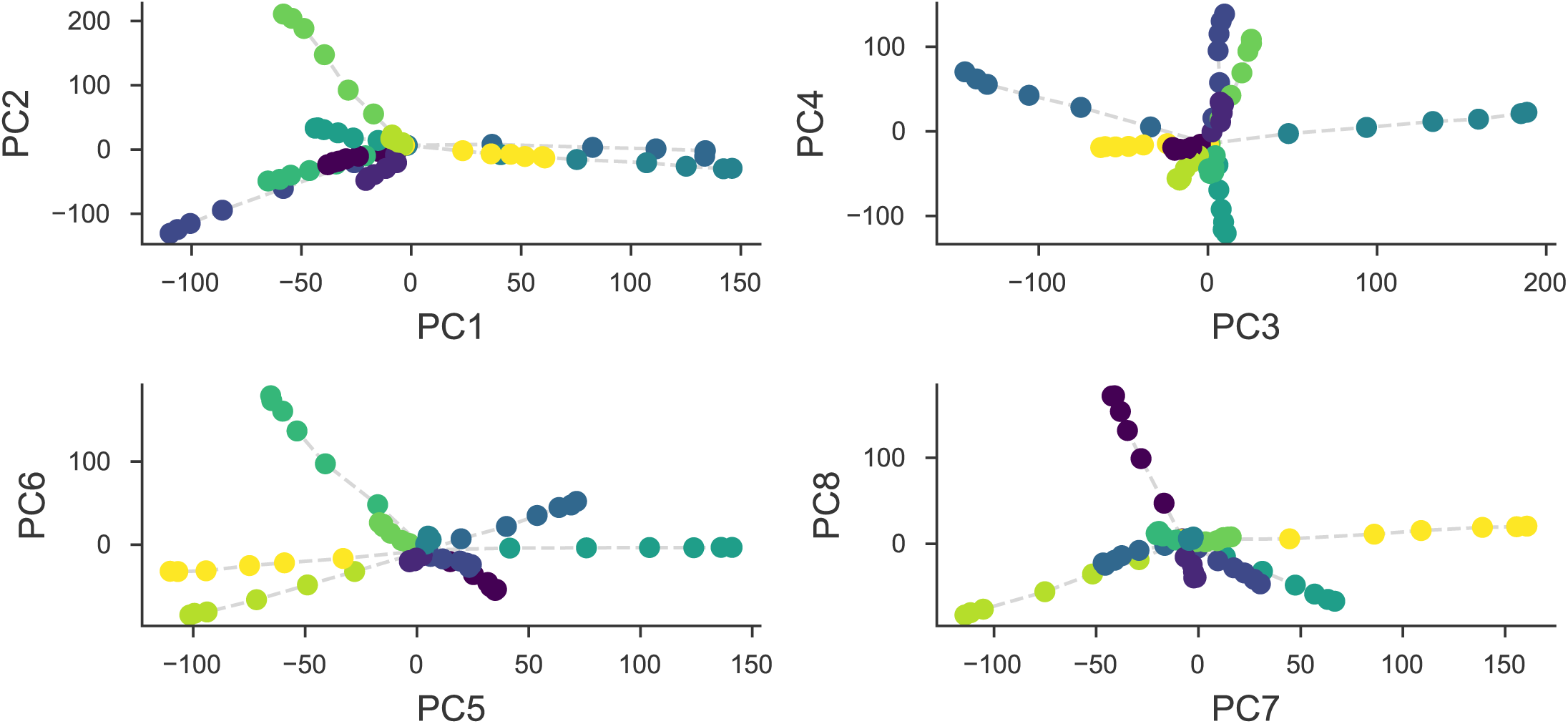
A PCA on the centered and standardized population frequencies for each replicate (each color) for all its sequenced timepoints (the connected series of points). All replicates start from the same source population, and thus are overlapping in the center; as each replicate evolves independently it diverges from the other replicates in PCA space.

#### S9.2 Bias Correction for Barghi et al. (2019)

We have investigated the effectiveness of our correction on real data by exploiting the relationship between sampling depth and the magnitude of the variance and covariance biases, and comparing the observed variances and covariances before and after correction. We plot the variance and covariance (between adjacent time intervals) before and after the bias correction against the average sample depth in 100kb genomic windows in Figure S22. Overall, we find the biased-correction procedure removes the relationship between variance and covariance and depth, indicating it is working adequately.

**Figure S22:**
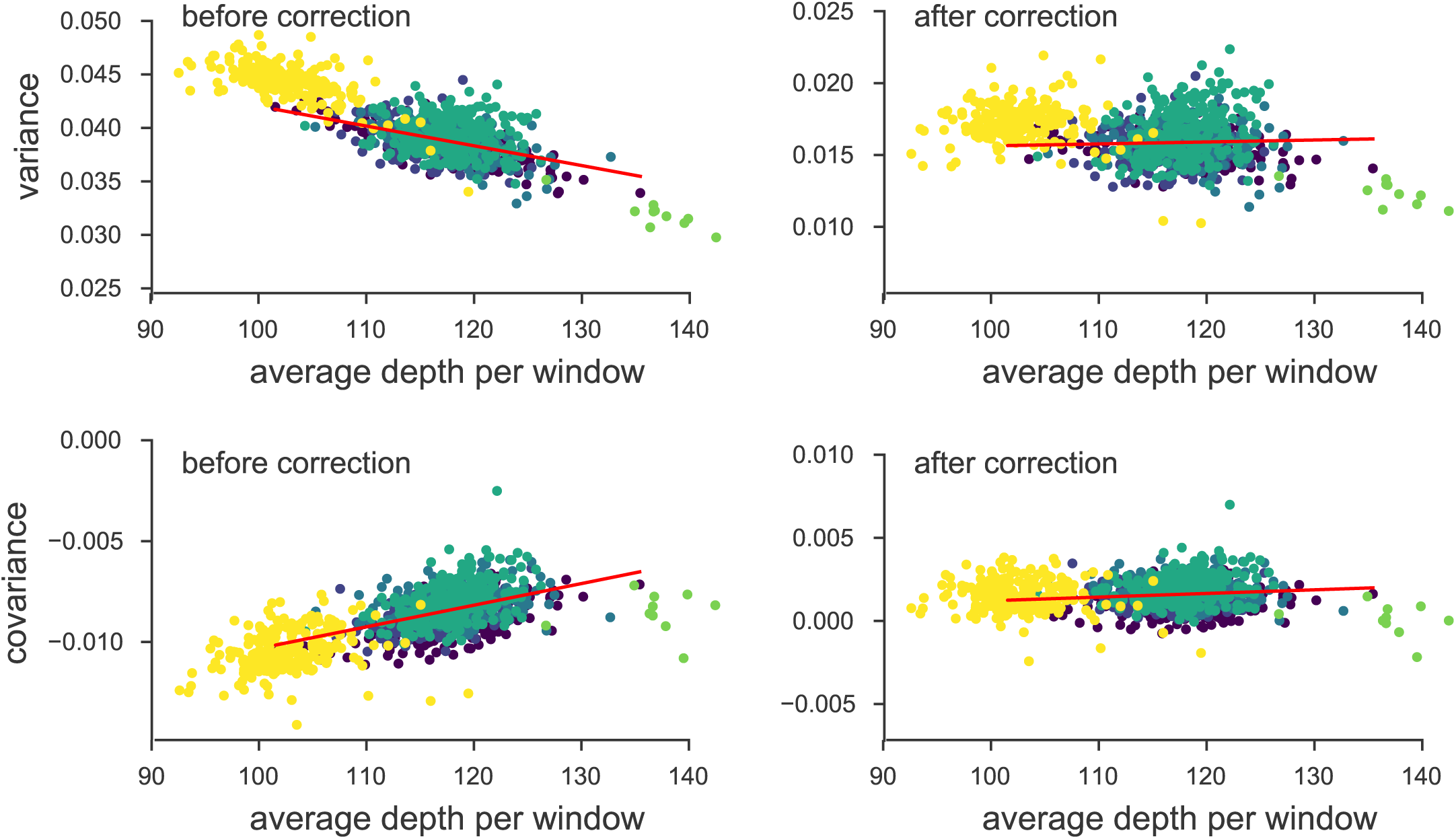
The variance and covariances from the Barghi et al. (2019) study, calculated in 100kb genomic windows plotted against average depth in a window before and after bias correction. Each panel has a leastsquares estimate between the variance and covariance, and the average depth. Overall, the bias correction corrects sampling bias in both the variance and covariance such that the relationship with depth is constant. Colors indicate the different chromosomes of *D. simulans*; we have excluded the X chromosome (yellow points) and chromosome 4 points (green points to far right) from the regression due to large differences in average coverage.

#### S9.3 Barghi et al. (2019) Temporal Covariances Per Replicate

**Figure S23:**
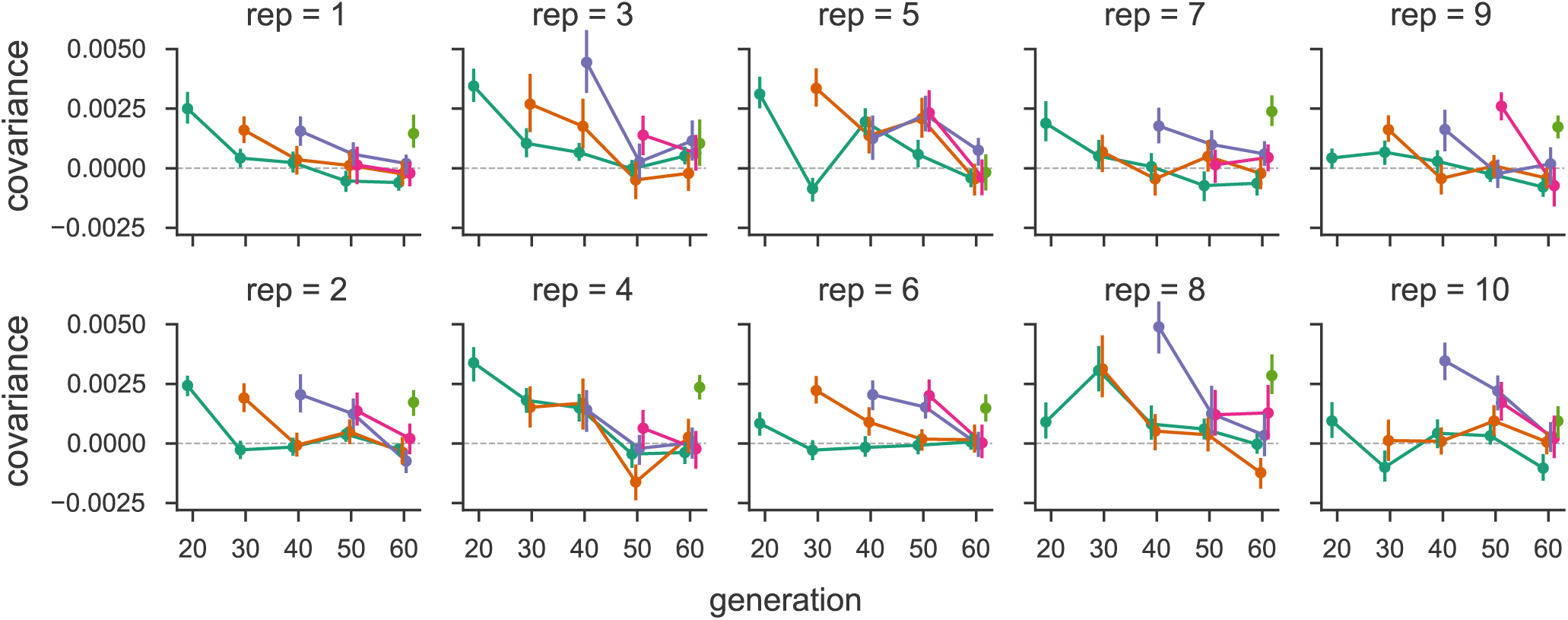
The temporal covariances from the Barghi et al. (2019) study, for each replicate individually. As in Figure 1, each line follows the temporal covariances from some initial reference generation through time, which represent the rows of temporal covariance matrix.

**Table S1:**
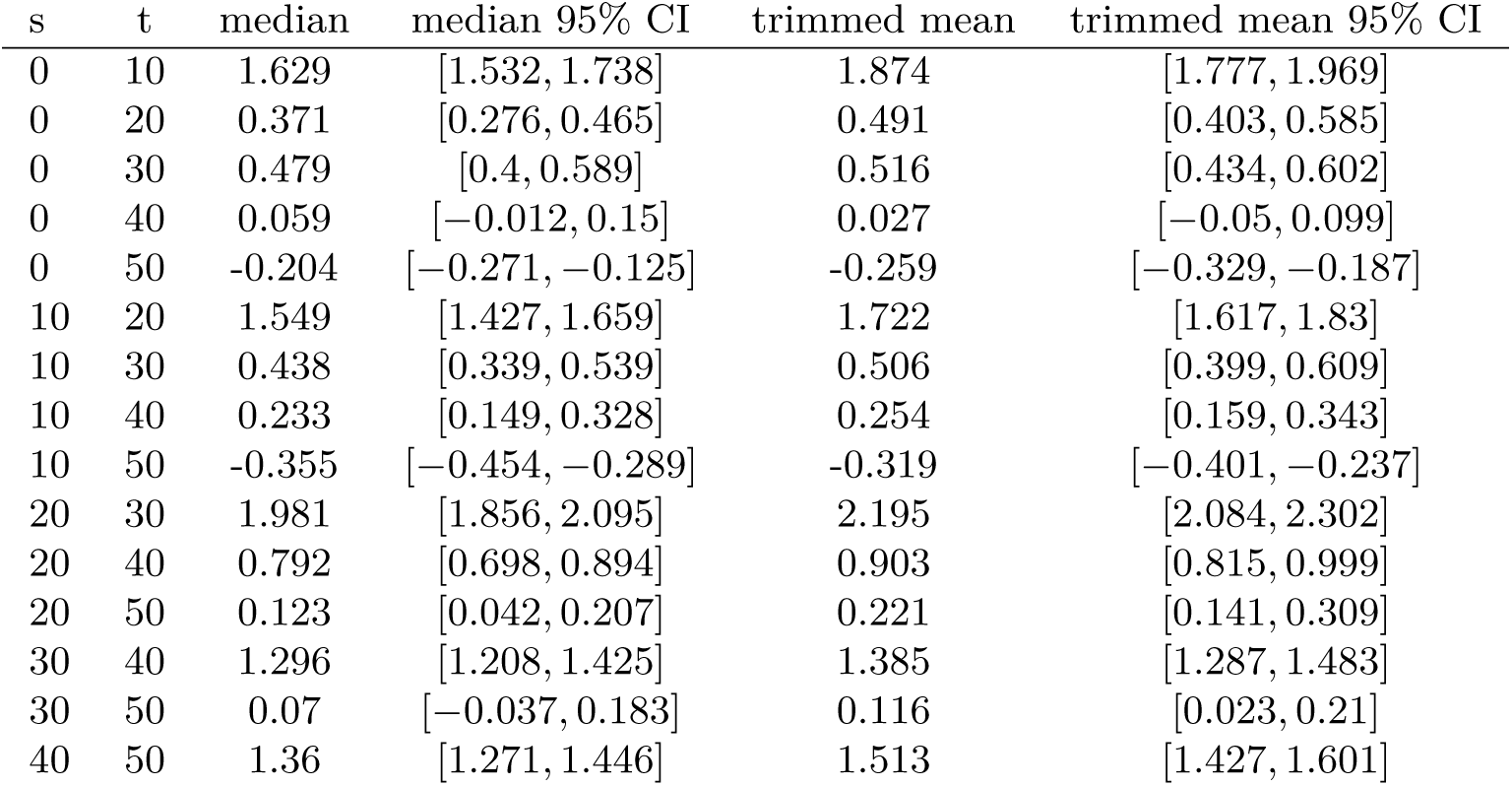
Table of median of windowed covariance estimates (Cov(Δ*p*_*s*_, Δ*p*_*t*_) × 100) between generations *t* and *s* and the trimmed mean windowed covariance which excludes the lower and upper 5% windows with the highest covariance.

#### S9.4 Barghi et al. (2019) Trimmed Window Covariances

Here we report median and trimmed mean of the windowed covariances (Supplementary Table S1). We note that the median covariance is also limiting result of a trimmed mean that symmetrically excludes the upper and lower *α* tails to calculate the trimmed average windowed covariance. As *α* increases to 0.5, the trimmed covariance converges to the median windowed covariance (by the definition of the median; see Supplementary Figure S24). Thus our genomic temporal covariances are non-zero due to the impact of selection on many genomic windows.

**Figure S24:**
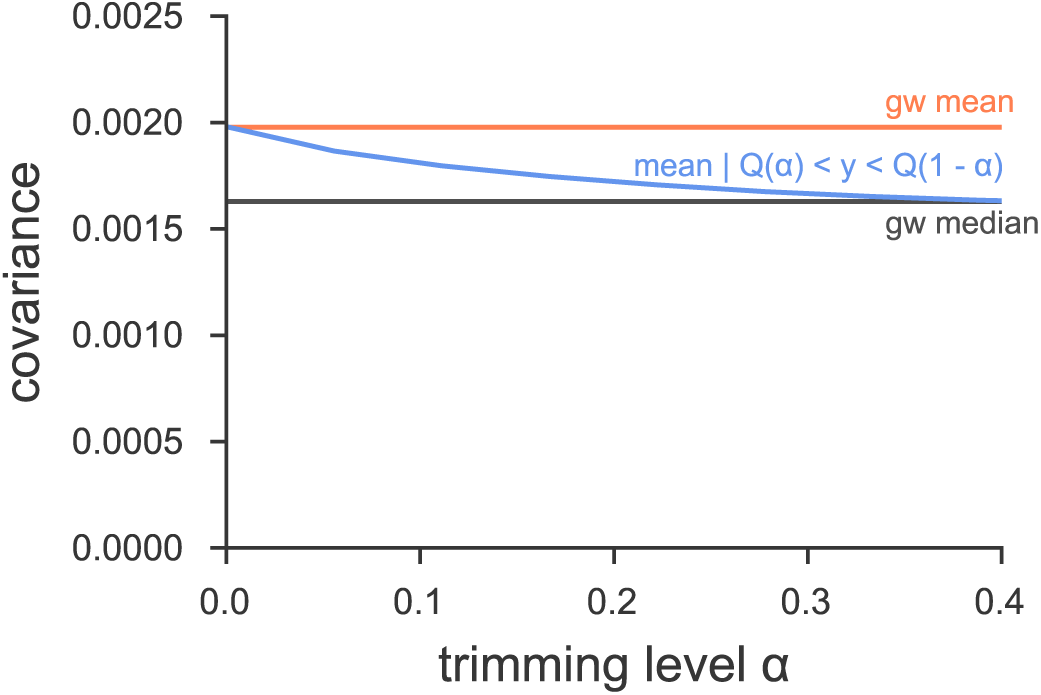
The genome-wide covariance (Cov(Δ*p*_0_, Δ*p*_10_) pooling all replicates) averaged (red line) and the median windowed covariance (blue) for the Barghi et al. (2019) dataset. The trimmed average window covariance, excluding the *α* lower and upper tails, converges to the median windowed covariance. This indicates that genome-wide covariances are not being overly dominated by a large-effect loci in few windows.

#### S9.5 Barghi et al. (2019) Empirical Null and Windowed Covariance Distributions

**Figure S25:**
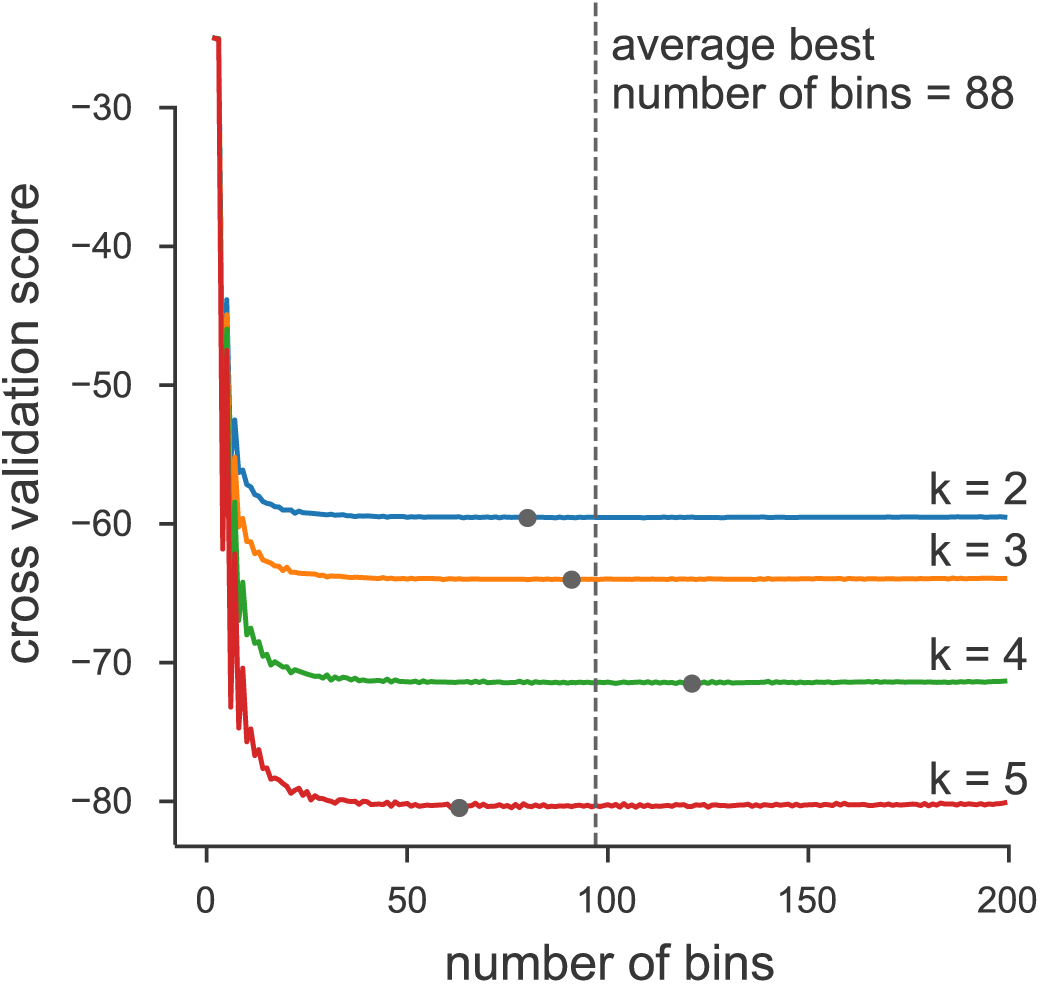
We chose number of bins used in the histograms of Figure 3 via an analytic expression for the cross-validation risk, based on the equation 6.16 of (Wasserman 2006, p. 129). Above, we plot the cross-validation risk for various numbers of bins, for each of the four off-diagonals of the temporal covariance matrix that we analyze. Overall, because the number of data points is large, oversmoothing is less of a problem, leading the cross-validation risk to be relatively flat across a large number of bins. Each gray point indicates the minimal risk for a particular off-diagonal, and the dashed line indicates the best average binwidth across off-diagonals.

**Figure S26:**
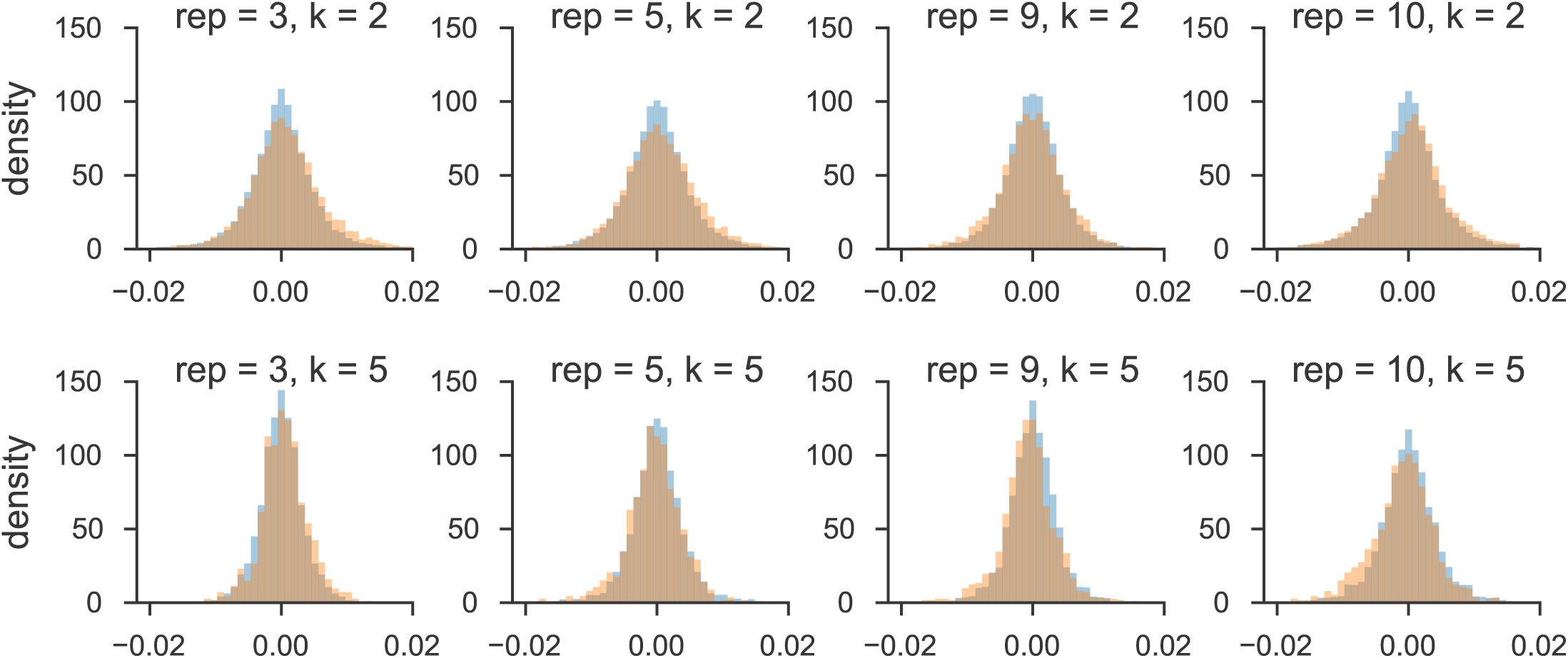
The distribution of windowed temporal covariances alongside the empirical neutral null for five randomly sampled replicates (columns), for *k* = 2 (first row) and *k* = 5 (second row). The main figure of the paper pools all replicate window and empirical neutral null covariances; we show here the windowed temporal covariances tend to shift from being positive (a heavier right tail) to become more negative (a heavier left tail) through time within particular replicates.

**Figure S27:**
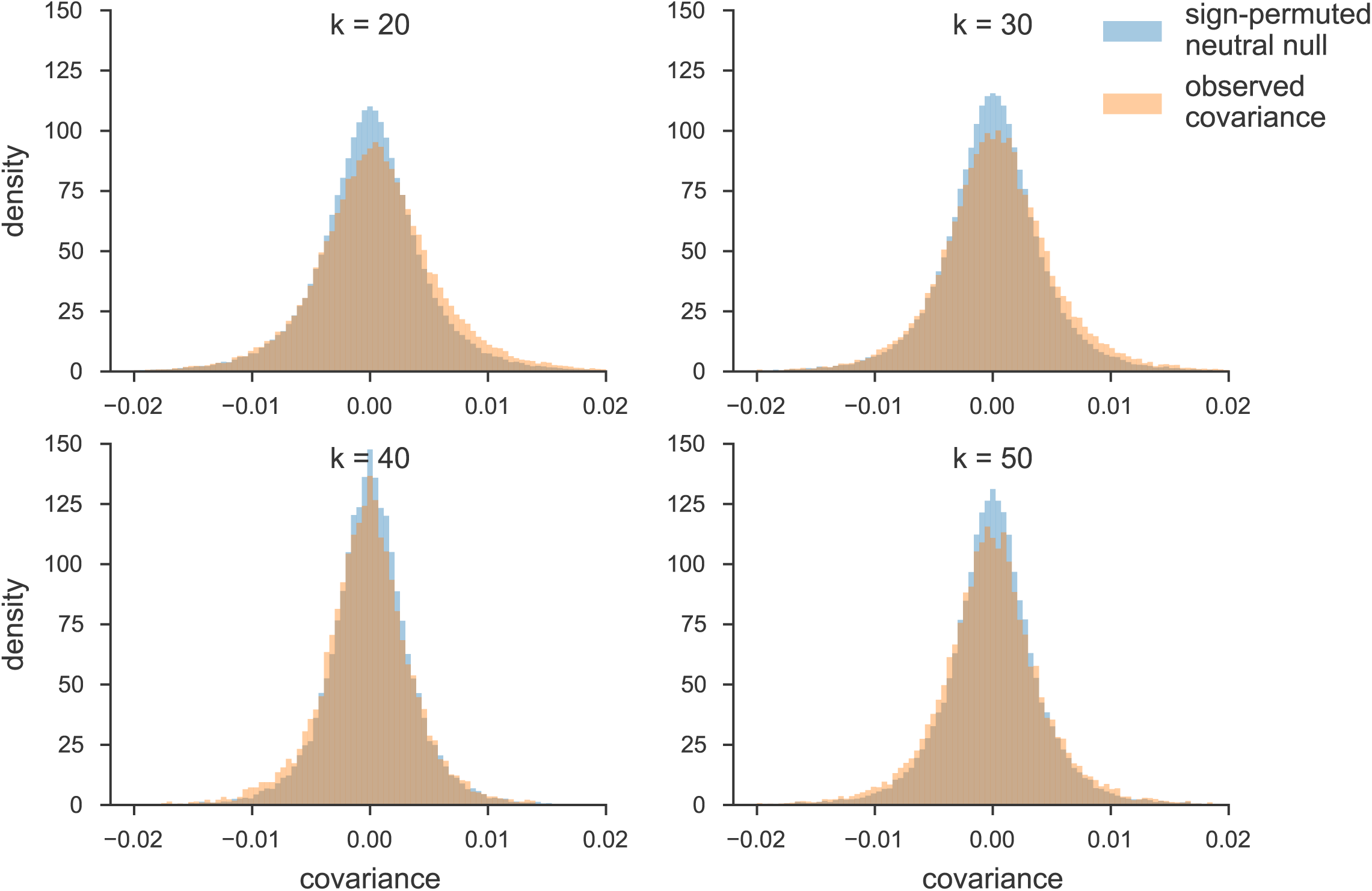
The distribution of temporal covariances calculated across 100kb genomic windows from Barghi et al. (2019)’s study (orange) and the block sign permuted empirical neutral null distribution of the windowed covariances (blue). Each panel shows these windowed covariances and the empirical null distribution for covariances Cov(Δ*p*_*t*_, Δ*p*_*t*+*k*_), *k* is the number of generations between allele frequency changes.

#### S9.6 Barghi et al. (2019) Tail Probabilities for Windowed Covariances Distributions

**Figure S28:**
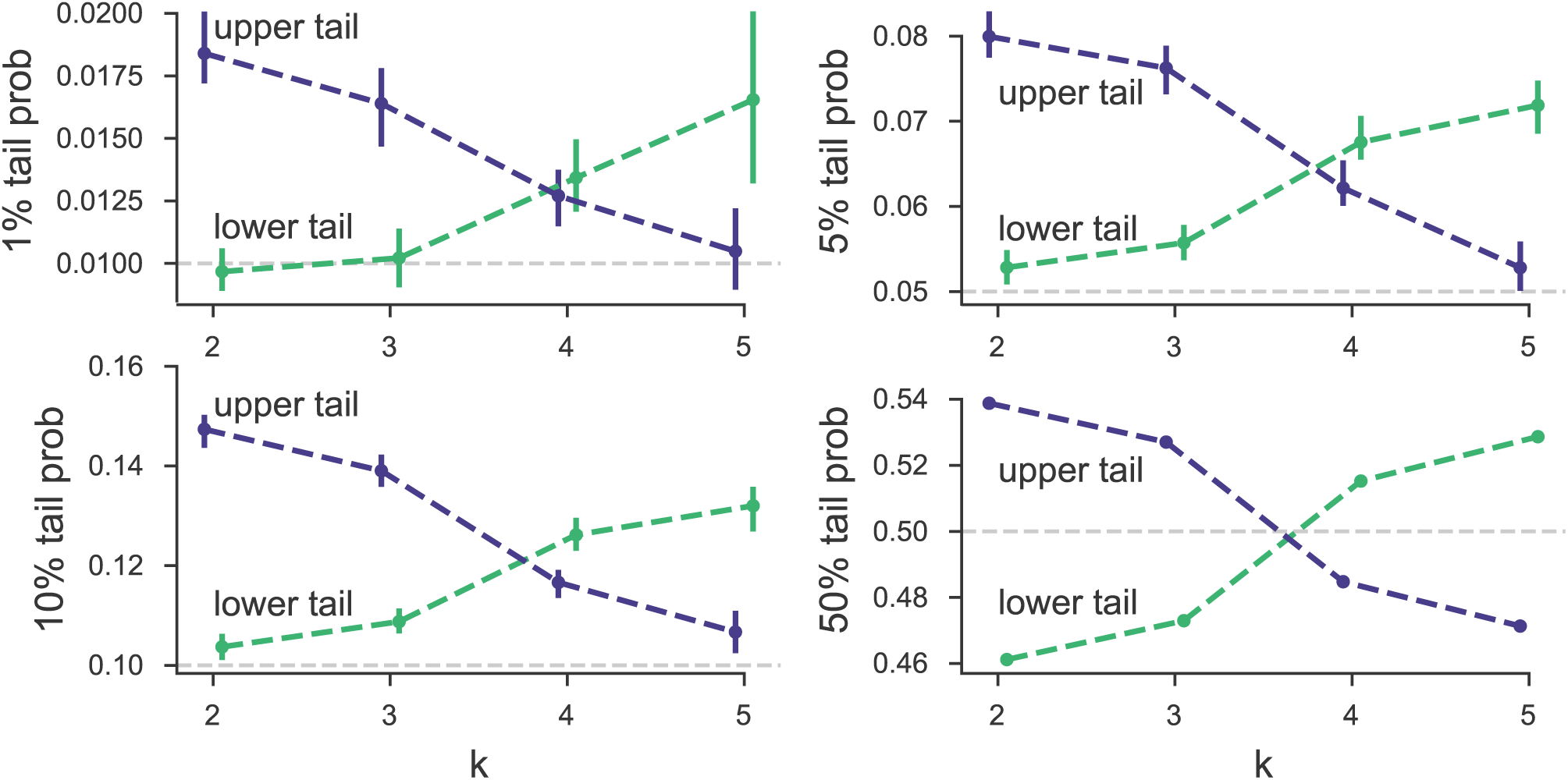
Barghi et al. (2019) tail probabilities compared to sign-permuted empirical null distribution for various *α* levels.

**Figure S29:**
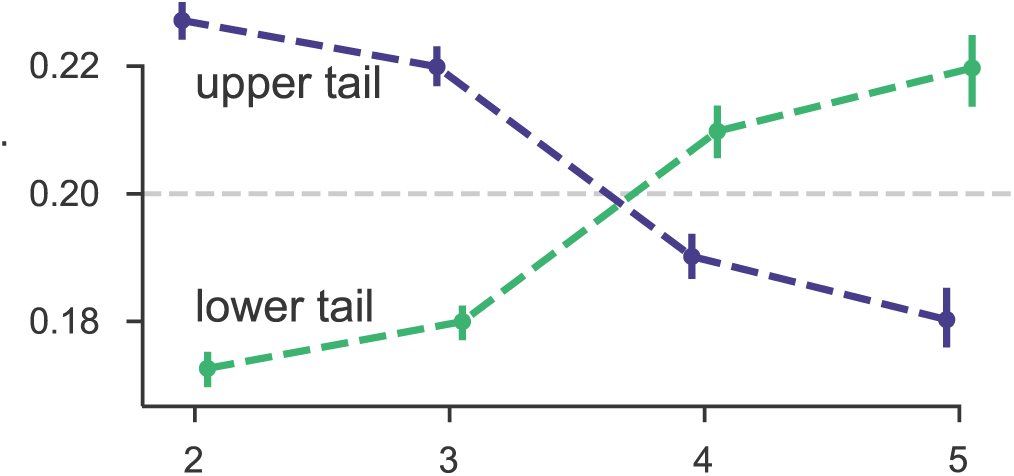
The 20% lower and upper tail probabilities for the observed windowed covariances from the Barghi et al. (2019) study, based on sign-permuting at the chromosome level. This permutation empirical null is robust to long-range linkage disequilibrium acting over entire chromosomes (see Supplementary Material section S5).

