## Supplementary material for "Estimating the genome-wide contribution of selection to temporal allele frequency change": revision difference

**Table 1:** Summary of the main selection studies we analyzed

| Study | Species | Selection | Replicates | Pop. Size† | Gens. | Tim |
|---|---|---|---|---|---|---|
| ~~Kelly and Hughes (2019)~~ Kelly and Hughes (2019) | *D. simulans* | lab adaptation | 3 | ∼1100 | 14 | |
| ~~Barghi et al. (2019)~~ Barghi et al. (2019) | *D. simulans* | lab adaptation | 10 | ∼1000 | 60 | |
| ~~Castro et al. (2019)~~ Castro et al. (2019) | *M. musculus* | tibiae length | 2 | 32 | 17 | |
| | | control | 1 | 28 | | |

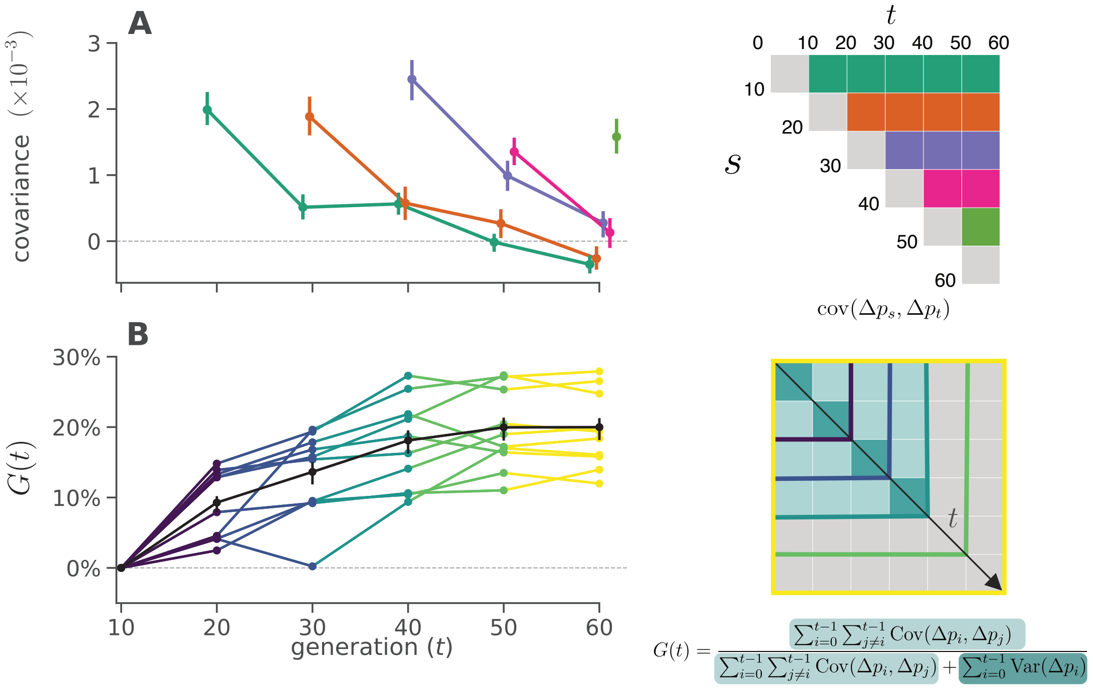

**Figure 1:** **A**: Temporal covariance, averaged across all ten replicate populations, through time from the ~~Barghi et al. (2019)~~ Barghi et al. (2019) study. Each line depicts the temporal covariance $\mathrm{Cov}(\Delta p_s, \Delta p_t)$ from some reference generation $s$ to a later time $t$ which varies along the x-axis; each line corresponds to a row of the upper-triangle of the temporal covariance matrix with the same color (upper right). The ranges around each point are 95% block-bootstrap confidence intervals. **B**: ~~The~~ A lower bound on the proportion of the total variance in allele frequency change explained by linked selection, $G(t)$, as it varies through time $t$ along the x-axis. The black line is the $G(t)$ averaged across replicates, with the 95% block-bootstrap confidence interval. The other lines are the $G(t)$ for each individual replicate, with colors indicating what subset of the temporal-covariance matrix to the right is being included in the calculation of $G(t)$.

Appendix, Fig. S23.

Since our covariances are averages over loci, the covariance estimate could be strongly affected by a few outlier regions. To test whether large outlier regions drive the genome-wide signal we see in the ~~Barghi et al. (2019)~~ Barghi et al. (2019) data, we calculate the covariances in 100kb windows along the genome (we refer to these as windowed covariances throughout) and take the median windowed covariance, and trimmed-mean windowed covariance, as a measure of the genome-wide covariance robust to large-effect loci. These robust estimates (~~Supplementary~~ SI Appendix, Table S1 and ~~Supplementary Figure~~ SI Appendix, Fig. S24) confirm the patterns we see using the mean covariance, ~~confirming~~ establishing that genomic temporal covariances are non-zero due to the impact of selection acting across many genomic regions.

While the presence of positive temporal covariances is consistent with selection affecting allele frequencies over time, this measure is not easily interpretable. We can calculate a more intuitive measure from the temporal covariances to quantify the impact of selection on allele frequency change: the ratio of total covariance in allele frequency change to the total variance in allele frequency change. We denote the change in allele frequency as $\Delta p_t = p_{t+1} - p_t$, where $p_t$ is the allele frequency in generation $t$. Since the total variation in allele frequency change can be partitioned into variance and covariance components, $\mathrm{Var}(p_t - p_0) = \sum_{i=0}^{t-1} \mathrm{Var}(\Delta p_i) + \sum_{i=0}^{t-1} \sum_{j \neq i}^{t-1} \mathrm{Cov}(\Delta p_i, \Delta p_j)$ (we bias-correct these for correct for biases due to sequencing depth), and the covariances are zero when drift acts alone, this is a lower bound on how much of the variance in allele frequency change is caused by linked selection (Buffalo and Coop 2019). We call this measure $G(t)$, defined as

$$\operatorname{cor}(\Delta p_s, \Delta p_t) = \frac{\mathbb{E}_{A \neq B}\left(\operatorname{Cov}(\Delta p_{s,A}, \Delta p_{t,B})\right)}{\mathbb{E}_{A \neq B}\left(\sqrt{\operatorname{Var}(\Delta p_{s,A})\operatorname{Var}(\Delta p_{t,B})}\right)} \tag{2}$$

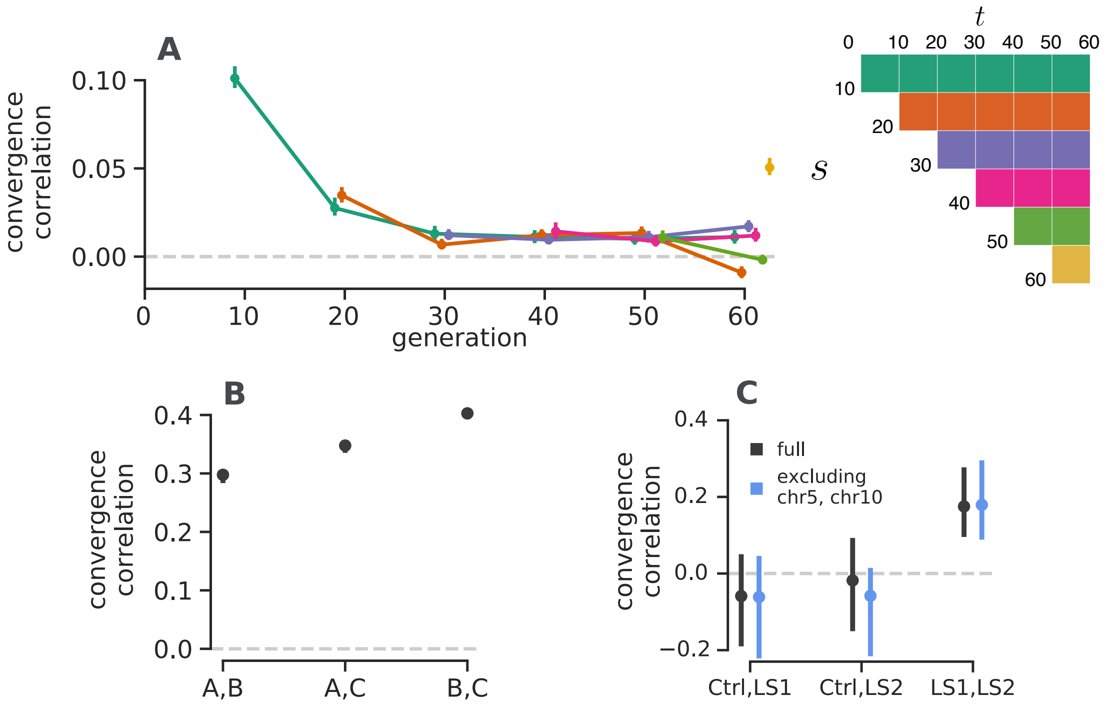

**Figure 2:** **A**: The convergence correlations, averaged across Barghi et al. (2019) Barghi et al. (2019) replicate pairs, through time. Each line represents the convergence correlation $\text{cor}(\Delta p_s, \Delta p_s)$ $\text{cor}(\Delta p_s, \Delta p_t)$ from a starting reference generation $s$ to a later time $t$, which varies along the x-axis; each line corresponds to a row of the temporal convergence correlation matrix depicted to the right (where the diagonal elements represent the convergence correlations between the same timepoints across replicate populations). We note that convergent correlation for the last timepoint is an outlier; we are unsure as to the cause of this, e.g. it does not appear to be driven by a single pair of replicates. **B**: The convergence correlations between individual pairs of replicates in the Kelly and Hughes (2019) Kelly and Hughes (2019) data (note the confidence intervals are plotted, but are small on this y-axis scale). **C**: The convergence correlations between individual pairs of replicates in Castro et al. (2019) data, for the two selection lines (LS1 and LS2) and the control (Ctrl); gray CIs are those using the complete dataset, blue CIs exclude chromosomes 5 and 10 which harbor the two regions Castro et al. (2019) Castro et al. (2019) found to have signals of parallel selection between LS1 and LS2. Through simulations, we have found that the differences in convergence correlation confidence interval widths between these *Drosophila* studies and the Longshanks study are due to the differing population sizes.

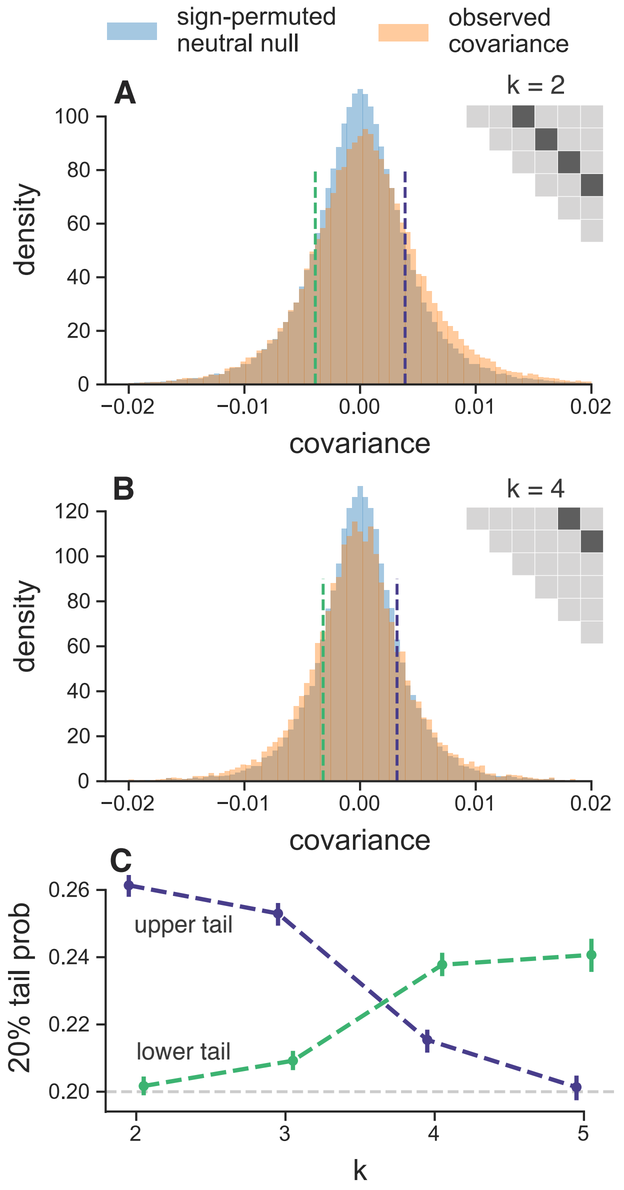

**Figure 3:** **A**, **B**: The distribution of temporal covariances calculated in 100kb genomic windows from the ~~Barghi et al. (2019)~~ Barghi et al. (2019) study, plotted alongside an empirical neutral null distribution created by recalculating the windowed covariances on 1,000 sign permutations of allele frequency changes within tiles. The ~~histogram bin~~ number of histogram bins is 88, chosen by cross validation (~~Supplementary Materials~~ SI Appendix, section S25). In subfigure **A**, windowed covariances $\mathrm{Cov}(\Delta p_t, \Delta p_{t+k})$ are separated by $k = 2 \times 10$ generations and in subfigure **A** the covariances are separated by $k = 4 \times 10$ generations; each $k$ is an off-diagonal from the variance diagonal of the temporal covariance matrix (see cartoon of upper-triangle of covariance matrix in subfigures **A** and **B**, where the first diagonal is the variance, and the dark gray indicates which off-diagonal of the covariance matrix is plotted in the histograms). **C**: The lower and upper tail probabilities of the observed windowed covariances, at 20% and 80% quintiles of the empirical neutral null distribution, for varying time between allele frequency changes (i.e. which off-diagonal $k$). The confidence intervals are 95% block-bootstrap confidence intervals, and the light gray dashed line indicates the 20% tail probability expected under the neutral null. Similar figures for different values of $k$ are in ~~Supplementary Figures~~ SI Appendix, section S27.

~~These simulation results are often~~ Averaging across replicates, these simulation results show $G(t)$ is relatively insensitive to the number of loci underlying the trait~~, suggesting that they are capturing highly polygenic signals. Indeed~~. However, if only a small number of loci influence the trait, the $G(t)$ trajectories are typically much more stochastic ~~across replicates(compare Figure 1B to Supplementary Material Figure S6). This again suggests~~ *across* replicates. This reaffirms that the genome-wide linked selection response we see in the ~~Barghi et al. (2019)~~ Barghi et al. (2019) data is highly polygenic ~~. Using~~ (compare Figure 1B to SI Appendix, Fig. S6). Furthermore, using our simulations we find that sampling only every 10 generations does indeed mean that our estimates of $G(t)$ are an underestimate of the ~~contribution~~ proportional effect of linked selection as they cannot include the covariance between closely spaced generations (see ~~Supplementary Material Figure~~ SI Appendix, Fig. S14).

Additionally, we explored other ~~selection schemes~~modes of selection with simulations. We find that the long term dynamics of the covariances under directional truncation selection, which generates substantial epistasis, are richer than we see under ~~GSS~~ Gaussian Stabilizing Selection (GSS) and multiplicative selection (~~Supplementary Material Figure~~ SI Appendix, Fig. S18). We also

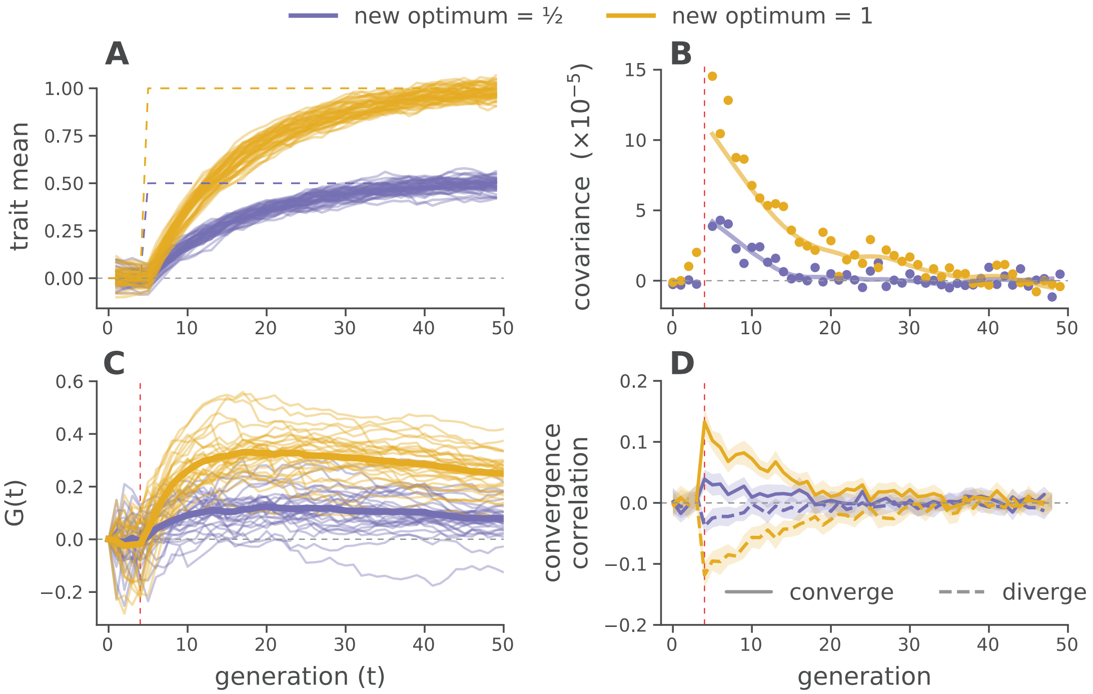

**Figure 4:** Forward-in-time simulations ~~of~~ demonstrate how temporal covariance, $G(t)$ trajectories, and convergence correlations arise during optima shifts of two different magnitudes, under Gaussian stabilizing selection. (**A**) Trait means across 30 replicate before and after optima shifts (solid lines), for two different magnitudes (indicated by color). The optimal trait values are indicated by the purple and yellow dashed lines. (**B**) Mean temporal covariance $\text{Cov}(\Delta p_5, \Delta p_t)$ across 30 simulation replicates, where $t$ varies along the x-axis (points), with a loess-smoothed average (solid line). (**C**) $G(t)$ trajectories through time, for 30 replicate simulations across two optima shifts. The solid line is a loess-smoothed average. (**D**) The convergence correlations between two populations (each 1000 diploids) split from a common population, that underwent either an optima shift in the same direction (converge) and opposite directions (diverge) at generation five. In subfigures (**B**), (**C**), and (**D**), directional selection begins at generation five, when the optima shifts; this is indicated by the vertical dashed red line (see SI Appendix Section S8.2 for details on these simulations).

It is worth building some intuition why temporal covariance allows us to detect such faint signals of polygenic linked selection from temporal genomic data. ~~Each variant is subject to both variance~~ Variance in allele frequency ~~due to~~ change is subject to both drift and sampling noise, which at any single locus may swamp the temporal covariance signal ~~and creates spurious covariances~~ due to selection, or create spurious covariances when selection is not acting. However, these spurious covariances do not share a directional signal whereas the covariances created by linked selection do; consequently, averaging across the entire genome, the temporal signal exceeds sampling noise.

### 2 ~~Materials and Methods~~

### Materials and Methods

#### 1.1 ~~Datasets Analyzed~~

#### Datasets Analyzed

We used available genomic data data from four studies: pooled population resequencing (pool-Seq) data from ~~Barghi et al. (2019), Kelly and Hughes (2019), and Bergland et al. (2014)~~Barghi et al. (2019), Kelly and Hughes (2019), and Bergland et al. (2014), and individual-level sequencing data from ~~Castro et al. (2019).~~ Castro et al. (2019). In all cases, we used the variants kept after the filtering criteria of the original studies.

#### 1.1 ~~Partitioning Unique and Shared Selection Effects in the Longshanks Study~~

#### Partitioning Unique and Shared Selection Effects in the Longshanks Study

The unselected control line in the Longshanks experiment allows us to additionally partition the total variance in allele frequency change into drift, shared effects of selection, and unshared effects of selection between selected replicates. We begin by decomposing the allele frequency change in Longshanks line 1 (LS1) as $\Delta p_{t,\text{LS1}} = \Delta_D p_{t,\text{LS1}} + \Delta_U p_{t,\text{LS1}} + \Delta_S p_{t,\text{LS}}$ where these terms are the drift in Longshanks replicate 1 ($\Delta_D p_{t,\text{LS1}}$), selection unique to the LS1 replicate ($\Delta_U p_{t,\text{LS1}}$), and selection response shared between the two Longshanks replicates ($\Delta_S p_{t,\text{LS}}$) respectively (and similarly for the Longshanks line 2, LS2). By construction, this decomposition assumes that each of these terms are uncorrelated within replicates, so the contribution of each term to the total variance in allele frequency change, $\text{Var}(\Delta p_{t,\text{LS1}})$, is the variance of that term's allele frequency change.

We estimate the effects of selection by first calculating the fraction of the total variance explained by drift. We assume the variance in allele frequency change observed in the unselected control line ($\text{Var}(\Delta p_{t,\text{Ctrl}})$) is driven entirely by neutral genetic drift, and since an identical breeding scheme was used across all three replicates (except breeders for the control line were chosen at random), we can use this as an estimate of the contribution of neutral genetic drift in the selected lines, $\text{Var}(\Delta p_{t,\text{Ctrl}}) = \text{Var}(\Delta_D p_{t,\text{LS1}}) = \text{Var}(\Delta_D p_{t,\text{LS2}})$. Then, we can estimate the increase in variance in allele frequency change due to selection as $(\text{Var}(\Delta p_{t,\text{LS1}}) + \text{Var}(\Delta p_{t,\text{LS2}}))/2 - \text{Var}(\Delta p_{t,\text{Ctrl}})$ and the shared effect of selection across selected lines as $\text{Cov}(\Delta p_{t,\text{LS1}}, \Delta p_{t,\text{LS2}})$. Finally, the covariance in allele change between replicates is used to estimate the shared effects of selection between lines, $\text{Cov}(\Delta p_{t,\text{LS1}}, \Delta p_{t,\text{LS2}}) = \text{Var}(\Delta_S p_{t,\text{LS}})$.

#### 1.1 ~~Forward-in-time Simulations~~

#### Forward-in-time Simulations

To explore how aspects of genetic architecture, ~~model~~ models of selection, and experimental design impact temporal covariance, the $G(t)$ trajectories, and convergence correlations, we ran extensive forward-in-time simulations using SLiM (Haller and Messer 2019); here we discuss the Gaussian Stabilizing Selection simulations in Figured 4, but ~~Supplementary Materials Section~~ SI Appendix, section S8 describes these simulation routines ~~in more detailas well as additional simulations we have conducted~~and others in detail.

### Supplementary ~~Material~~Information

### Contents

**1 Introduction** **2**

**2 ~~Results~~** **3**

**2 ~~Discussion~~** **12**

**2 ~~Materials and Methods~~** **14**

1.1 ~~Datasets Analyzed~~ . . . . . 14
1.1 ~~Variance and Covariance Estimates~~ . . . . . 14
1.1 ~~Estimating Uncertainty with a Block Bootstrap~~ . . . . . 15
1.1 ~~Partitioning Unique and Shared Selection Effects in the Longshanks Study~~ . . . . . 15
1.1 ~~Windowed Covariance and the Empirical Neutral Null~~ . . . . . 15
1.1 ~~Forward-in-time Simulations~~ . . . . . 16

**2 ~~Acknowledgments~~** **17**

**S1 Estimator Bias Correction** **22**

S1.1 Correcting variance bias with a single depth sampling process . . . . . 22
S1.2 Correcting variance bias with individual and depth sampling processes . . . . . 23
S1.3 Covariance Correction . . . . . 24
S1.4 Temporal-Replicate Covariance Matrix Correction . . . . . 25

**S2 Barghi et al. (2019) Temporal Covariances** **26**

**S3 Block Bootstrap Procedure** **26**

**S4 Replicate $G(t)$ and Partitioning the Variance in Allele Frequency** **27**

**S5 The Empirical Neutral Null Windowed Covariance Distribution** **29**

**S6 Bergland et al. (2014) Re-Analysis** **29**

**S7 Approximating the Reduction in Diversity from $G(t)$** **32**

**S8 Simulation Results** **34**

S8.1 The Effects of the Genetic Architecture under Exponential Directional Selection . . 34

### S1 Estimator Bias Correction

#### S1.1 Correcting variance bias with a single depth sampling process

Following Waples (1989), we have that the variance in allele frequency change at a locus in the initial generation, which is entirely due to a binomial sampling process, is $\text{Var}(p_0) = {}^{p_0(1-p_0)}/_{d_0}$ where $d_0$ is the number of binomial draws (e.g. read depth). At a later timepoint, the variance in allele frequency is a result of both the binomial sampling process at time $t$ and the evolutionary process. Using the law of total variation we can partition the variation from each process,

$$\text{Var}(\widetilde{p_t}) = \mathbb{E}(\text{Var}(\widetilde{p_t}|p_t)) + \text{Var}(\mathbb{E}(\widetilde{p_t}|p_t)) \tag{S1}$$

$$= \underbrace{\frac{p_t(1-p_t)}{d_t}}_{\text{generation } t \text{ sampling noise}} + \underbrace{\text{Var}(p_t)}_{\text{variance due to evolutionary process}}. \tag{S2}$$

Under a drift-only process, $\text{Var}(p_t) = p_0(1-p_0)\left[1 - \left(1 - \frac{1}{2N}\right)^t\right]$. However, with heritable variation in fitness, we need to consider the covariance in allele frequency changes across generations (Buffalo and Coop 2019). We can write

$$\text{Var}(p_t) = \text{Var}\left(p_0 + (p_1 - p_0) + (p_2 - p_1) + \ldots + (p_t - p_{t-1})\right) \tag{S3}$$

$$= \text{Var}\left(p_0 + \Delta p_0 + \Delta p_1 + \ldots + \Delta p_{t-1}\right) \tag{S4}$$

$$= \text{Var}(p_0) + \sum_{i=0}^{t-1} \text{Cov}(p_0, \Delta p_i) + \sum_{i=0}^{t-1} \text{Var}(\Delta p_i) + \sum_{0 \le i < j}^{t-1} \text{Cov}(\Delta p_i, \Delta p_j). \tag{S5}$$

Each allele frequency change is equally like to be positive as it is to be negative; thus by symmetry this second term is zero. Additionally $\text{Var}(p_0) = 0$, as we treat $p_0$ as a fixed initial frequency. We can write,

$$\text{Var}(p_t) = \sum_{i=0}^{t-1} \text{Var}(\Delta p_i) + \sum_{0 \le i < j}^{t-1} \text{Cov}(\Delta p_i, \Delta p_j). \tag{S6}$$

The second term, the cumulative impact of variance in allele frequency change can be partitioned into heritable fitness and drift components (Buffalo and Coop 2019; Santiago and Caballero 1995)

$$\text{Var}(p_t) = \sum_{i=0}^{t-1} \text{Var}(\Delta_D p_i) + \sum_{i=0}^{t-1} \text{Var}(\Delta_H p_i) + \sum_{0 \le i < j}^{t-1} \text{Cov}(\Delta p_i, \Delta p_j). \tag{S7}$$

where $\Delta_H p_t$ and $\Delta_D p_t$ indicate the allele frequency change due to heritable fitness variation and drift respectively. Then, sum of drift variances in allele frequency change is

$$\sum_{i=0}^{t-1} \text{Var}(\Delta_D p_i) = \sum_{i=0}^{t-1} \frac{p_i(1-p_i)}{2N} \tag{S8}$$

replacing the heterozygosity in generation $i$ with its expectation, we have

$$\sum_{i=0}^{t-1} \mathrm{Var}(\Delta_D p_i) = p_0(1-p_0) \sum_{i=0}^{t-1} \frac{1}{2N}\left(1-\frac{1}{2N}\right)^i \tag{S9}$$

$$= p_0(1-p_0)\left[1-\left(1-\frac{1}{2N}\right)^t\right] \tag{S10}$$

which is the usual variance in allele frequency change due to drift. Then, the total allele frequency change from generations 0 to $t$ is $\mathrm{Var}(\widetilde{p}_t - \widetilde{p}_0) = \mathrm{Var}(\widetilde{p}_t) + \mathrm{Var}(\widetilde{p}_0) - 2\,\mathrm{Cov}(\widetilde{p}_t, \widetilde{p}_0)$, where the covariance depends on the nature of the sampling plan (see Nei and Tajima 1981; Waples 1989). In the case where there is heritable variation for fitness, and using the fact that $\mathrm{Cov}(\widetilde{p}_t, \widetilde{p}_0) = {}^{p_0(1-p_0)}/_{2N}$ for Plan I sampling procedures (Waples 1989), we write,

$$\mathrm{Var}(\widetilde{p}_t - \widetilde{p}_0) = \mathrm{Var}(\widetilde{p}_t) + \mathrm{Var}(\widetilde{p}_0) - 2C\,\mathrm{Cov}(\widetilde{p}_t, \widetilde{p}_0) \tag{S11}$$

$$= \frac{p_t(1-p_t)}{d_t} + \frac{p_0(1-p_0)}{d_0} + p_0(1-p_0)\left[1-\left(1-\frac{1}{2N}\right)^t\right] + \tag{S12}$$

$$\sum_{i=0}^{t-1} \mathrm{Var}(\Delta_H p_i) + \sum_{0\le i<j}^{t-1} \mathrm{Cov}(\Delta p_i, \Delta p_j) - \frac{C p_0(1-p_0)}{2N} \tag{S13}$$

$$\frac{\mathrm{Var}(\widetilde{p}_t - \widetilde{p}_0)}{p_0(1-p_0)} = 1 + \frac{p_t(1-p_t)}{p_0(1-p_0)d_t} + \frac{1}{d_0} - \left(1-\frac{1}{2N}\right)^t + \tag{S14}$$

$$\sum_{i=0}^{t-1} \frac{\mathrm{Var}(\Delta_H p_i)}{p_0(1-p_0)} + \sum_{0\le i<j}^{t-1} \frac{\mathrm{Cov}(\Delta p_i, \Delta p_j)}{p_0(1-p_0)} - \frac{C}{N} \tag{S15}$$

where $C = 1$ if Plan I is used, and $C = 0$ if Plan II is used (see Waples 1989, p. 380 and Figure 1 for a description of these sampling procedures; throughout the paper we use sampling Plan II). Rearranging, we can create a bias-corrected estimator for the population variance in allele frequency change, and replace all population heterozygosity terms with the unbiased sample estimators, e.g. $\frac{d_t}{d_t-1}\widetilde{p}_t(1-\widetilde{p}_t)$,

$$\frac{d_0-1}{d_0}\frac{\mathrm{Var}(\widetilde{p}_1 - \widetilde{p}_0)}{\widetilde{p}_0(1-\widetilde{p}_0)} - \frac{(d_0-1)}{d_0(d_1-1)}\frac{\widetilde{p}_1(1-\widetilde{p}_1)}{\widetilde{p}_0(1-\widetilde{p}_0)} - \frac{1}{d_0} + \frac{C}{N} = \frac{\mathrm{Var}(\Delta_H p_0)}{p_0(1-p_0)} + \frac{1}{2N} \tag{S16}$$

#### S1.2 Correcting variance bias with individual and depth sampling processes

Here, we extend the sampling bias correction described above to handle two binomial sampling processes: one as individuals are binomially sampled from the population, and another as reads are binomially sampled during sequencing. (see also Jónás et al. 2016). Let $X_t \sim \mathrm{Binom}(n_t, p_t)$ where $X_t$ is the count of alleles and $n_t$ is the number of diploids sampled at time $t$. Then, these individuals are sequenced at a depth of $d_t$, and $Y_t \sim \mathrm{Binom}(d_t, {}^{X_t}/_{n_t})$ reads have the tracked allele. We let $\widetilde{p}_t = {}^{Y_t}/_{d_t}$ be the observed sample allele frequency. Then, the sampling noise is

$$\mathrm{Var}(\widetilde{p}_t|p_t) = \mathbb{E}(\mathrm{Var}(\widetilde{p}_t|X_t)) + \mathrm{Var}(\mathbb{E}(\widetilde{p}_t|X_t)) \tag{S17}$$
$$= p_t(1-p_t)\left(\frac{1}{n_t} + \frac{1}{d_t} - \frac{1}{n_t d_t}\right) \tag{S18}$$

$$\mathrm{Var}(\widetilde{p}_t - \widetilde{p}_0) = p_t(1-p_t)\left(\frac{1}{n_t} + \frac{1}{d_t} - \frac{1}{n_t d_t}\right) + p_0(1-p_0)\left(\frac{1}{n_0} + \frac{1}{d_0} - \frac{1}{n_0 d_0}\right) \tag{S19}$$
$$- \frac{C p_0(1-p_0)}{N} + p_0(1-p_0)\left[1 - \left(1 - \frac{1}{2N}\right)^t\right] + \sum_{i=0}^{t-1} \mathrm{Var}(\Delta_{_H} p_i) \tag{S20}$$
$$+ \sum_{0 \le i < j}^{t-1} \mathrm{Cov}(\Delta p_i, \Delta p_j) \tag{S21}$$

Through the law of total expectation (see Kolaczkowski et al. 2011 Supplementary File 1 for a sample proof), one can find that an unbiased estimator of the half the heterozygosity is

$$\frac{n_t d_t}{(n_t-1)(d_t-1)} \widetilde{p}_t(1-\widetilde{p}_t). \tag{S22}$$

Replacing this unbiased estimator for half of the heterozygosity into our expression above, the total sample variance is

$$\mathrm{Var}(\widetilde{p}_t - \widetilde{p}_0) = \frac{n_t d_t \widetilde{p}_t(1-\widetilde{p}_t)}{(n_t-1)(d_t-1)}\left(\frac{1}{n_t} + \frac{1}{d_t} - \frac{1}{n_t d_t}\right) + \frac{n_0 d_0 \widetilde{p}_0(1-\widetilde{p}_0)}{(n_0-1)(d_0-1)}\left(\frac{1}{n_0} + \frac{1}{d_0} - \frac{1}{n_0 d_0}\right) + \tag{S23}$$

$$\frac{n_0 d_0 \widetilde{p}_0(1-\widetilde{p}_0)}{(n_0-1)(d_0-1)}\left[1 - \left(1 - \frac{1}{2N}\right)^t\right] - \frac{C}{N}\frac{n_0 d_0 \widetilde{p}_0(1-\widetilde{p}_0)}{(n_0-1)(d_0-1)} + \sum_{i=0}^{t-1} \mathrm{Var}(\Delta_{_H} p_i) + \sum_{0 \le i < j}^{t-1} \mathrm{Cov}(\Delta p_i, \Delta p_j). \tag{S24}$$

As with equation (S16), we can rearrange this to get a biased-corrected estimate of the variance in allele frequency change between adjacent generations, $\mathrm{Var}(\Delta p_t)$.

#### S1.3 Covariance Correction

We also need to apply a bias correction to the temporal covariances (and possibly the replicate covariances if the initial sample frequencies are all shared). The basic issue is that $\mathrm{Cov}(\Delta\widetilde{p}_t, \Delta\widetilde{p}_{t+1}) = \mathrm{Cov}(\widetilde{p}_{t+1} - \widetilde{p}_t, \widetilde{p}_{t+2} - \widetilde{p}_{t+1})$, and thus shares the sampling noise of timepoint $t+1$. Thus acts to bias the covariance by subtracting off the noise variance term of $\mathrm{Var}(\widetilde{p}_{t+1})$, so we add the expectation of this bias, derived above, back in. We discuss this in more detail below in deriving the bias correction for the temporal-replicate variance covariance matrix.

#### S1.4 Temporal-Replicate Covariance Matrix Correction

In practice, we simultaneously estimate the temporal and replicate covariance matrices for each replicate, which we call the temporal-replicate covariance matrix. This needs a bias correction; we extend the bias corrections for single locus variance and covariance described in Supplementary Material Sections S1.1, S1.2, and S1.3 to multiple sampled loci and the temporal-replicate covariance matrix here. With frequency data collected at $T+1$ timepoints across $R$ replicate populations at $L$ loci, we have multidimensional arrays $\mathbf{F}$ of allele frequencies, $\mathbf{D}$ of sequencing depths, and $\mathbf{N}$ of the number of individuals sequenced, each of dimension $R \times (T+1) \times L$. We calculate the array $\mathbf{\Delta F}$ which contains the allele frequency changes between adjacent generations, and has dimension $R \times T \times L$. The operation $\text{flat}(\mathbf{\Delta F})$ flattens this array to a $(R \cdot T) \times L$ matrix, such that rows are grouped by replicate, e.g. for timepoint $t$, replicate $r$, and locus $l$ such that for allele frequencies $p_{t,r,l}$, the frequency change entries are

$$\text{flat}(\mathbf{\Delta F}) = \begin{bmatrix} \Delta p_{1,0,0} & \Delta p_{2,0,0} & \cdots & \Delta p_{1,1,0} & \Delta p_{2,1,0} & \cdots & \Delta p_{T,R,0} \\ \Delta p_{1,0,1} & \Delta p_{2,0,1} & \cdots & \Delta p_{1,1,1} & \Delta p_{2,1,1} & \cdots & \Delta p_{T,R,1} \\ \vdots & \vdots & \ddots & \vdots & \vdots & \ddots & \vdots \\ \Delta p_{1,0,L} & \Delta p_{2,0,L} & \cdots & \Delta p_{1,1,L} & \Delta p_{2,1,L} & \cdots & \Delta p_{T,R,L} \end{bmatrix} \tag{S25}$$

where each $\Delta p_{t,r,l} = p_{t+1,r,l} - p_{t,r,l}$. Then, the sample temporal-replicate covariance matrix $\mathbf{Q}'$ calculated on $\text{flat}(\mathbf{\Delta F})$ is a $(R{\cdot}T)\times(R{\cdot}T)$ matrix, with the $R$ temporal-covariance block submatrices along the diagonal, and the $R(R-1)$ replicate-covariance submatrices matrices in the upper and lower triangles of the matrix,

$$\mathbf{Q}' = \begin{bmatrix} \mathbf{Q}'_{1,1} & \mathbf{Q}'_{1,2} & \cdots & \mathbf{Q}'_{1,R} \\ \mathbf{Q}'_{2,1} & \mathbf{Q}'_{2,2} & \cdots & \mathbf{Q}'_{2,R} \\ \vdots & \vdots & \ddots & \vdots \\ \mathbf{Q}'_{R,1} & \mathbf{Q}'_{R,2} & \cdots & \mathbf{Q}'_{R,R} \end{bmatrix} \tag{S26}$$

where each submatrix $\mathbf{Q}'_{i,j}$ $(i \neq j)$ is the $T \times T$ sample replicate covariance matrix for replicates $i$ and $j$, and the submatrices along the diagonal $\mathbf{Q}'_{r,r}$ are the temporal covariance matrices for replicate $r$.

Given the bias of the sample covariance of allele frequency changes, we calculated an expected bias matrix $\mathbf{B}$, averaging over loci,

$$\mathbf{B} = \frac{1}{L}\sum_{l=1}^{L} \frac{\mathbf{h}_l}{2} \circ \left( \frac{1}{\mathbf{d}_l} + \frac{1}{2\mathbf{n}_l} + \frac{1}{2\mathbf{d}_l \circ \mathbf{n}_l} \right) \tag{S27}$$

where $\circ$ denotes elementwise product, and $\mathbf{h}_l$, $\mathbf{d}_l$, and $\mathbf{n}_l$, are rows corresponding to locus $l$ of the unbiased heterozygosity arrays $\mathbf{H}$, depth matrix $\mathbf{D}$, and number of diploids matrix $\mathbf{N}$. The unbiased $R \times (T+1) \times L$ heterozygosity array can be calculated as

$$\mathbf{H} = \frac{2\mathbf{D} \circ \mathbf{N}}{(\mathbf{D}-1) \circ (\mathbf{N}-1)} \circ \mathbf{F} \circ (1 - \mathbf{F}) \tag{S28}$$

where division here is elementwise. Thus, $\mathbf{B}$ is a $R \times (T+1)$ matrix. As explained in Supplementary Material Section S1.2 and S1.3, the temporal variances and covariances require bias corrections, meaning each temporal covariance submatrix $\mathbf{Q}_{r,r}$ requires two corrections. For an element $Q_{r,t,s} = \text{Cov}(\Delta p_t, \Delta p_s)$ of the temporal covariance submatrix for replicate $r$, $\mathbf{Q}_{r,r}$, we apply the following correction

$$Q_{r,t,s} = \begin{cases} Q'_{r,t,s} - b_{r,t} - b_{r,t+1}, & \text{if } t = s \\ Q'_{r,t,s} + b_{r,\max(t,s)}, & \text{if } |t-s| = 1 \end{cases} \tag{S29}$$

where $b_{r,t}$ is element in row $r$ and column $t$ of $\mathbf{B}$.

### S2 Barghi et al. (2019) Temporal Covariances

Since each replicate population was sequenced every ten generations, the timepoints $t_0 = 0$ generations, $t_1 = 10$ generations, $t_2 = 20$ generations, etc., lead to observed allele frequency changes across ten generation blocks, $\Delta p_{t_0}, \Delta p_{t_1}, \ldots, \Delta p_{t_6}$. Consequently, the ten temporal covariance matrices for each of the ten replicate populations have off-diagonal elements of the form $\text{Cov}(\Delta p_{t_0}, \Delta p_{t_1}) = \text{Cov}(p_{t_1} - p_{t_0}, p_{t_2} - p_{t_1}) = \sum_{i=0}^{10} \sum_{j=10}^{20} \text{Cov}(\Delta p_i, \Delta p_j)$. Each diagonal element has the form $\text{Var}(\Delta p_{t_0}) = \sum_{i=0}^{t_0-1} \text{Var}(\Delta p_i) + \sum_{i=0}^{t_0-1} \sum_{j \neq i}^{t_0-1} \text{Cov}(\Delta p_i, \Delta p_j)$, and is thus a combination of the effects of drift and selection, as both the variance in allele frequency changes and cumulative temporal autocovariances terms increase the variance in allele frequency. With sampling each generation, one could more accurately partition the total variance in allele frequency change (Buffalo and Coop 2019); while we cannot directly estimate the contribution of linked selection to the variance in allele frequency change here, the presence of a positive observed covariance between allele frequency change can only be caused linked selection.

### S3 Block Bootstrap Procedure

The estimators used in this paper are predominantly ratios, e.g. temporal-replicate covariance standardized by half the heterozygosity, $G(t)$ which is the ratio of cumulative covariance to total variance, and the convergence correlation (equation (2)). In these cases, we can exploit the linearity of the expectation to make the bootstrap procedure more computationally efficient, by pre-calculating the statistics of the ratio's numerator and denominator, $N(\mathbf{x}_i)$ and $D(\mathbf{x}_i)$, on the data $\mathbf{x}_i$ for all blocks $i \in \{1, 2, \ldots, W\}$ in the genome. Then we draw $W$ bootstrap samples with replacement, and compute the estimate for bootstrap sample $b$ with an average weighted by the fraction $w_i$ of total loci contained in each block,

$$\tilde{\theta}_b = \frac{\sum_{i=1}^{W} w_i N(\mathbf{x}_i)}{\sum_{i=1}^{W} w_i D(\mathbf{x}_i)} \tag{S30}$$

Note that computing the ratio of averages rather than the average of a ratio is a practice common for population genetic statistics like $F_{ST}$ (Bhatia et al. 2013). With these $B$ bootstrap estimates, we calculate the $\alpha/2$ and $1-\alpha/2$ quantiles, which we use to estimate the $1-\alpha = 95\%$ pivot confidence intervals (p. 33 Wasserman 2006, p. 194 Davison and Hinkley 2013) throughout the paper,

$$C_\alpha = \left(2\widehat{\theta} - q_{1-\alpha/2}, 2\widehat{\theta} - q_{\alpha/2}\right). \tag{S31}$$

where $\widehat{\theta}$ is the estimate, and $q_x$ is bootstrap quantile for probability $x$.

#### S4 Replicate $G(t)$ and Partitioning the Variance in Allele Frequency

We define a statistic similar to $G(t)$ for estimating the proportion of allele frequency change common between two replicate populations due to linked selection. Covariance in allele frequency change between two replicate populations is due to convergent selection pressure selecting haplotypes shared between the two replicate populations, which acts to perturb linked neutral variation in parallel way.

$$G_R(t) = \frac{\mathbb{E}_{A \neq B}(\sum_{i \neq j}^{t} \text{Cov}(\Delta p_{i,A}, \Delta p_{j,B}))}{\mathbb{E}_R(\text{Var}(p_{t,R} - p_{0,R}))} \tag{S32}$$

where $\mathbb{E}_{A \neq B}$ indicates that the expectation is taken over all ordered pairs of replicates (e.g. summing all off-diagonal elements replicate covariances), and $\mathbb{E}_R$ indicates taking expectation over all replicates. This measures the fraction of variance in allele frequency change (averaged across replicates) due to shared selection pressure.

Extending our theoretic work in Buffalo and Coop (2019), we can partition the allele frequency change in two replicates into drift, and shared selection and replicate-specific selection components of allele frequency change. For two replicates, $A$ and $B$,

$$\Delta p_{t,A} = \Delta_D p_{t,A} + \Delta_U p_{t,A} + \Delta_S p_t \tag{S33}$$

$$\Delta p_{t,B} = \Delta_D p_{t,B} + \Delta_U p_{t,B} + \Delta_S p_t \tag{S34}$$

where $\Delta_D p_{t,A}$ is allele frequency change due to drift (this is specific to a replicate, and equal to $\Delta_N p_{t,A} + \Delta_M p_{t,A}$ in the notation of Buffalo and Coop 2019), $\Delta_U p_{t,A}$ is the allele frequency change from indirect selection specific to replicate $A$ (and likewise with $\Delta_U p_{t,A}$ for replicate $B$), and $\Delta_S p_t$ is the allele frequency change from indirect selection shared across the replicates $A$ and $B$ (this term lacks a replicate subscript since by construction it is identical between replicates). By construction, each of these terms is uncorrelated, so the variance and be written as:

$$\text{Var}(\Delta p_{t,A}) = \text{Var}(\Delta_D p_{t,A}) + \text{Var}(\Delta_U p_{t,A}) + \text{Var}(\Delta_S p_t) \tag{S35}$$

(S36)

The shared effects of indirect selection can be quantified from the observed allele frequency changes, since the covariance in allele frequency change across replicates is the covariance of the shared term by construction,

$$\mathrm{Cov}(\Delta p_{t,A}, \Delta p_{t,B}) = \mathrm{Cov}(\Delta_S p_t, \Delta_S p_t) = \mathrm{Var}(\Delta_S p_t) \tag{S37}$$

In artificial selection studies with a control (non-selected) line, such as the Castro et al. (2019) study, this allows us to estimate the contribution of the effects of shared and unique indirect selection. In the case of this study, we can estimate the drift, unique selection effect, and shared selection effect terms using the fact that,

$$\Delta p_{t,\mathrm{LS1}} = \Delta_D p_{t,\mathrm{LS1}} + \Delta_U p_{t,\mathrm{LS1}} + \Delta_{\mathrm{LS}} p_t \tag{S38}$$
$$\Delta p_{t,\mathrm{LS2}} = \Delta_D p_{t,\mathrm{LS2}} + \Delta_U p_{t,\mathrm{LS2}} + \Delta_{\mathrm{LS}} p_t \tag{S39}$$
$$\Delta p_{t,\mathrm{Ctrl}} = \Delta_D p_{t,\mathrm{Ctrl}}. \tag{S40}$$

Note that since the control replicate does not undergo artificial selection, we assume that its allele frequency changes are determined entirely by genetic drift. With free mating individuals (such as in a cage population), this may not be the case, and sequencing adjacent generations would allow one to differentiate the effects of selection and drift.

We assume that we can approximate the contribution of genetic drift in the Longshanks selection lines with the observed variance in the control line, or $\mathrm{Var}(\Delta p_{t,\mathrm{Ctrl}}) = \mathrm{Var}(\Delta_{D p_{t,\mathrm{LS1}}}) = \mathrm{Var}(\Delta_{D p_{t,\mathrm{LS2}}})$. Then, the combined effects of selection can be estimated by averaging the variances of the two Longshanks selection lines, and subtracting the variance in allele frequency change in the control line, which we treat as driven by drift alone (since matings are random). Note that each variance is bias-corrected according to the methods described in Supplementary Materials S1.4, and the average sequencing depths between lines are nearly identical. Thus, we have

$$(\mathrm{Var}(\Delta p_{t,\mathrm{LS1}}) + \mathrm{Var}(\Delta p_{t,\mathrm{LS2}}))/2 - \mathrm{Var}(\Delta p_{t,\mathrm{Ctrl}}) = \overline{\mathrm{Var}(\Delta_U p_{t,\mathrm{LS}})} + \mathrm{Var}(\Delta_{\mathrm{LS}} p_t) \tag{S41}$$

where the bar indicates values averaged both Longshanks selection lines. Additionally, use the fact that

$$\mathrm{Cov}(\Delta p_{t,\mathrm{LS1}}, \Delta p_{t,\mathrm{LS2}}) = \mathrm{Var}(\Delta_{\mathrm{LS}} p_t) \tag{S42}$$

we can also separate out the unique and shared components by subtracting off this covariance,

$$\overline{\mathrm{Var}(\Delta_U p_{t,\mathrm{LS}})} = (\mathrm{Var}(\Delta p_{t,\mathrm{LS1}}) + \mathrm{Var}(\Delta p_{t,\mathrm{LS2}}))/2 - \mathrm{Var}(\Delta p_{t,\mathrm{Ctrl}}) - \mathrm{Cov}(\Delta p_{t,\mathrm{LS1}}, \Delta p_{t,\mathrm{LS2}}). \tag{S43}$$

It is unclear how strong the fluctuations would have to be to generate a genome-wide average signal of fluctuating selection from temporal covariances. For example, many loci could still show

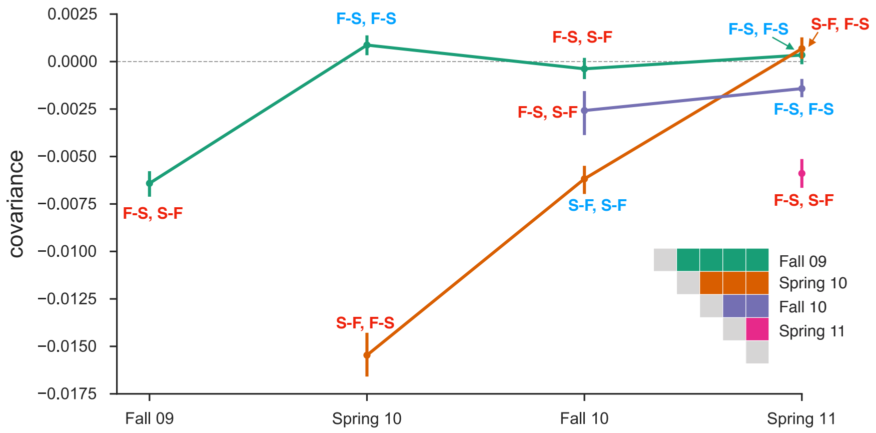

**Figure S1:** Temporal covariances from the Bergland et al. (2014) study, from varying reference generations (e.g. rows along the temporal covariance matrix). Each covariance is labeled indicating whether the covariance is between two like seasonal transitions (e.g. the covariance between allele frequency changes from fall to spring in one year, and fall to spring in another) or two dislike seasons (e.g. the covariance between fall to spring in one year, and spring to fall in another year). Covariances between like transitions are expected to be positive when there is a genome-wide effect of fluctuating selection (and these labels are colored blue), while covariances between dislike transitions are expected to be negative (and these labels are colored red). 95% confidence intervals were constructed by a block-bootstrapping procedure where the blocks are megabase tiles.

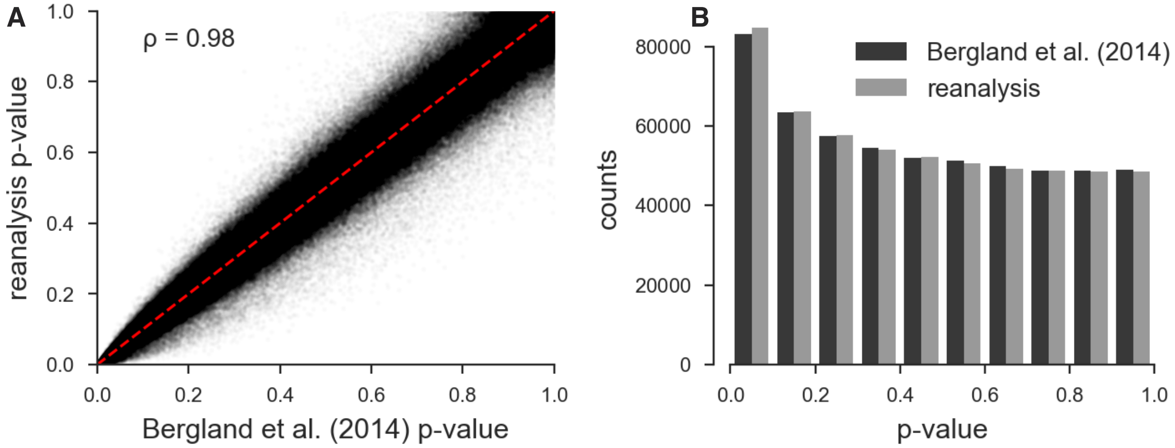

**Figure S2:** **A**: Scatterplot of the original unadjusted p-values from Bergland et al. (2014) and the p-values from our reanalysis of the same data using the same statistical methods; the minor discrepancy is likely due to software version differences. **B**: The histograms of the p-values of our reanalysis and the original Bergland et al. (2014) data; again the minor discrepancy is likely due to software differences. Overall, our implementation of Bergland et al.'s statistical methods produces results very close to the original analysis.

a signal of fluctuating selection, but the average signal could be overwhelmed by other signals of other selection. To investigate whether there was a genome-wide excess of loci showing evidence of fluctuating selection we reanalyzed the data of (Bergland et al. 2014) using the same seasonal fluctuating model as the original paper. This model is a Binomial logit-linked GLM fit per-locus, where the frequencies are regressed on the Spring/Fall seasons are encoded as a dummy variable. We use the same binomial weighting procedure as Bergland et al. (2014), where the weights are determined by the effective number of chromosomes, $N_{eff} = (2n_t d_t - 1)/(2n_t + d_t)$ ($n_t$ and $d_t$ are the number of diploid individuals and the read depth at timepoint $t$, respectively). We fit this model on all loci marked as used in the VCF provided with the Bergland et al. (2014) study (doi:10.5061/dryad.v883p). Overall, our p-values for the Wald test for each locus closely match those of the original paper (Pearson correlation coefficient 0.98, p-value $< 2.2 \times 10^{-16}$; see Supplementary Figure S2 A), and the histograms of the p-values are nearly identical (Supplementary Figure S2 B). Bergland et al. (2014) find loci with a significant association with season after a Benjamini and Hochberg FDR p-value adjustment (Benjamini and Hochberg 1995), however, the null hypothesis of the Wald test does not give us an idea of the expected number of variants that may spuriously fit the pattern of seasonal fluctuating selection as it does not account for genetic drift or other forms of hitchhiking.

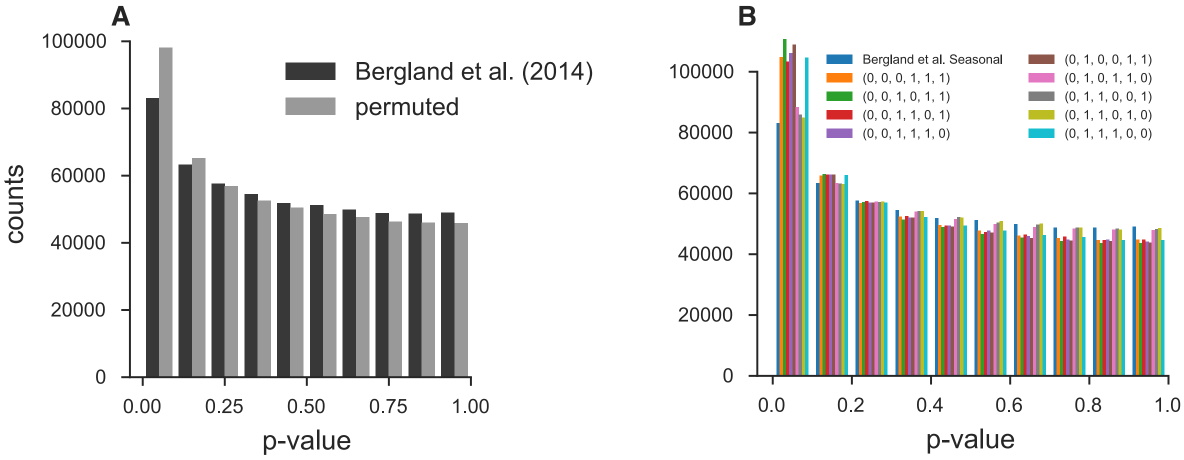

**Figure S3:** **A**: Histogram of original Bergland et al. (2014) seasonal p-values and p-values creating by randomly permuting the seasons at each locus. **B**: Histogram of original Bergland et al. (2014) p-values alongside all unique permutations (ignoring symmetries that lead to identical p-values).

To investigate whether there is a genome-wide evidence of an enrichment of fluctuating selection we created an empirical null distribution by randomly permuting the season labels and re-running the per-locus seasonal GLM model, as proposed by Machado et al. (2018). We find, regardless of whether we permute at the locus-level or the permutation replicate-level, that the observed seasonal p-value distribution Bergland et al. (2014) is not enriched for significant p-values beyond what we would expect from the permutation null. In fact, there appears there is more enrichment for low p-values when seasonal labels are randomly permuted (Supplementary Figure S3, suggesting by random chance we might expect more variants with a seasonal fluctuating pattern than found in the original Bergland et al. (2014) study. While surprising, this could be explained by the presence of temporal structure across the samples not consistent with seasonal fluctuating selection. Some

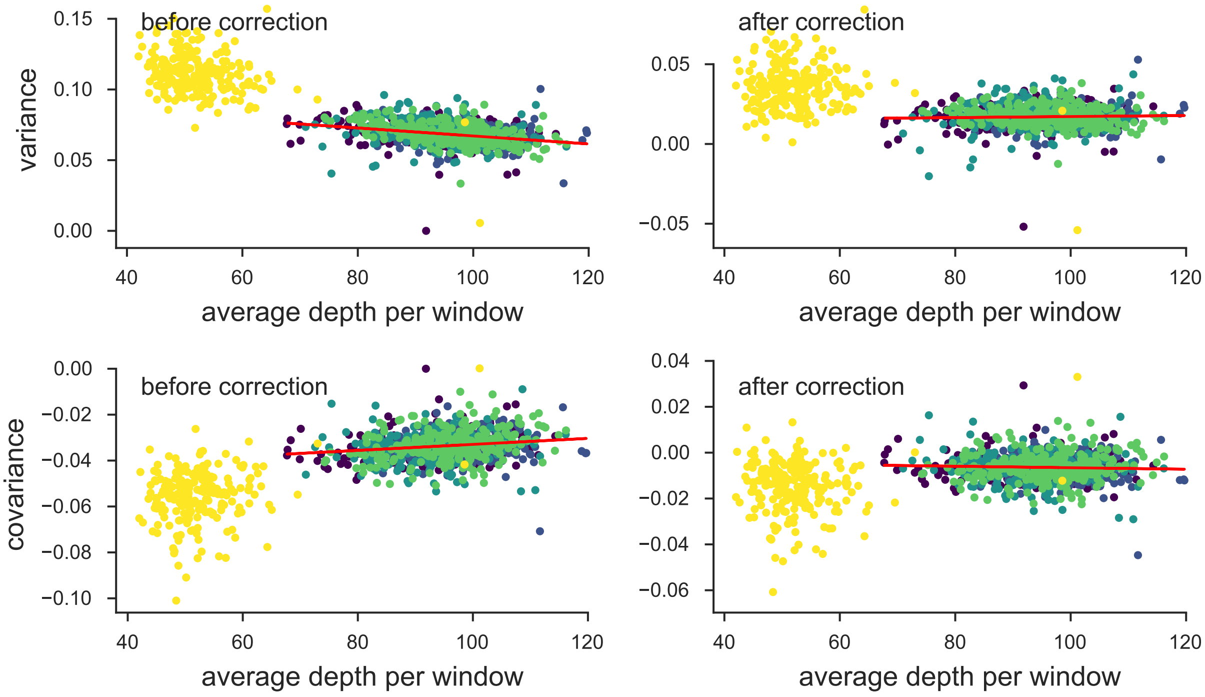

**Figure S4:** The variance and covariances from the Bergland et al. (2014) study, calculated in 100kb genomic windows plotted against average depth in a window before and after bias correction. Each panel has a least-squares estimate between the variance and covariance, and the average depth. The bias correction procedure is correcting sampling bias in both the variance and covariance such that the relationship with depth is constant. Colors indicate the different chromosomes of *D. melanogaster*; we have excluded the X chromosome (yellow points; chromosome 4 was not in the original study) from the regression due to large differences in average coverage.

fraction of the permutations happen to fit non-seasonal structure well, leading to an enrichment of small p-values. We note that genetic drift is not accounted for in the model used to estimate the significance of seasonally fluctuations and so some of these issues from non-seasonal structure may be due to a poorly calibrated null model. Furthermore, non-seasonal temporal structure is also evident in our temporal covariances (Supplementary Figure S1), where we see strong evidence of selection (non-zero temporal covariances), yet the pattern does not follow that of seasonal fluctuating selection.

### S7 Approximating the Reduction in Diversity from $G(t)$

To help reconcile our measure of linked selection, $G(t)$, with familiar expressions as a reduction in neutral diversity, as parameterized by $N_e$, we develop some rough approximations here. Note, however, that since linked selection generates temporal covariance, the overall effect is qualitatively different than just a simple reduction in $N_e$, as drift alone cannot generate temporal covariances. First, we introduce some convenient notation. Let $V_T = \text{Var}(p_t - p_0)$ be the total observed ~~variation~~ variance in allele frequency, $C_{LS} = \sum_{i=0}^{t-1} \sum_{j \neq i}^{t-1} \text{Cov}(\Delta p_i, \Delta p_j)$ be the contribution of all pairwise temporal covariances (observable), $V_D$ be the unobservable drift-only variance in allele frequency, and $V_{LS} = \sum_{i=0}^{t-1} \text{Var}(\Delta_{LS} p_i)$ which is the (unobservable) effect linked selection has on the variances in allele frequency change. Then,

$$V_T = V_{LS} + C_{LS} + V_D. \tag{S44}$$

Our measure $G(t)$ is then,

$$G(t) = \frac{C_{LS}}{V_T}, \tag{S45}$$

meaning we can express the fraction of total variance due to drift alone as

$$\begin{aligned}
\frac{V_D}{V_T} &= 1 - G(t) - \frac{V_{LS}}{V_T} && \text{(S46)}\\
&= 1 - G(t) - \varepsilon && \text{(S47)}\\
&\leq 1 - G(t), && \text{(S48)}
\end{aligned}$$

where $\varepsilon \geq 0$ since these variances are positive. Throughout this section, for convenience, we assume that the covariances contributing to $C_{LS}$, and thus $G(t)$, are all positive.

Rather than expressing this relationship in terms of variances in allele frequencies, we can express the same relationship in terms of $N_e$. In a neutral Wright–Fisher population, the total variation in allele frequency change over $t$ generations is

$$V_T = \text{Var}(p_t - p_0) = p_0(1 - p_0)\left[1 - (1 - 1/2N_e)^t\right]. \tag{S49}$$

For small $t$, a common temporal estimator for the variance effective population size $N_e$ is,

$$N_e \approx \frac{t p_0(1 - p_0)}{2V_T}. \tag{S50}$$

Then, the drift-only $N_e$ estimate (that is, removing the effects of linked selection) replaces the observable $V_T$ with unobservable $V_D$, and uses the $G(t)$ estimate to bound this:

$$\begin{aligned}
N_{e,D} &\approx \frac{t p_0(1 - p_0)}{2V_T(1 - G(t) - \varepsilon)} && \text{(S51)}\\
&\gtrsim \frac{t p_0(1 - p_0)}{2V_T(1 - G(t))} && \text{(S52)}\\
&\gtrsim \frac{N_e}{1 - G(t)} && \text{(S53)}\\
\frac{N_e}{N_{e,D}} &\lesssim 1 - G(t) && \text{(S54)}
\end{aligned}$$

In the linked selection literature, it is common to translate the impact of linked selection as a reduction in the level of neutral pairwise diversity in the absence of linked selection, $\pi_0$. Under the coalescent, $\pi_0 = 2\mu\mathbb{E}(T_2)$ where $\mathbb{E}(T_2)$ is the pairwise time to coalescence, which under drift alone in a constant population of size $N_e$, is $\mathbb{E}(T_2) = 2N$. The fraction of neutral diversity (in the absence of linked selection) reduced by selection is then, $1 - \bar{\pi}/\pi_0$, or equivalently, $1 - N_e/N_{e,D}$. Then,

$$G(t) \lesssim 1 - \frac{N_e}{N_{e,D}}, \tag{S55}$$

which shows that our measure $G(t)$ is a lower bound, over much shorter timescales, to the familiar measure $1 - \bar{\pi}/\pi_0$.

Elyashiv et al. (2016) estimated that linked selection in *Drosophila melanogaster* had resulted in a $1 - \bar{\pi}/\pi_0 = 77\%$ reduction in the level of neutral diversity. Thus our estimate of $G(t) \approx 20\%$ in *Drosophila simulans* is smaller than that seen over long timespans in a closely related species.

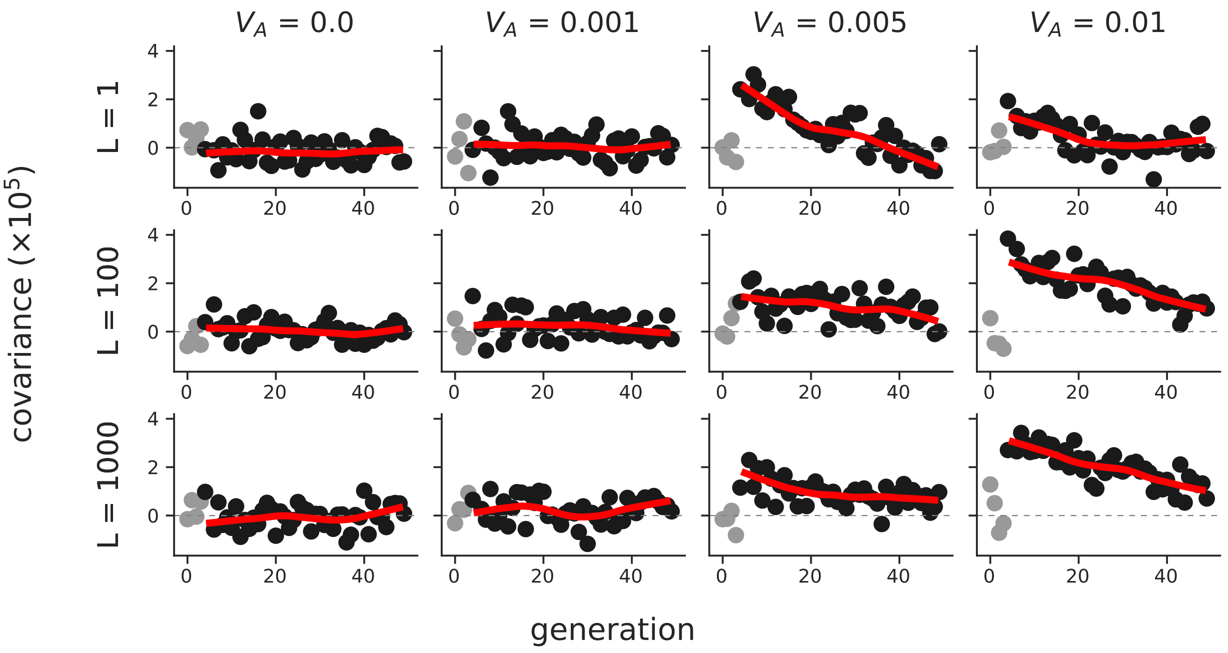

**Figure S5:** The temporal covariances $\text{Cov}(\Delta p_5, \Delta p_t)$ from the onset of selection (generation 5) to a later time point $t$, which varies along the x-axis, across a variety of different initial trait additive genetic variances ($V_A$, columns) and number of sites contributing to the trait ($L$, rows). Each point is the temporal covariance averaged over 50 replicate simulations; dark gray points are temporal covariances after the onset of selection, and light gray points are before. The red line is a loess-smoothed curve through the covariances after the onset of selection. Selection on the trait was imposed through an exponential fitness function.

During this burnin, sites were either marked as neutral (with mutation rate $\mu_{\text{neutral}} = 10^{-8}$ per gamete per generation) or contributed to the trait's value (with mutation rate $\mu_{\text{trait}}$), but were not selected until generation $10N + 5$ (the five generations after burnin serve as a neutral control). The trait mutation rate, $\mu_{\text{trait}}$ was set by targeting a particular architecture, the number of selected sites, $L$. Each site contributing to the trait's value was randomly chosen to have effect size $\pm\alpha$ with equal probability, where $\alpha$ was set to target a particular additive genetic variance for the trait, $V_A$, for the target number of selected sites $L$.

Overall, we ~~confirm a finding in Buffalo and Coop (2019)~~again see a finding of Buffalo and Coop (2019): that the initial expected temporal covariance conditioned on $V_A$, is invariant to the number of loci determining the trait's value, $L$ (Supplementary Figure S5). We do find some evidence that the decay in temporal covariance is faster when the trait has a monogenic basis (see the third column of Supplementary Figure S5); this is expected since the selection coefficients are larger for these monogenic simulations, leading to faster allele frequency changes and a rapid change in additive genetic variance.

In our previous work, we did not investigate the affect of trait architecture on our measure $G(t)$. Using the exponential fitness function simulations, we also calculated $G(t)$ for each of the replicate simulations. We find that the $G(t)$ trajectories can vary considerably across replicates depending on the number of sites ($L$) determining the trait's value (Supplementary Figure S6). When a trait is reasonably monogenic ($L \approx 1$), $G(t)$ trajectories vary considerably across replicate lines, as certain

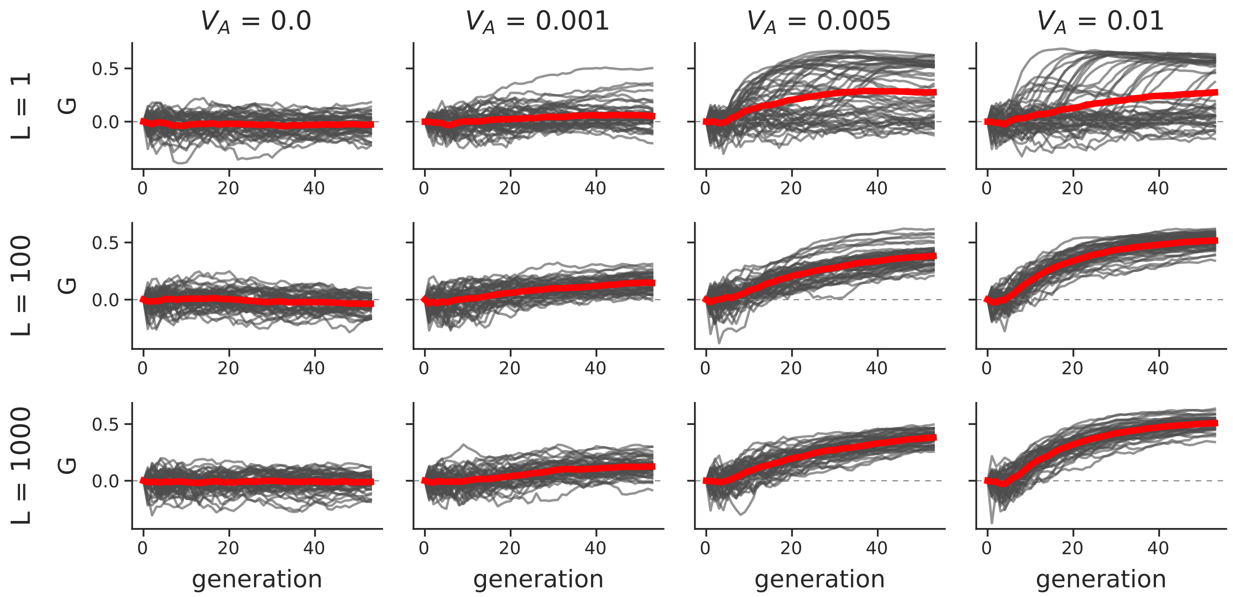

**Figure S6:** The $G(t)$ trajectories of 50 replicate simulations, across different trait architectures ($L$ is the target number of sites affecting the trait's value, and $V_A$ is the target trait additive genetic variance). The red line is the mean trajectory across all replicate simulations. Like Supplementary Figure S5, the onset of selection is five generations after the $10N$ generation burnin; this is evident by the initial flat period of the $G(t)$ trajectory.

lines may stochastically lose the few copies of the selected alleles (top row of Supplementary Figure S6). However, with a polygenic trait, ($L \geq 100$), the $G(t)$ trajectories across replicates are similar as each replicate contains an abundance of trait alleles (bottom rows of Supplementary Figure S6). Comparing the simulated $G(t)$ replicate trajectories of Supplementary Figure S6 with the Barghi et al. (2019) $G(t)$ trajectories in Figure 1B, we again confirm a finding of Barghi et al. (2019): that there is considerable genetic redundancy among beneficial alleles, meaning because of the polygenic architecture, there are multiple routes to adaptation. We should note that our simplified simulation routines are slightly different from the Barghi et al. (2019) study in that the burnin populations are all independent; however we expect the same qualitative result.

#### S8.2 Temporal Covariances and $G(t)$ under Gaussian Stabilizing Selection

Additionally, we wanted to ensure that our temporal covariances and $G(t)$ trajectories were robust to more complicated, but realistic fitness models. To this end, we also simulated Gaussian stabilizing selection (GSS) on a trait during burnin, followed by one of two ~~two optima~~ optimum shift routines: (1) sudden optima shifts of $\mu_{\text{sudden}} = \{0.1, 0.5, 1\}$, and (2) very graduate optima shifts of $\mu_{\text{gradual}} = \{0.001, 0.01\}$ per generation using the same two population simulation scheme described above (these shifts are in standard deviations, since $V_S = 1$). We used a polygenic architecture for these simulations, with trait alleles assigned a $\pm 0.01$ effect size with equal probability, trait mutation rate $10^{-8}$, and the optima shift began at five generations after a $10N$ generation burnin. Across our GSS simulations, we see the expected selection response (Supplementary Material Figure S7).

Overall, we see the same qualitative results under Gaussian stabilizing selection with optima

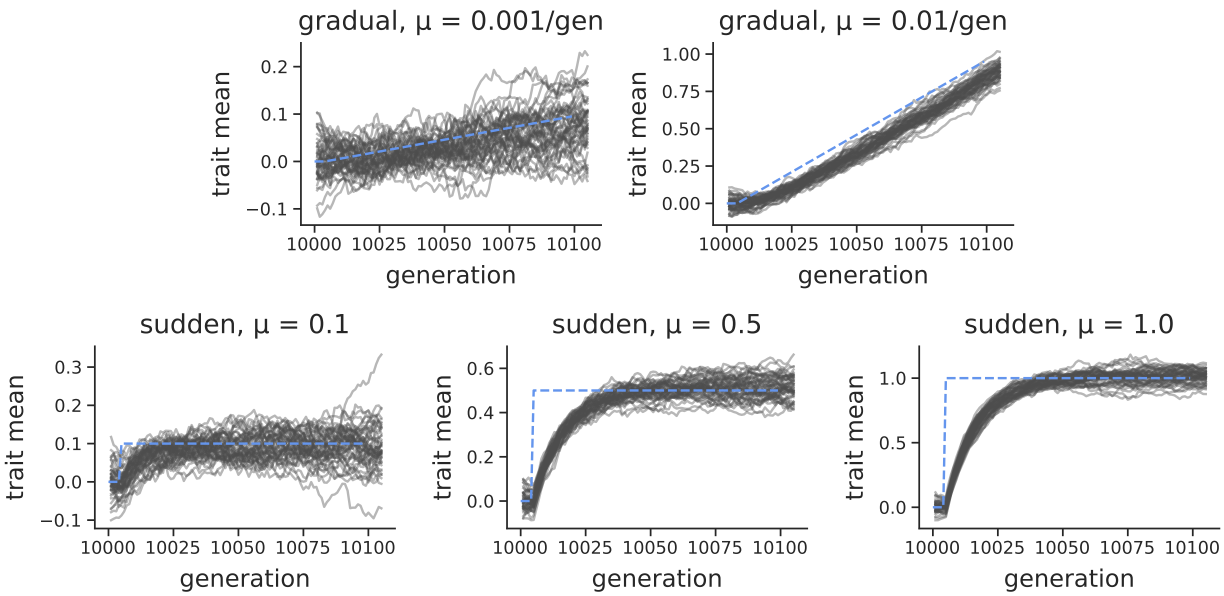

**Figure S7:** The population mean trait value under the Gaussian stabilizing selection simulations (gray lines) and the trait optima (dashed blue lines). The first row shows the selection response during a graduate shift in optima per generation, while the second row shows the selection response during a sudden optima shift.

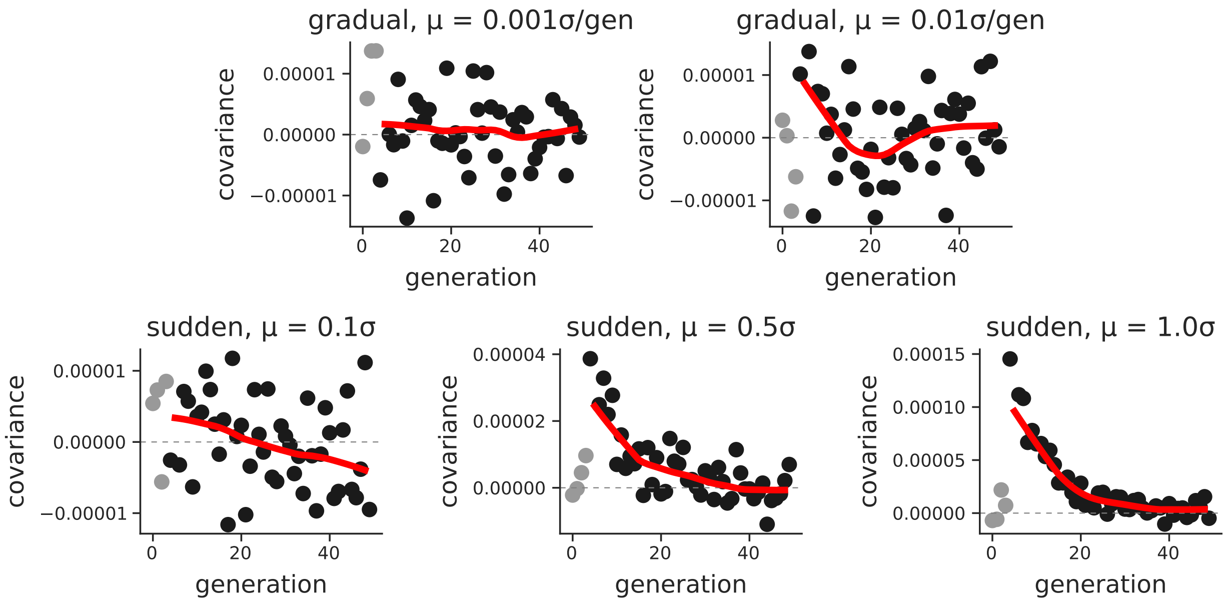

**Figure S8:** Mean temporal covariance ($\text{Cov}(\Delta p_5, \Delta p_t)$, with $t$ varying across the x-axis) across 30 replicate simulations (light gray points are before the onset of selection; dark gray points are after selection begins), under different Gaussian stabilizing selection with optima shift regimes. The solid red line is a loess-smoothed average of these points.

shifts as under exponential directional selection. Stronger directional selection, here determined by larger sudden optima shifts or larger gradual shifts per generation, lead to stronger temporal covariances (Supplementary Materials Figure S8). Furthermore, we see a stronger effect of linked selection, as measured by $G(t)$, under stronger directional selection (Supplementary Material Figure S9).

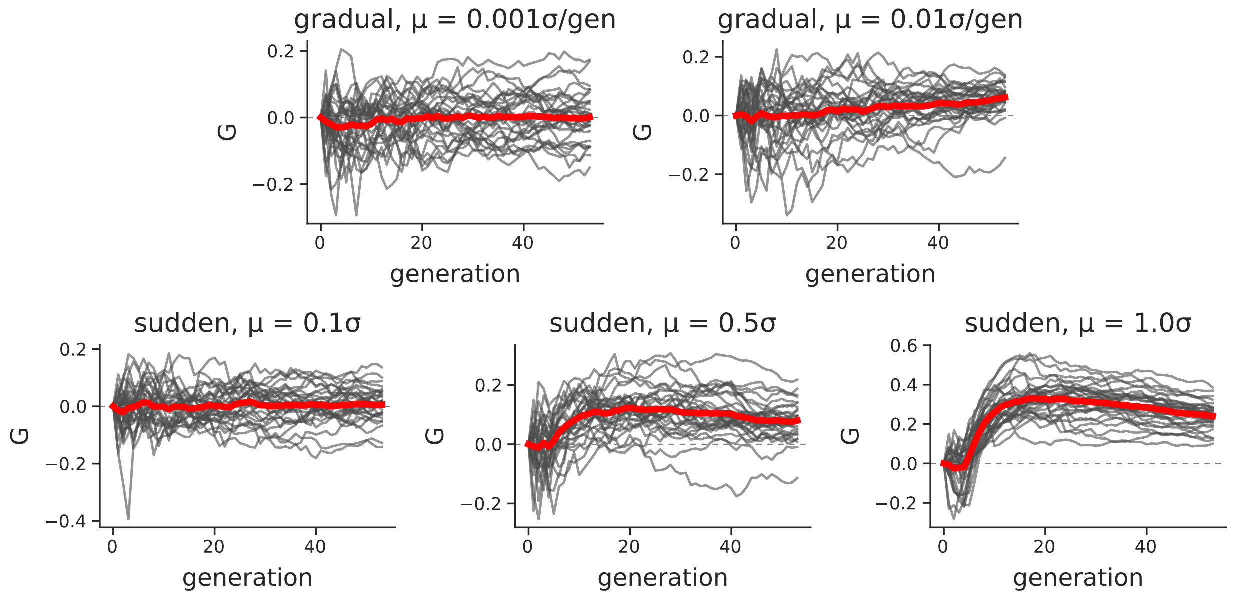

**Figure S9:** $G(t)$ trajectories across 30 replicate Gaussian stabilizing selection with optima shift regimes. The solid red line is a loess-smoothed average across replicates.

Additionally, we looked at the effect ~~the size of each replicate population~~ of the replicate population size drawn from the same population has on a single population's $G(t)$ trajectories. These simulations had the same $10N$ generation burnin, followed by a change in population size emulating the bottlenecks associated with creating selection lines. Overall, we find that smaller population sizes lead to a reduced $G(t)$ (Supplementary Material Figure S10). This is expected, as the denominator of $G(t)$ is $\text{Var}(p_t - p_0)$, which has an inverse relationship with $N_e$; as replicate population size is reduced, the proportion of allele frequency change driven by linked selection is lower, since the rate of drift is increased. To isolate the effects of varying replicate population size, we also looked at just the magnitude of temporal covariances (Supplementary Material Figure S11). We find that smaller replicate population sizes lead to *larger* temporal covariances. We then looked at the initial trait variance, which as expected, does not vary with replicate population size (since all burnin populations had the same size). This implies that the linkage disequilibria is higher in smaller populations, due to founder effects, which has the effect inflating the temporal covariance as predicted by our theory (Buffalo and Coop 2019).

##### S8.3 Convergence Correlations

Using the same exponential fitness function simulations described above, we also investigated how the convergence correlation is impacted by (1) genetic architecture, (2) the design of the selection experiment, e.g. how many individuals are selected for each line from the founding population, and

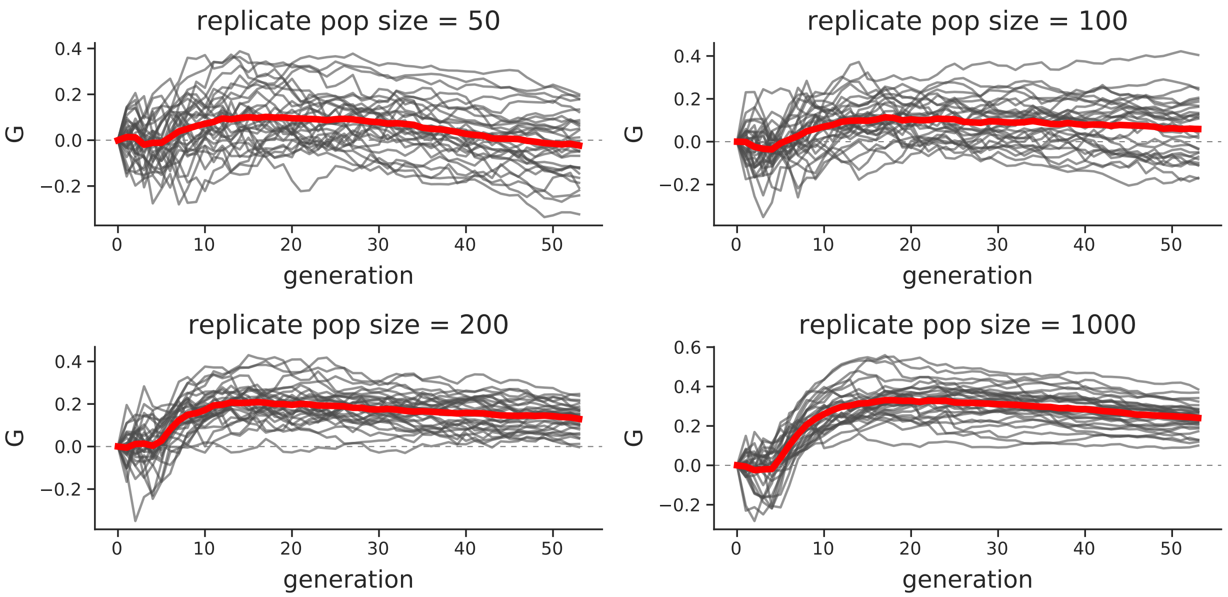

**Figure S10:** $G(t)$ trajectories under GSS after sudden optima shift of 1 at generation five, for varying replicate population sizes.

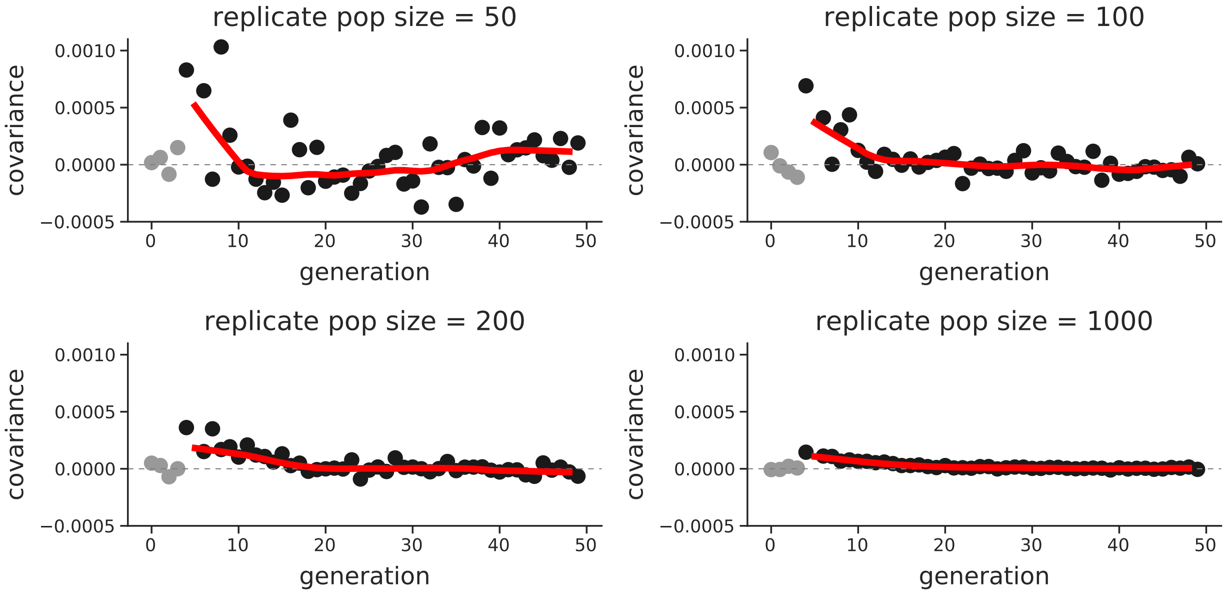

**Figure S11:** Temporal covariance $\text{Cov}(\Delta p_5, \Delta p_t)$, where $t$ varies along the x-axis, for a sudden optima shift of 1, for varying replicate population sizes. The reference time point is the first generation of selection; dark gray points are the temporal covariance after selection began, and the light gray points are before.

(3) the direction of selection across the two populations "lines". After burning in $N = 1000$ diploid populations for $10N$ generations, we simulated two equally-sized lines of sizes $n = \{50, 500, 1000\}$ diploids, and imposed three selection schemes across different simulation runs. First, we imposed a

convergent selection scheme, where the populations undergo exponential directional selection in the same direction. We expect that the convergent correlation under this convergent scheme should be positive, as the two lines should share some haplotypes carrying beneficial alleles, and these are selected in the same direction across the two lines. Second, we imposed divergent selection, where the two lines again undergo exponential directional selection, except in different directions. Here, we expect the convergence correlation to be negative, as haplotypes that increase the selected trait in one population are beneficial in the upward selected line, but deleterious in the downward selected line. Third, we have a control selection scheme, where one line is selected and the other is not; this is akin to the control line in the Castro et al. (2019) study (see Figure 2C). In this case, we expect to see no convergence correlation, as only one line is being selected. Finally, across these two-line simulation studies, we expect that smaller selection line sizes should show weaker convergent correlations, as the probability that the same haplotypes are selected between the two lines decreases with size.

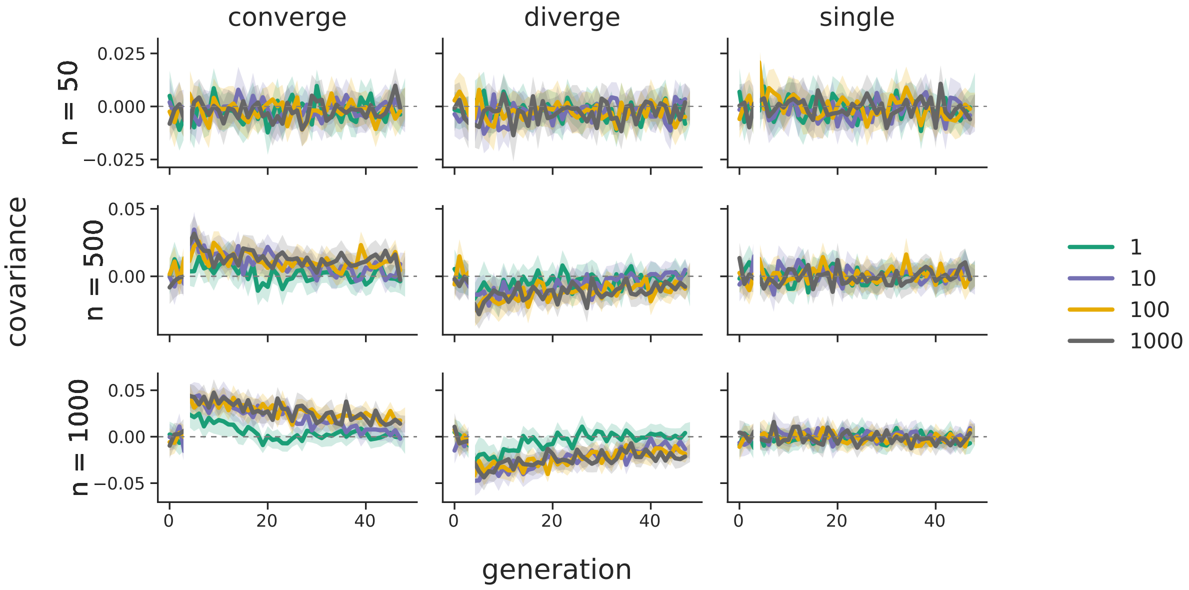

**Figure S12:** The convergence correlations across the two population line exponential directional selection simulations; panel rows are for differing line population sizes, and panel columns are the modes of selection across the lines (convergent, divergent, and only a single selected line control). Line color indicates the target genetic architecture, in number of loci affecting the trait's value. 95% confidence intervals are also shown. Note that selection begins at generation five, which is the reference generation; this is indicated by the split in the lines.

Overall, our simulations confirm our hypotheses; see Supplementary Material Figure S12. We also find that in simulations ~~where we target~~ with a monogenic genetic architecture (i.e. the target number of trait-affecting loci is $L = 1$), the convergence correlations are generally much weaker than those under a polygenic architecture. However, this effect is mediated by the line population size; the difference in convergence correlation between $L = 1$ and $L = 1000$ are more dissimilar when the line population sizes are larger (compare the first column, last two rows). Like the convergence correlations calculated on the Barghi et al. (2019) data, we find in simulations convergence correlations decay through time. Additionally, populations selected in opposite directions lead to negative convergence correlations, as expected. Overall, we find that the convergence correlation is affected by both genetic architecture and the size of the selected population lines.

We also wanted to test whether we see similar convergence correlations under Gaussian stabilizing selection. In these simulations, rather than targeting a particular $V_A$, we fix the trait mutation rate at $10^{-8}$ (thus region-wide $\theta = 2000$). Like the exponential directional selection simulations, we impose directional selection in the same direction across the two populations (converge), different directions (diverge), and only in one population (single). We also vary the type (gradual versus sudden) and magnitude of optima shifts in the two populations. Overall, simulations show convergence correlations for the sudden optima shifts in Supplementary Figure S13. Again, optima shifts in the opposite direction cause negative convergence correlations. We found that for slow moving optima shifts, the convergence correlations are generally too weak to be distinguished from zero reliably ~~, (top row of Supplementary Material Figure S13~~(not shown).

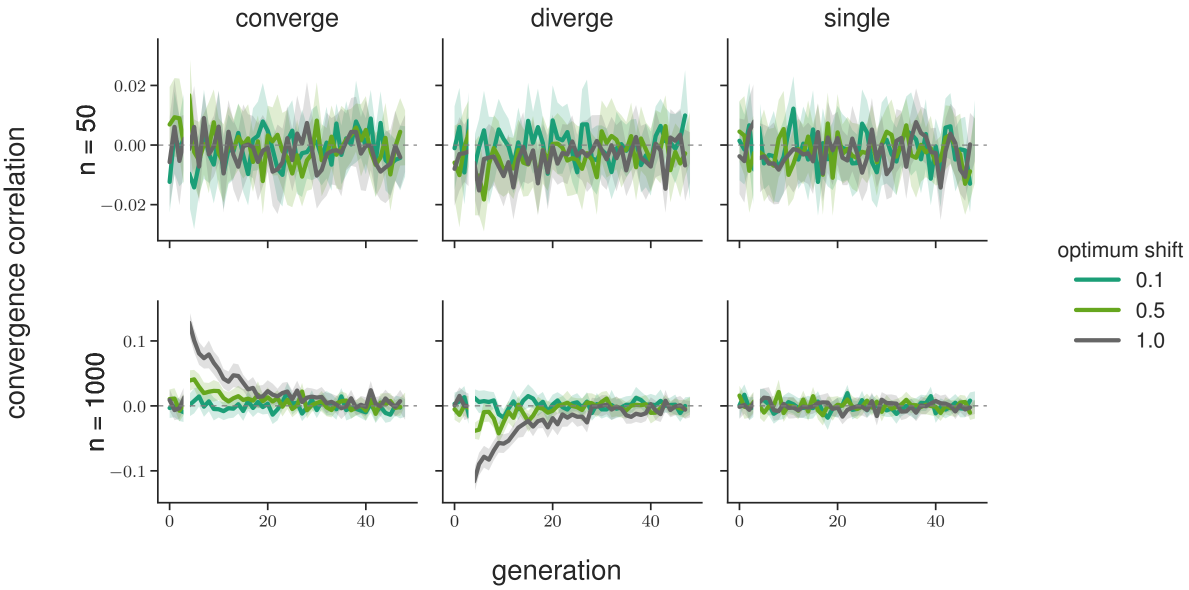

**Figure S13:** The convergence correlations across the two population line Gaussian stabilizing selection sudden optima shift simulations; selection line population sizes vary across rows, and panel columns are the modes of selection across the lines (convergent, divergent, and only a single selected line control). All simulations have a target number of loci affecting the trait of $L = 1000$; line color indicates the size of the sudden optima shift in standard deviations of $V_S$ 95% confidence intervals are also shown. Note that selection begins at generation five, which is the reference generation; this is indicated by the split in the lines.

#### S8.4 Sampling in Temporal Blocks

In our analysis of the Barghi et al. (2019) data, we describe our statistic $G(t)$ as a lower bound for two reasons: (1) the population is sequenced every ten generations, meaning the temporal covariances between adjacent generations cannot contribute to the numerator of $G(t)$ but contributes to the denominator, and (2) the estimate of $G(t)$ ignores linked selection's contribution to the per-generation variance in allele frequency change~~(this is the difference between our $G(t)$ and $G'(t)$~~

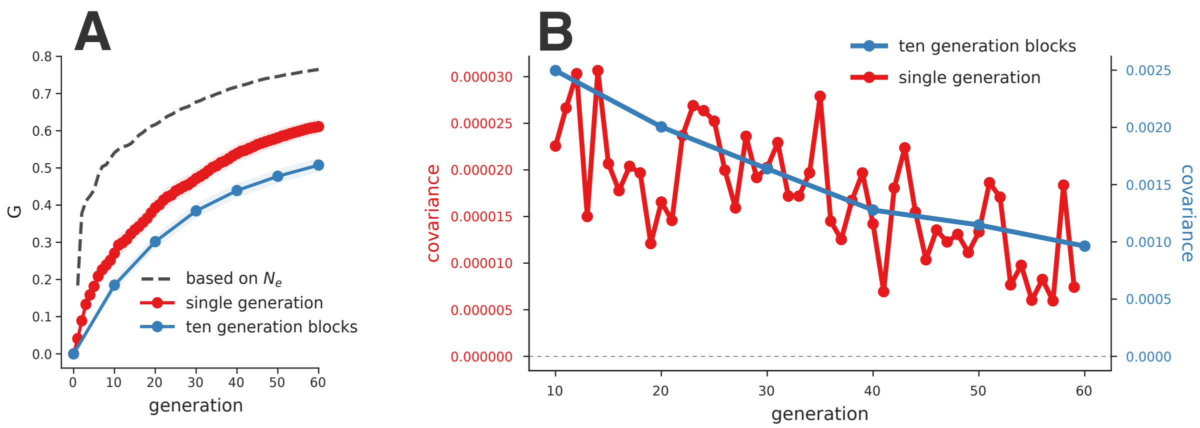

**Figure S14:** A: The $G(t)$ averaged over 50 replicate simulations with $V_A = 0.01$ and $L = 1000$. The blue line shows $G(t)$ calculated over ten generation blocks, similar to the calculation of temporal covariances of the Barghi et al. (2019) study. The red line shows the average $G(t)$ estimates when the population is sampled every generation and all covariances can contribute to the numerator of $G(t)$. The dashed dark gray line indicates the ~~$G(t)'$~~ $G'(t)$ estimate, which uses the known drift effective population size of the simulations. B: The temporal covariances calculated each generation (red line) and on ten generation blocks (blue line) using the same simulation data.

~~estimators, the latter of which includes~~ . In Buffalo and Coop (2019), we define an alternative estimator that includes selection's effects on these variance terms~~).~~ ,

$$G'(t) = 1 - \frac{t\mathbb{E}(p_0(1-p_0))}{2N_e \operatorname{Var}(p_t - p_0)}, \tag{S56}$$

which can be calculated when the drift-effective population size $N_e$ can be estimated (see equation 26 in Buffalo and Coop 2019 for details).

To verify that $G(t)$ estimated every ten generations is indeed a lower bound, we used a simulation procedure similar to the exponential fitness function simulations (described in Supplementary Material Section S8.1), and calculated the temporal covariances and $G(t)$ both each generation, and every ten generations. Unlike the simulations described in S8.1 ~~, we began selection~~ where selection began at $10N + 5$ generations, selection starts at generation $10N$ ~~generations~~here, and used trait $V_A = 0.01$ and targeted $L = 1000$ sites affecting the trait.

First, comparing $G(t)$ when sampling population frequencies every generation versus every ten generations, we confirm that the ten-generation block $G(t)$ is a lower bound of the $G(t)$ trajectory when sampling is every generation (red and blue lines in Supplementary Figure S14A). Furthermore, since we control the population size and reproductive sampling scheme in our simulations at $N =$ 1000 diploids, we know the ~~drift effective~~ drift-effective population size in the absence of selection~~.~~ ~~This~~ , which allows us to estimate ~~$G(t)'$, which is a measure of $G(t)$ that accounts for the linked selection's inflation of the variance in allele frequency change between two generations (equation 26, Buffalo and Coop 2019)~~$G'(t)$. Plugging in the ~~drift effective~~ drift-effective population size $N_e =$ 1000 into the expression for $G'(t)$ and using the $\operatorname{Var}(p_t - p_0)$ calculated for different $t$'s, we see that $G(t)$ calculated every generation does not account for linked selection's inflation of $\operatorname{Var}(\Delta p_t)$ and underestimates the true impact of linked selection as expected (dashed gray line in Supplementary Figure S14A).

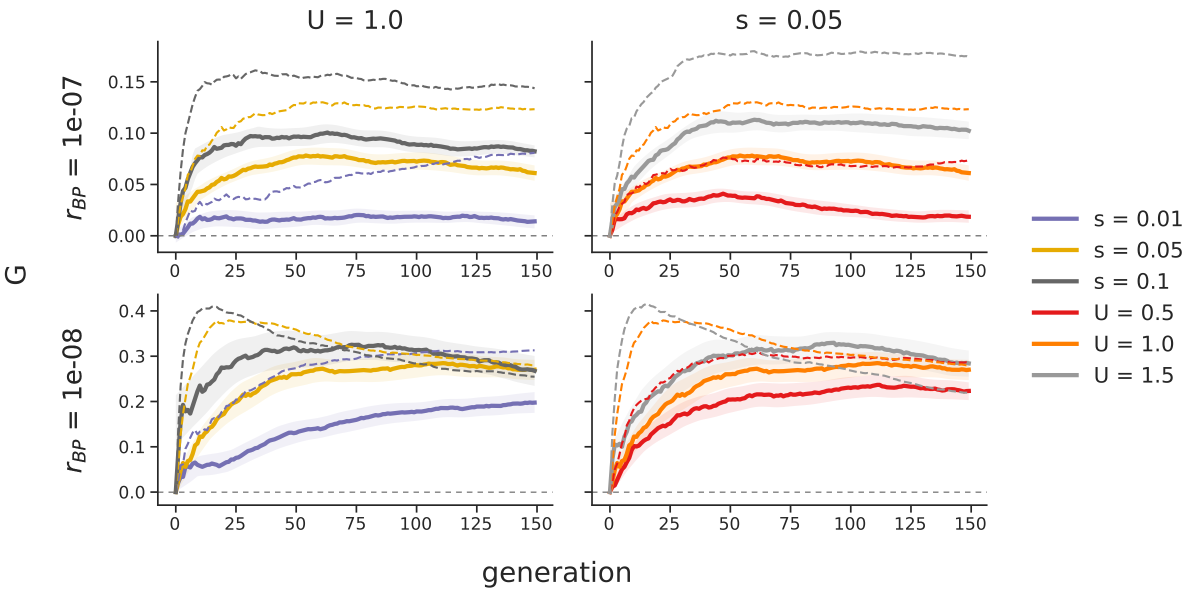

**Figure S15:** The trajectories of $G(t)$ through time under background selection, under different recombination rates ($r_{BP}$, rows), selection coefficients ($s$), and deleterious mutation rates ($U$). The first column ~~sets~~ fixes $U = 1.0$, and $s$ varies, while in the second column $s = 0.05$ is held constant, and $U$ varies. ~~Calculated~~ $G(t)$ is calculated using both including fixed sites (solid lines) and not including fixed sites (dashed lines).

Finally, we found in processing these background selection simulation results that including or excluding sites fixed or lost through time can lead to differences in the estimated $G(t)$ trajectories and temporal covariance. We discuss this extensively in a subsequent section, Supplementary Materials Section S8.7.

#### S8.6 Truncation Selection

We also explored how directional truncation selection generates temporal covariances. In these simulations we select the top 10%, 25%, or 50% of the phenotypic distribution of individuals to form the next generation. We burnin these simulations using a neutral burnin routine, where trait alleles are selectively neutral until directional selection is imposed. More extreme directional truncation selection generated larger initial covariances (Supplementary Material Figure S18B). However, weaker truncation selection generated more sustained positive covariances and so ~~had~~ have a larger long-term impact on the variance in allele frequency change. We again noticed fixation or loss of sites has a strong effect on temporal covariance and $G(t)$ under truncation selection. Here, since only a (potentially small) fraction of individuals contribute to the next generation, sites can fix over very short timescales. Furthermore, the small number of effective breeders contributing to the next generation shrinks $N_e$ considerably, which increases $\text{Var}(p_t - p_0)$, the denominator of $G(t)$. We see both the effect of handling fixed/lost sites differently and the faster rate of drift in Supplementary Materials Figure S18A, where weaker truncation selection actually has higher levels of $G(t)$. Looking just at temporal covariances, we find that stronger truncation selection (e.g. a smaller tail of individuals selected) does lead to greater temporal covariances. Overall, these truncation selection simulations demonstrate the value of considering both absolute measures

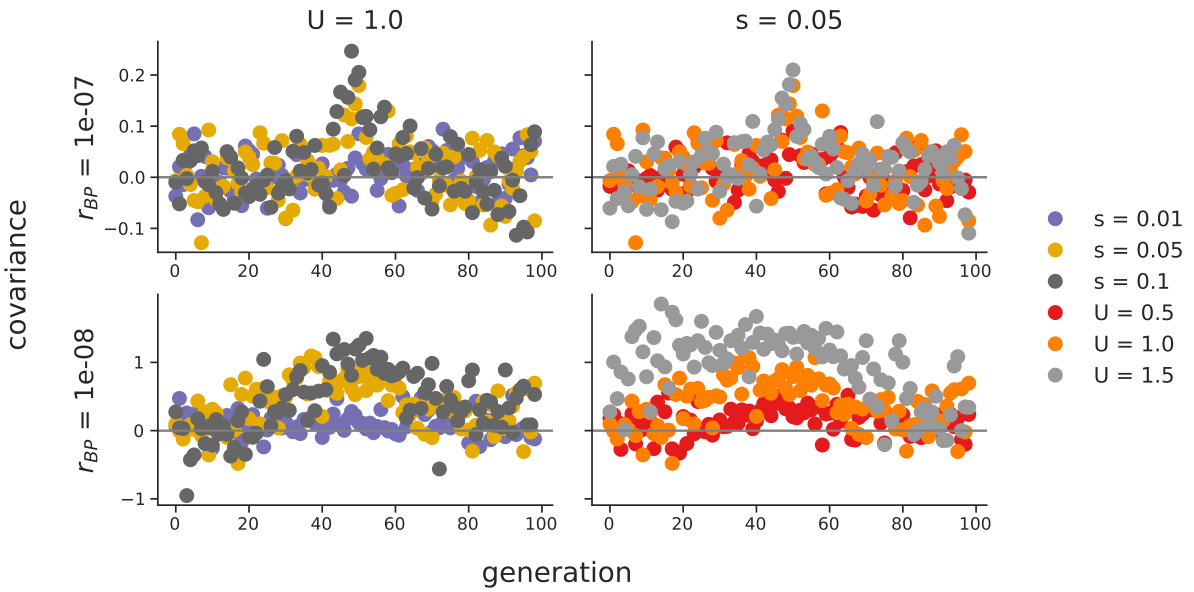

**Figure S16:** The temporal covariances, $\text{Cov}(\Delta p_{50}, \Delta p_t)$ (where $t$ varies along the x-axis) created by background selection, under different recombination rates ($r_{BP}$, rows), selection coefficients ($s$), and deleterious mutation rates ($U$). Unlike directional selection figures, where we choose the reference generation to be the first generation after the onset of selection, here we choose an arbitrary reference generation (generation 50). The symmetry of temporal covariance around the reference generation, is expected, since unlike directional selection the level of additive genetic variance for fitness has hit mutation-selection-drift balance. Note that the first column sets constant $U = 1.0$, and $s$ varies, while the second column sets $s = 0.05$ constant, and varies $U$.

of selection's effect on allele frequency changes, i.e. temporal covariance, as well as measures relative to drift, i.e. the $G(t)$ trajectories. While selecting a smaller tail of individuals is associated with *stronger* selection, it also is leading to a faster rate of drift, which can distort conclusions inferred from considering the $G(t)$ trajectories alone.

##### S8.7 The Effect of Fixations in the Empirical Datasets

In our simulation results, we noticed the temporal covariances and $G(t)$ statistics can differ depending on how allele frequencies of zero or one are handled. Generally, temporal covariances should be calculated on polymorphic sites; once a site has reached fixation or loss, its allele frequency change $\Delta p_t = 0$ and including these sites in the temporal covariance calculation can lead to biases. However, with *sample* allele frequencies, rather than population frequencies, a site with observed frequency zero or one may still be segregating, but by chance not sampled at a timepoint. Here, we discuss the effect of including sites with frequency zero or one, and show our empirical results are not qualitatively different when analyzed excluding fixed sites.

With empirical data calculated on sample allele frequencies, low frequency minor alleles may not be sampled at some timepoints, and excluding these observations (instead of treating it as a trajectory that has a 0 frequency timepoint) biases estimates of quantities such as $N_e$ towards intermediate frequency alleles. Additionally, we tried only dropping fixed or loss sites from the temporal covariance calculations that were at the end or the beginning of a trajectory (e.g., as if

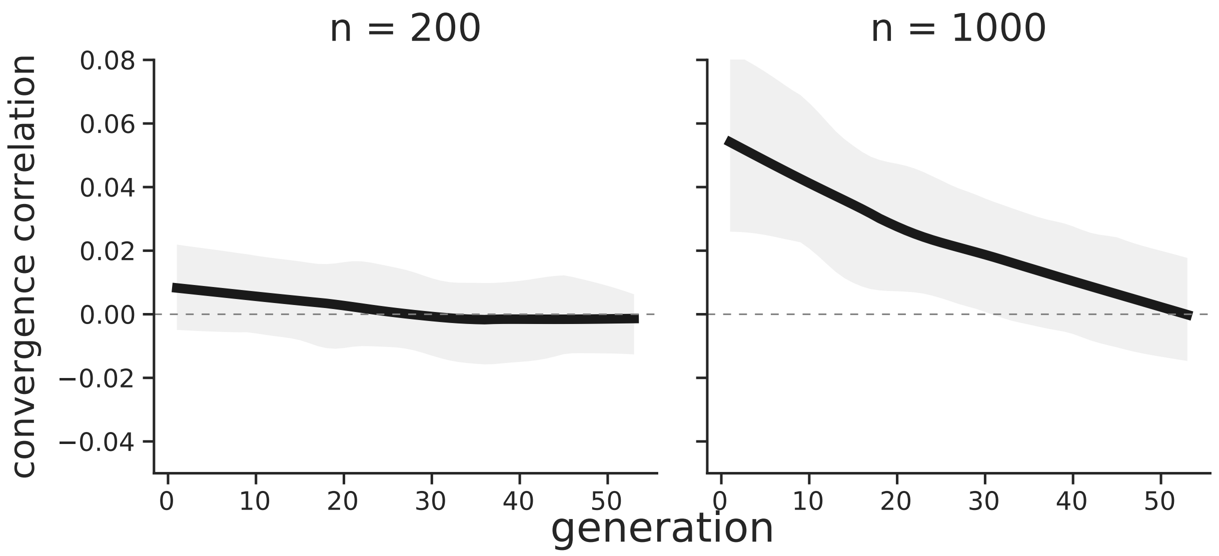

**Figure S17:** The convergence correlation created by background selection through time, since the population split. The replicate population size varies between the two panels. Values are averaged over 30 replicate simulations, while the interval is a 95% confidence interval.

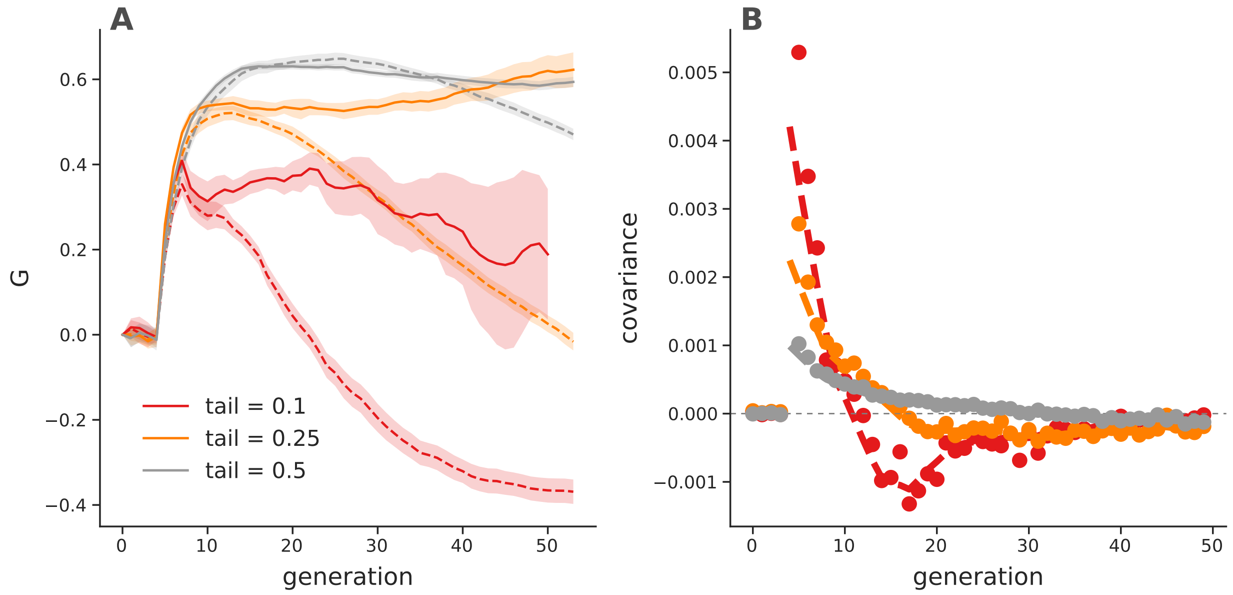

**Figure S18:** $G(t)$ trajectories (A) and temporal covariances (B) from truncation selection simulations for different numbers of individuals selected (line color). Dashed lines indicate $G(t)$ trajectories and temporal covariances calculated *including* fixed sites, while the solid lines exclude fixed sites. All values are averaged over 30 replicate simulations; the lines in the right figure are loess smoothed, while points are averages. The solid lines of the temporal covariances have been excluded in the left figure for clarity, but are similar except they do not become negative.

the site was created by a new mutation or fixed); while this ameliorated some of this bias it did not remove all of it. Overall, we found by trying all these approaches that not removing fixed or lost sites was the best way to deal with sample allele frequencies that could be missing from some timepoints.

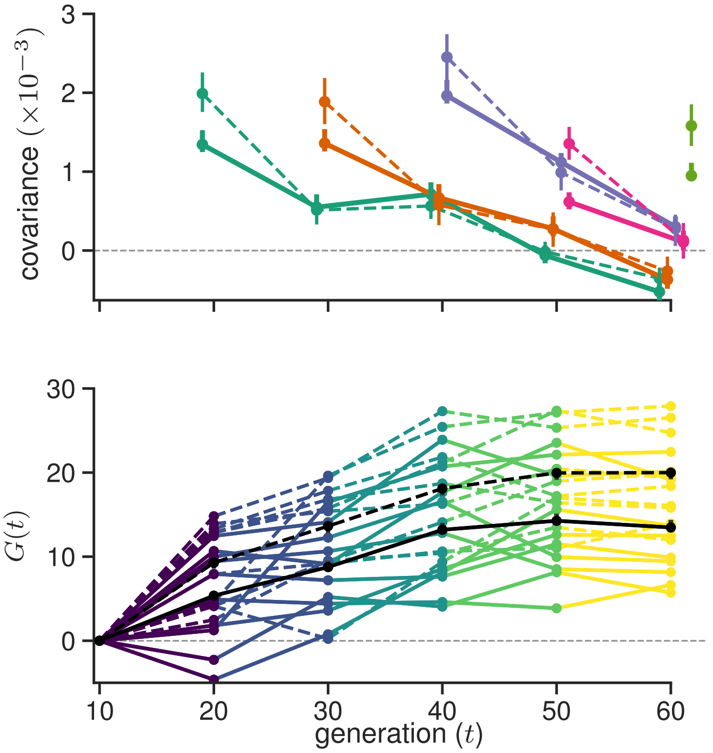

**Figure S19:** The effect of excluding fixed/lost sites in the calculation of the temporal covariances and $G(t)$ trajectories of the Barghi et al. (2019) data. Dashed lines are those including fixed/lost sites (i.e. the original Figure 1), and solid lines are excluding fixed/lost sites.

To ensure that our findings were robust to handling sites with a frequency of zero or one differently, we regenerated Figures 1 and Figure 3 but excluded frequencies of zero or one. Specifically, we wanted to ensure that our finding that temporal covariances were negative at later timepoints was not spuriously caused by the way fixed or lost sites are handled. We see no qualitative difference (Supplementary Material Figures S19 and S20) that emerges when sites with frequency zero or one are excluded.

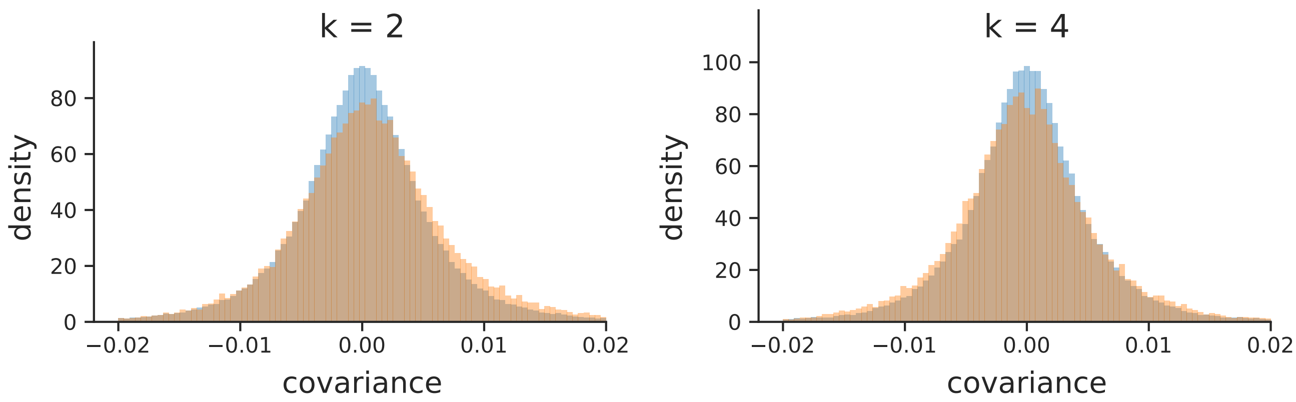

**Figure S20:** A version of Figure 3 (A) and (B) excluding fixed and lost sites. The same qualitative pattern holds as the original figure, which did not exclude fixed and lost sites: there is an enrichment of positive temporal covariances between near timepoints ($k = 2$) in the Barghi et al. (2019) study, and an excess of negative temporal covariances at more distant timepoints ($k = 2$).

### S9 ~~Supplementary~~ Additional Figures

#### S9.1 PCA of Barghi et al. (2019) replicates

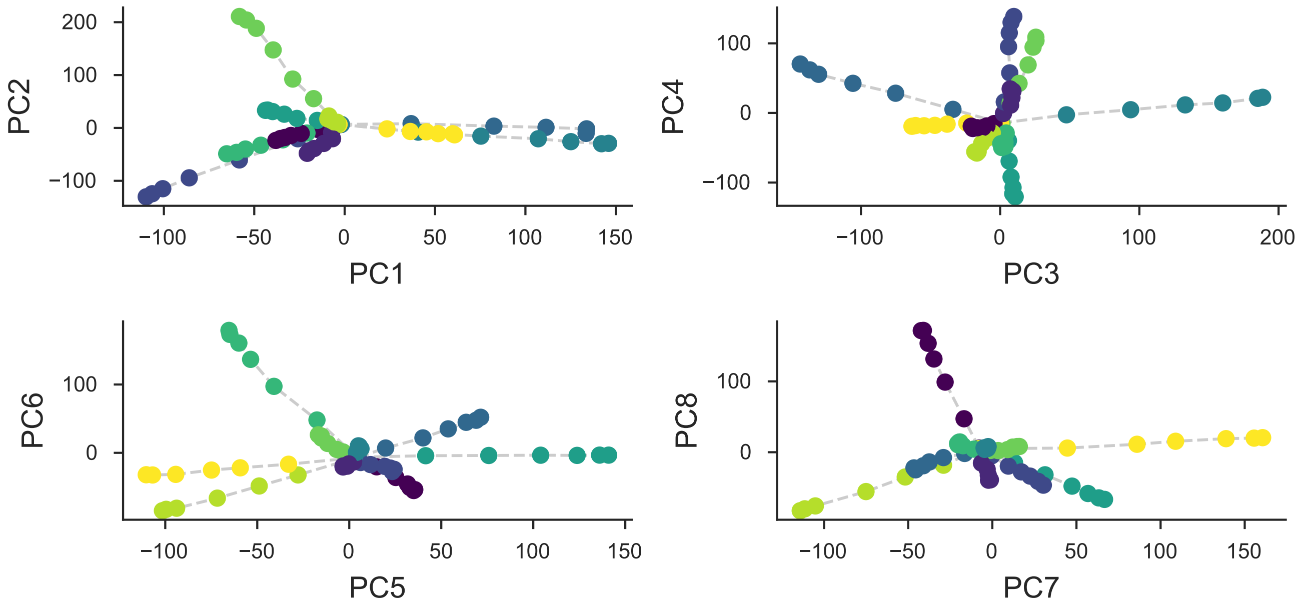

**Figure S21:** A PCA on the centered and standardized population frequencies for each replicate (each color) for all its sequenced timepoints (the connected series of points). All replicates start from the same source population, and thus are overlapping in the center; as each replicate evolves independently it diverges from the other replicates in PCA space.

#### S9.2 Bias Correction for Barghi et al. (2019)

We have investigated the effectiveness of our correction on real data by exploiting the relationship between sampling depth and the magnitude of the variance and covariance biases, and comparing the observed variances and covariances before and after correction. We plot the variance and covariance (between adjacent time intervals) before and after the bias correction against the average sample depth in 100kb genomic windows in Figure S22. Overall, we find the biased-correction procedure removes the relationship between variance and covariance and depth, indicating it is working adequately.

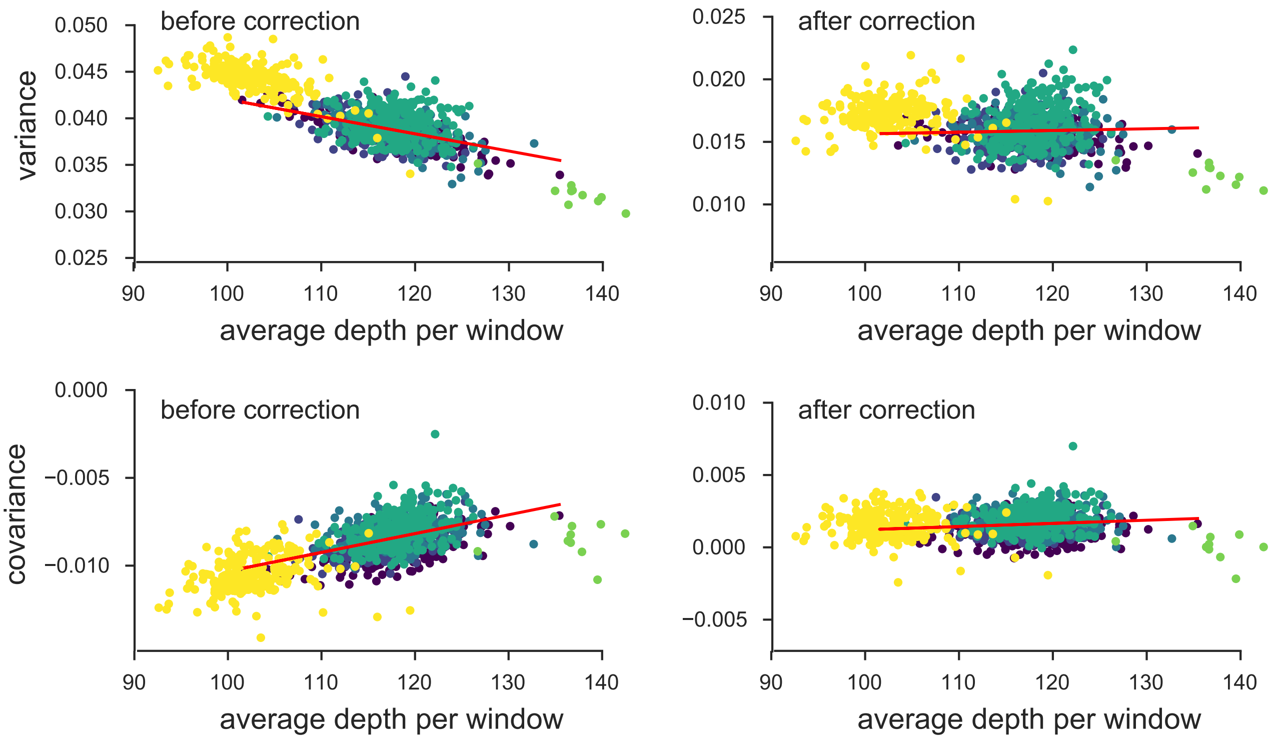

**Figure S22:** The variance and covariances from the Barghi et al. (2019) study, calculated in 100kb genomic windows plotted against average depth in a window before and after bias correction. Each panel has a least-squares estimate between the variance and covariance, and the average depth. Overall, the bias correction corrects sampling bias in both the variance and covariance such that the relationship with depth is constant. Colors indicate the different chromosomes of *D. simulans*; we have excluded the X chromosome (yellow points) and chromosome 4 points (green points to far right) from the regression due to large differences in average coverage.

#### S9.3 Barghi et al. (2019) Temporal Covariances Per Replicate

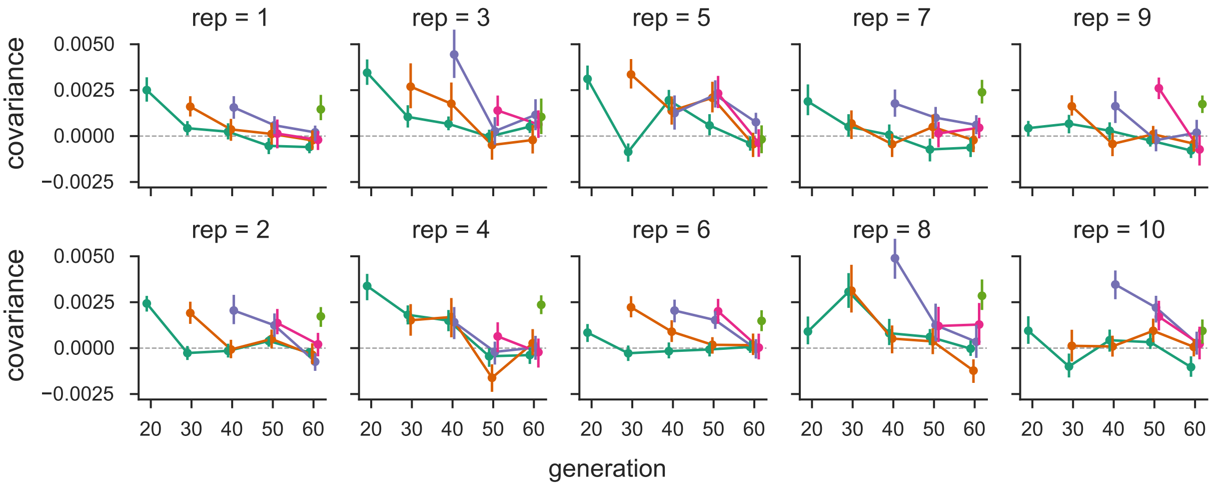

**Figure S23:** The temporal covariances from the Barghi et al. (2019) study, for each replicate individually. As in Figure 1, each line follows the temporal covariances from some initial reference generation through time, which represent the rows of temporal covariance matrix.

| s | t | median | median 95% CI | trimmed mean | trimmed mean 95% CI |
|---|---|---|---|---|---|
| 0 | 10 | 1.629 | $[1.532, 1.738]$ | 1.874 | $[1.777, 1.969]$ |
| 0 | 20 | 0.371 | $[0.276, 0.465]$ | 0.491 | $[0.403, 0.585]$ |
| 0 | 30 | 0.479 | $[0.4, 0.589]$ | 0.516 | $[0.434, 0.602]$ |
| 0 | 40 | 0.059 | $[-0.012, 0.15]$ | 0.027 | $[-0.05, 0.099]$ |
| 0 | 50 | -0.204 | $[-0.271, -0.125]$ | -0.259 | $[-0.329, -0.187]$ |
| 10 | 20 | 1.549 | $[1.427, 1.659]$ | 1.722 | $[1.617, 1.83]$ |
| 10 | 30 | 0.438 | $[0.339, 0.539]$ | 0.506 | $[0.399, 0.609]$ |
| 10 | 40 | 0.233 | $[0.149, 0.328]$ | 0.254 | $[0.159, 0.343]$ |
| 10 | 50 | -0.355 | $[-0.454, -0.289]$ | -0.319 | $[-0.401, -0.237]$ |
| 20 | 30 | 1.981 | $[1.856, 2.095]$ | 2.195 | $[2.084, 2.302]$ |
| 20 | 40 | 0.792 | $[0.698, 0.894]$ | 0.903 | $[0.815, 0.999]$ |
| 20 | 50 | 0.123 | $[0.042, 0.207]$ | 0.221 | $[0.141, 0.309]$ |
| 30 | 40 | 1.296 | $[1.208, 1.425]$ | 1.385 | $[1.287, 1.483]$ |
| 30 | 50 | 0.07 | $[-0.037, 0.183]$ | 0.116 | $[0.023, 0.21]$ |
| 40 | 50 | 1.36 | $[1.271, 1.446]$ | 1.513 | $[1.427, 1.601]$ |

**Table S1:** Table of median of windowed covariance estimates ($\text{Cov}(\Delta p_s, \Delta p_t) \times 100$) between generations $t$ and $s$ and the trimmed mean windowed covariance which excludes the lower and upper 5% windows with the highest covariance.

#### S9.4 Barghi et al. (2019) Trimmed Window Covariances

Here we report median and trimmed mean of the windowed covariances (Supplementary Table S1). We note that the median covariance is also limiting result of a trimmed mean that symmetrically excludes the upper and lower $\alpha$ tails to calculate the trimmed average windowed covariance. As $\alpha$ increases to 0.5, the trimmed covariance converges to the median windowed covariance (by the definition of the median; see Supplementary Figure S24). Thus our genomic temporal covariances are non-zero due to the impact of selection on many genomic windows.

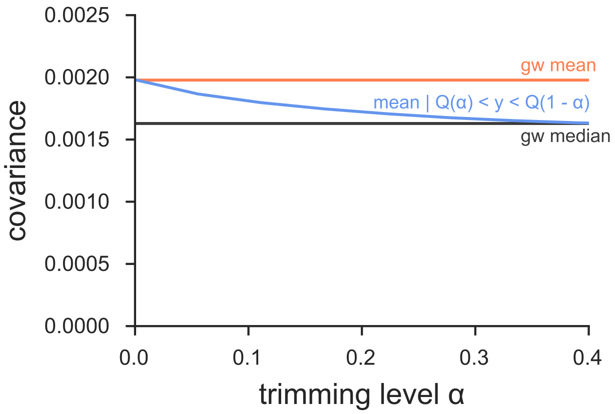

**Figure S24:** The genome-wide covariance ($\text{Cov}(\Delta p_0, \Delta p_{10})$ pooling all replicates) averaged (red line) and the median windowed covariance (blue) for the Barghi et al. (2019) dataset. The trimmed average window covariance, excluding the $\alpha$ lower and upper tails, converges to the median windowed covariance. This indicates that genome-wide covariances are not being overly dominated by a large-effect loci in few windows.

1238 ## S9.5 Barghi et al. (2019) Empirical Null and Windowed Covariance Distributions

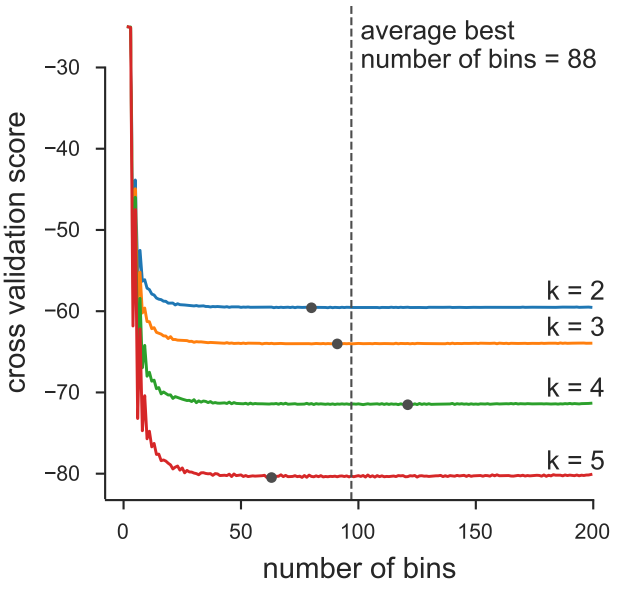

**Figure S25:** We chose number of bins used in the histograms of Figure 3 via an analytic expression for the cross-validation risk, based on the equation 6.16 of (Wasserman 2006, p. 129). Above, we plot the cross-validation risk for various numbers of bins, for each of the four off-diagonals of the temporal covariance matrix that we analyze. Overall, because the number of data points is large, oversmoothing is less of a problem, leading the cross-validation risk to be relatively flat across a large number of bins. Each gray point indicates the minimal risk for a particular off-diagonal, and the dashed line indicates the best average binwidth across off-diagonals.

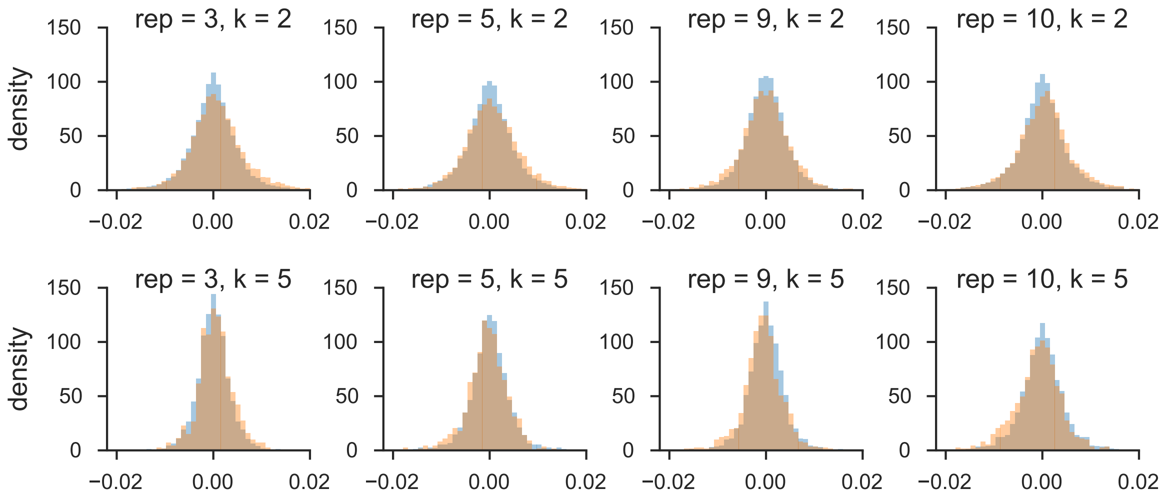

**Figure S26:** The distribution of windowed temporal covariances alongside the empirical neutral null for five randomly sampled replicates (columns), for $k = 2$ (first row) and $k = 5$ (second row). The main figure of the paper pools all replicate window and empirical neutral null covariances; we show here the windowed temporal covariances tend to shift from being positive (a heavier right tail) to become more negative (a heavier left tail) through time within particular replicates.

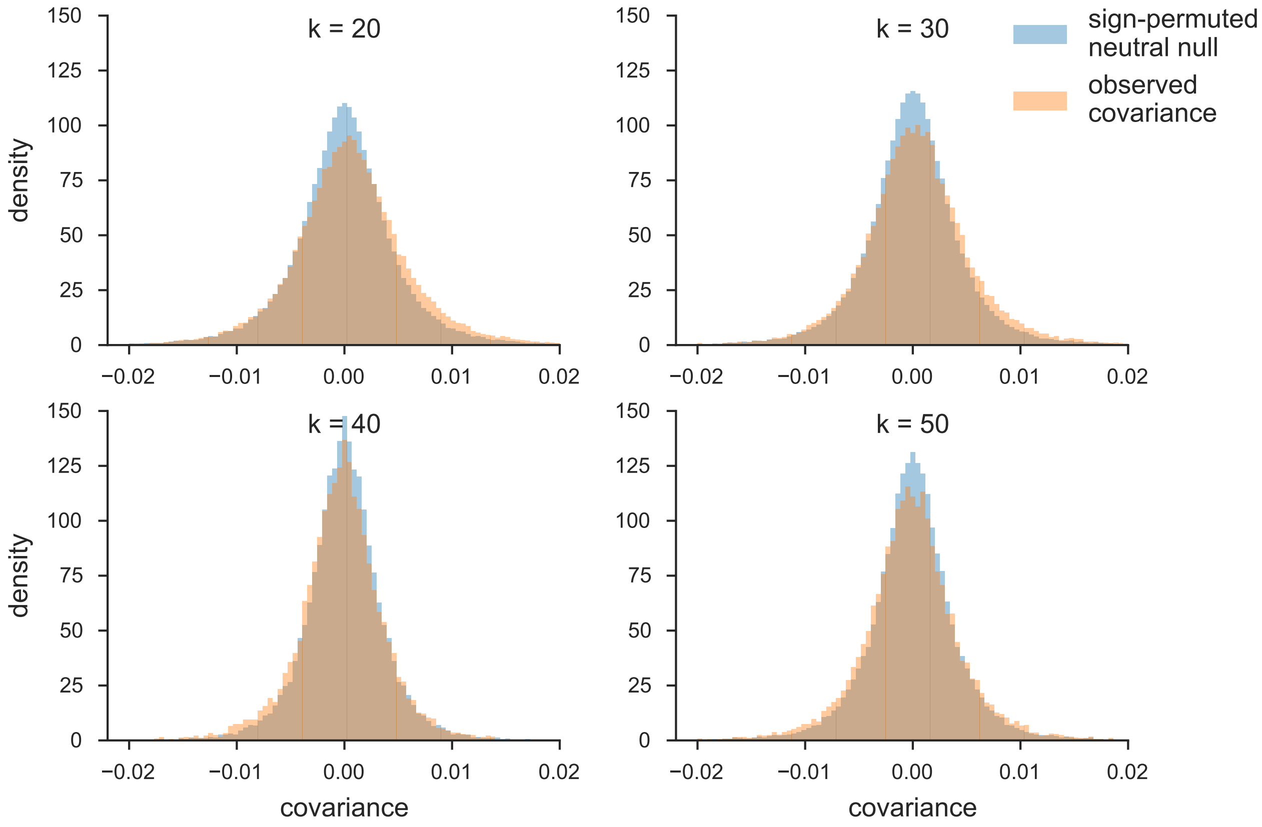

**Figure S27:** The distribution of temporal covariances calculated across 100kb genomic windows from Barghi et al. (2019)'s study (orange) and the block sign permuted empirical neutral null distribution of the windowed covariances (blue). Each panel shows these windowed covariances and the empirical null distribution for covariances $\mathrm{Cov}(\Delta p_t, \Delta p_{t+k})$, $k$ is the number of generations between allele frequency changes.

#### S9.6 Barghi et al. (2019) Tail Probabilities for Windowed Covariances Distributions

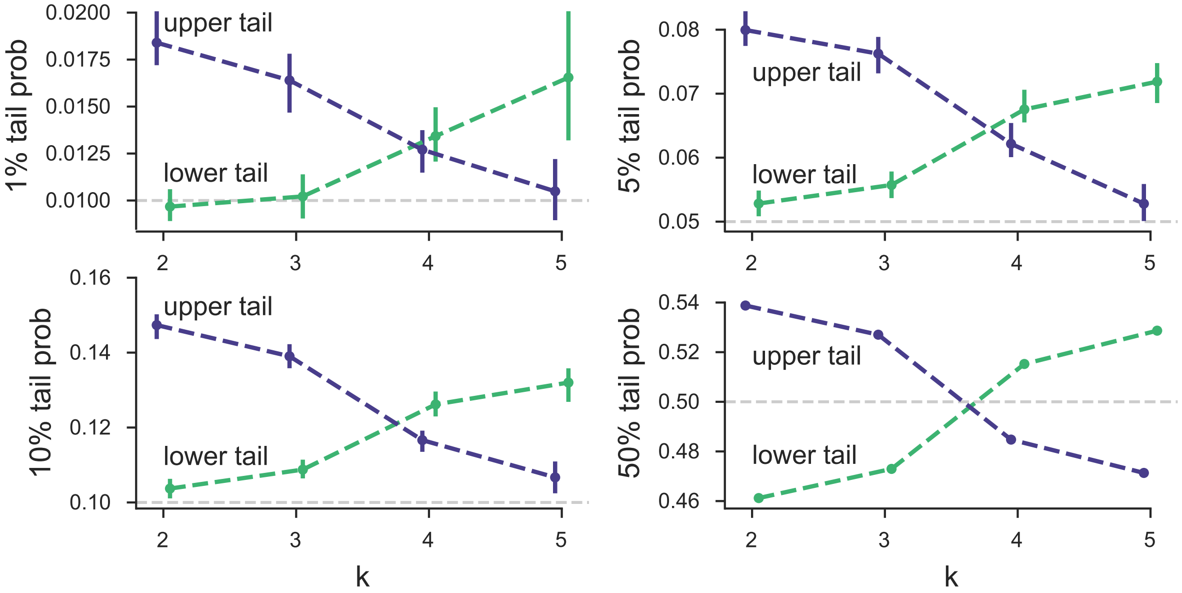

**Figure S28:** Barghi et al. (2019) tail probabilities compared to sign-permuted empirical null distribution for various $\alpha$ levels.

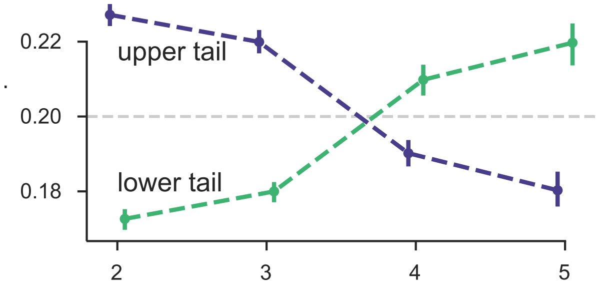

**Figure S29:** The 20% lower and upper tail probabilities for the observed windowed covariances from the Barghi et al. (2019) study, based on sign-permuting at the chromosome level. This permutation empirical null is robust to long-range linkage disequilibrium acting over entire chromosomes (see Supplementary Material section S5).
